# Diagnose Harmonisation (DHARM): An open-source tool for the harmonisation and analysis of batch effects in MRI-derived measures

**DOI:** 10.64898/2026.10.30.763051

**Authors:** Jacob Turnbull, Gaurav Bhalerao, Thomas E. Nichols, Paul McCarthy, Grace Gillis, Jasmine Blane, Brian Zhaoyi Mo, Shaun Warrington, Stamatios N. Sotiropoulos, Renato Besenczi, Fidel Alfaro-Almagro, Stephen M. Smith, Ludovica Griffanti

## Abstract

Technical differences between scanners, acquisition protocols, machine hardware and processing pipelines can introduce systematic variation into MRI-derived measures. Numerous harmonisation methods have been developed to reduce these effects. However, validation of these methods is often performed using different criteria depending on the method itself, the datasets used and the study design. Consequently, determining whether harmonisation is required, selecting an appropriate method and establishing whether or not technical variation has been sufficiently removed without compromising biological signal remain challenging. In this manuscript, we introduce Diagnose Harmonisation (DHARM), an open-source Python toolbox for diagnosing batch effects, applying harmonisation and evaluating harmonisation performance in cross-sectional and repeated-measures settings. DHARM is user-friendly, comes with GUI-based and command-line interfaces, and has graphical reporting. It provides standardised reporting workflows combining complementary measures of batch-associated differences in feature location, scale, empirical distributions, covariance and multivariate structure, together with measures assessing preservation of biological covariate effects and repeated-measures consistency. The framework includes several established harmonisation approaches, but its evaluation workflow is method-agnostic: users can also supply data harmonised using external or newly developed methods and benchmark their outputs using the same common set of diagnostics. We demonstrate DHARM using multisite cross-sectional MRI measures from UK Biobank, Cam-CAN and OxBAC as well as travelling-heads data from the ON-Harmony dataset. Across the cross-sectional datasets, harmonisation substantially reduced additive batch effects, while residual differences in variance and empirical distributions varied between methods. Multivariate diagnostics further demonstrated that reduction of batch-associated structure can coexist with preservation or increased detectability of biologically relevant variation. In the repeated-measures example, harmonisation reduced batch-associated and within-subject variability, while broadly preserving subject-level and biological relationships. DHARM provides a reproducible framework for diagnosing batch effects, applying different harmonisation methods and evaluating harmonisation as a multi-criteria problem rather than relying on a single measure of success. By jointly characterising technical variation and preservation of biologically meaningful information, DHARM can support dataset- and objective-specific selection, benchmarking and validation of harmonisation strategies.

## 1 Introduction

Magnetic resonance imaging (MRI) is a powerful, non-invasive technique for investigating the structure and function of the brain and other organs. However, MRI measurements are strongly influenced by technical factors, including acquisition parameters, scanner hardware, reconstruction methods, and image-processing pipelines. These sources of variation can introduce systematic differences between datasets that are unrelated to the underlying biology. Consequently, MRI studies were historically conducted at a single site, and combining data acquired across scanners or institutions was often considered unreliable.

Advances in image acquisition, the development of standardised protocols, and the increasing use of reproducible image-processing pipelines have made multisite imaging studies more feasible. Open-science platforms such as Pulseq and OpenNeuro have also facilitated the sharing of acquisition protocols and imaging data. As a result, datasets collected at different sites, and sometimes for different original purposes, can increasingly be pooled retrospectively. Such aggregation can substantially increase sample size, statistical power, population diversity, and the overall research potential of existing datasets.

Nevertheless, technical differences between datasets remain. These differences, commonly described as batch, scanner, or site effects, may be present in the images themselves and in subsequently derived imaging phenotypes. Early approaches generally attempted to account for these effects by including categorical site variables in statistical models or by applying regression-based deconfounding procedures (Alfaro-Almagro et al., 2021). At the image level, relatively simple methods such as intensity normalisation and histogram matching were also used to make images acquired under different conditions more comparable. These approaches demonstrated that cross-site comparisons were possible, but they also exposed important limitations. Regression-based methods can effectively remove average differences between sites, particularly when site effects are approximately additive. However, they do not necessarily align the variances, distributional shapes, or covariance structures of different datasets. They may also be unable to distinguish subject-level biological variability from acquisition-dependent measurement noise. Similarly, simple image-level normalisation methods may reduce gross intensity differences without addressing more complex scanner-related effects.

These limitations motivated the development of dedicated statistical harmonisation methods. The most widely applied of these is ComBat, which was originally developed for removing batch effects from high-dimensional genomic data and was subsequently adapted for neuroimaging (Fortin et al., 2017; Johnson et al., 2007). ComBat uses a regression model to estimate and preserve variation associated with biological covariates before applying an empirical Bayes procedure to estimate site-specific differences in the location (mean) and scale (variance) of the data. By adjusting both, ComBat can correct aspects of site-related variation that cannot be addressed through mean-centring or conventional regression alone.

ComBat nevertheless relies on several assumptions about the form of the data and the nature of the batch effects. A growing family of extensions has therefore been developed for settings in which these assumptions may not hold. ComBat-GAM (Pomponio et al., 2020), for example, uses generalised additive models to represent non-linear relationships between imaging features and continuous covariates such as age. CovBat (Chen et al., 2022) extends location-and-scale adjustment by also targeting differences in covariance structure between batches. Reference-based implementations can harmonise datasets relative to a designated reference population, which may be useful when one dataset is regarded as a technical standard or is substantially larger than the others (Turnbull et al., 2026).

Other approaches use image quality metrics (IQMs) to characterise acquisition-related variation. IQMs may be used as proxies for otherwise unobserved technical effects or included alongside site labels to provide additional information, particularly when the number of observations per site is limited (Bhalerao et al., 2026; Gaser et al., 2026; Prevot et al., 2025; Yu et al., 2024; A. H. Zhu et al., 2024). These approaches may be valuable when technical variation exists within sites, when scanner metadata are incomplete, or when a categorical site variable does not adequately describe the relevant sources of measurement heterogeneity.

In parallel with statistical methods applied to imaging-derived features, a broad range of image-level harmonisation techniques has been proposed. These include histogram matching, intensity standardisation, synthesis-based methods, and deep learning approaches that either map images between acquisition domains or learn representations intended to be invariant to scanner and site (Hu et al., 2023; Wen et al., 2023). Such methods may address spatially complex effects that cannot be captured through downstream feature-level adjustment. However, image-based approaches, particularly those based on deep learning, are not yet suitable for every harmonisation problem. They can require large and representative training datasets, substantial computational resources, and careful validation across acquisition settings. Their performance may be sensitive to training-data bias, and models may overfit to the scanners, populations, or contrasts represented during development. They may also alter anatomical or pathological features when the distinction between technical and biological variation is not adequately constrained. These limitations can restrict their scalability, robustness, and generalisability across sites and datasets (Hu et al., 2023; Wada et al., 2024; Wen et al., 2023; Wrobel et al., 2020; J.-Y. Zhu et al., 2017)

The expanding range of harmonisation methods reflects the diversity of problems encountered in multisite MRI research. Methods differ in their assumptions, required inputs, computational demands, and the forms of variation that they are designed to remove. Some operate on images, whereas others operate on derived features. Some require explicit batch labels, while others use measured indicators of image quality or acquisition characteristics. Some are designed to correct differences in means and variances, while others target nonlinear effects, covariance structure, or spatially varying image properties (Beer et al., 2020; Carré et al., 2022; Gaser et al., 2026; Pomponio et al., 2020; Prevot et al., 2025). These differences suggest that harmonisation performance is likely to depend on the characteristics of the data and the specific sources of variation present, motivating systematic evaluation across methods and acquisition settings. Characterising the type and amount of bias present in a given scenario is the first critical step in selecting the most suitable harmonisation approach.

Selecting the most appropriate method is also particularly challenging because of the way harmonisation approaches have historically been evaluated. Individual methods are frequently assessed using different datasets, simulation settings, outcome measures, and downstream tasks. Performance reported in one study may therefore not be directly comparable with performance reported elsewhere. A method that appears effective on a large, balanced dataset with pronounced site effects may not behave similarly in a small or highly imbalanced cohort, in the presence of biological confounding, or when the technical differences between sites are subtle (Bayer et al., 2022). A set of standardised metrics to capture harmonisation performance would enable selection of the method that is best suited to remove the bias identified in the data.

There is also no widely accepted framework for determining whether a harmonisation method has adequately removed technical variation while preserving meaningful biological signal. Existing studies commonly use some combination of uni- and multivariate tests such as effects on the mean; measures of residual site association through correlations or linear modelling; visual inspection of the data distributions; site-classification accuracy; or downstream predictive performance on biological variables (Fortin et al., 2017; Gardner et al., 2025; Hu et al., 2023). Each of these approaches captures a different aspect of harmonisation efficacy, but none provides a complete assessment in isolation. For example, reducing the ability of a classifier to predict a site does not necessarily demonstrate that all relevant batch effects have been removed, while aligning marginal feature distributions does not ensure that multivariate relationships or covariate associations have been preserved. Conversely, a method may appear to remove site effects successfully while also attenuating genuine biological differences that are unevenly distributed across sites.

Robust evaluation therefore requires several complementary questions to be considered. Firstly, is harmonisation necessary, and what are the differences (e.g., feature location, scale, distribution, covariance) between batches? If harmonisation is necessary, what are the main differences, and are they improved through the application of a given method? Have known covariate effects and biologically meaningful relationships been preserved? The answers may not be in full agreement across metrics, but the relative importance of each criterion will depend on the intended use of the harmonised data.

In this manuscript, we introduce Diagnose Harmonisation (DHARM), a software package designed to support the diagnosis of batch effects in a dataset and the systematic implementation and evaluation of commonly used harmonisation approaches for MRI-derived measures. The package brings together multiple harmonisation procedures within a consistent workflow and provides a broad set of complementary metrics and visualisations for assessing their effects. These assessments are designed to characterise initial batch-related variation in the data, as well as the extent to which it has been reduced after harmonisation, and the degree to which covariate effects, biological relationships, and other meaningful properties of the data have been preserved.

Rather than assuming that one method will be universally optimal, the package is intended to support method selection by allowing users to evaluate performance in relation to their dataset, study design, and scientific objective. In doing so, it provides a practical foundation for more reproducible, transparent, and evidence-based harmonisation workflows. Importantly, the diagnostic framework is not restricted to the harmonisation methods implemented within DHARM. Users can provide outputs generated by external or newly developed harmonisation approaches and evaluate them using the same reporting workflow. DHARM can also then serve as a benchmarking framework for method development, providing a consistent set of metrics for determining which forms of technical variation have been reduced and whether biologically meaningful variation has been preserved. This enables newly proposed methods to be evaluated alongside established approaches using common diagnostics rather than method-specific definitions of successful harmonisation. By providing pre-built reporting pipelines for different harmonisation scenarios (i.e., cross-sectional and repeated-measures), DHARM enables researchers to compare methods more transparently and to identify trade-offs that may be missed when relying on a single performance measure.

## 2 Methods

### 2.1 Overall workflow

The tool is designed to provide quantitative measures to diagnose the magnitude and type of batch effect present in the data, facilitate the application of harmonisation methods, and assess harmonisation effectiveness (Figure 1). We provide sets of measures as organised workflows for two scenarios: cross-sectional, where each observation is treated as unique, and repeated-measures, where repeated observations need to be taken into account. Additionally, we provide an array of statistical harmonisation methods within the tool, including linear modelling and removal of batch while preserving covariate effects; a modular form of ComBat which combines the functionality of many of its variants into one function to allow for easier use of the ComBat extensions (e.g., non-linear modelling of covariate effects, reference batch, and harmonisation of covariance) when harmonising data; and IQM-based harmonisation using linear mixed modelling (Bhalerao et al., 2026; Pomponio et al., 2020). However, harmonisation does not need to be performed within DHARM. Users may instead provide data harmonised using an external implementation or a newly developed method and pass these outputs directly to the diagnostic or comparison reports. Consequently, the harmonisation functions provided within DHARM represent convenient implementations rather than a requirement of the evaluation workflow. Multiple externally harmonised datasets can be supplied to the comparison report, allowing new methods to be benchmarked against existing approaches using an identical set of diagnostic criteria. The overall flowchart shown in Figure 1 can be used in either repeated-measures or cross-sectional harmonisation scenarios.

The cross-sectional and repeated-measures workflows answer the same broad diagnostic questions: what are the biological and technical batch effects, and are batch effects removed and biological effects preserved after harmonisation? In repeated-measures data, these questions must be evaluated after accounting for the non-independence of repeated observations from the same participant. The repeated-measures workflow, therefore, retains several cross-sectional diagnostics but replaces independence-based comparisons with repeated-measures models where appropriate, while adding measures of within-subject stability and temporal consistency for test-retest settings.

**Figure 1:**
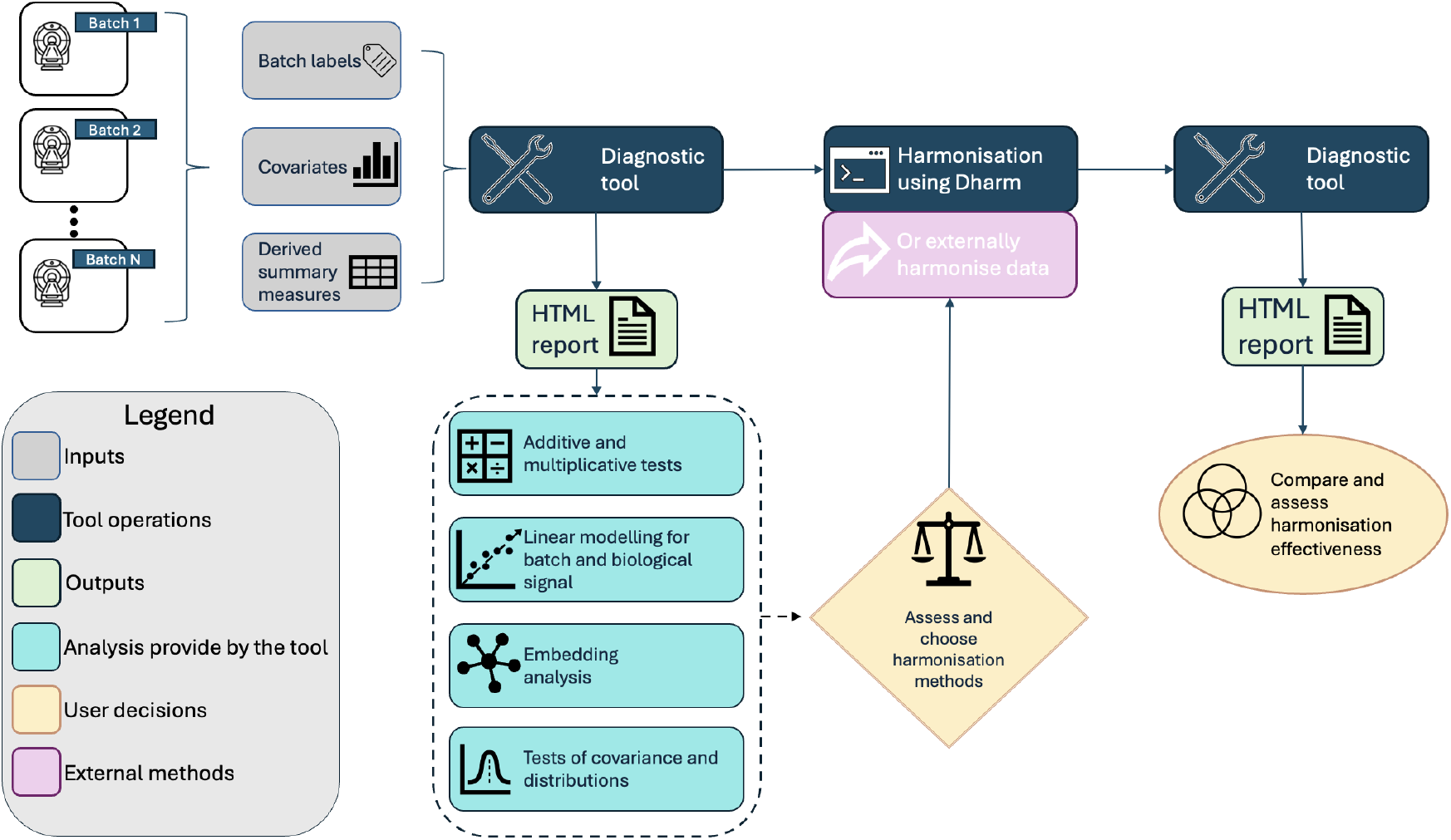
A flowchart showing DHARM workflow for diagnosing batch effects and evaluating harmonisation. Following initial diagnosis, users may apply a harmonisation method implemented within DHARM or supply data harmonised using an external method. Raw and harmonised datasets can then be assessed using the same diagnostic framework.

### 2.2 Using the tool

The current release of the tool can be used by downloading the Diagnose Harmonisation Python package from PyPI (“pip install DiagnoseHarmonisation”) and the development version can be installed similarly from GitHub (“pip install git+https://github.com/JakeTurnbull/HarmonisationDiagnostics”). The main functionality of the diagnostic side of the tool comes from using the automatic reporting functions. These functions produce a structured HTML file with the key results displayed as static figures and as summary metrics, as well as brief suggestions for harmonisation method selection generated automatically. Additionally, these reports will optionally save the results of each test as separate CSV files and figures for later use. The reporting tool can be called in one of three ways: Firstly, within a Python script using the DiagnosticReport module:

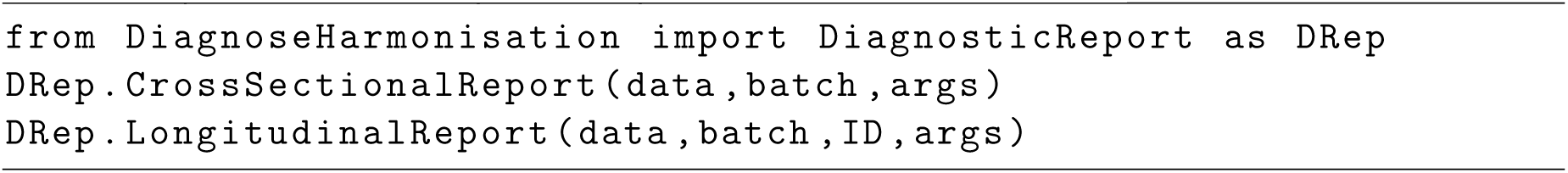

Secondly, from the command line by providing the location of relevant CSV files containing the data, batch and covariate information. Here, the batch label can be passed directly as a separate CSV file or included as a column in the covariates file under the header ‘batch’, ‘site’ or ‘scanner’:

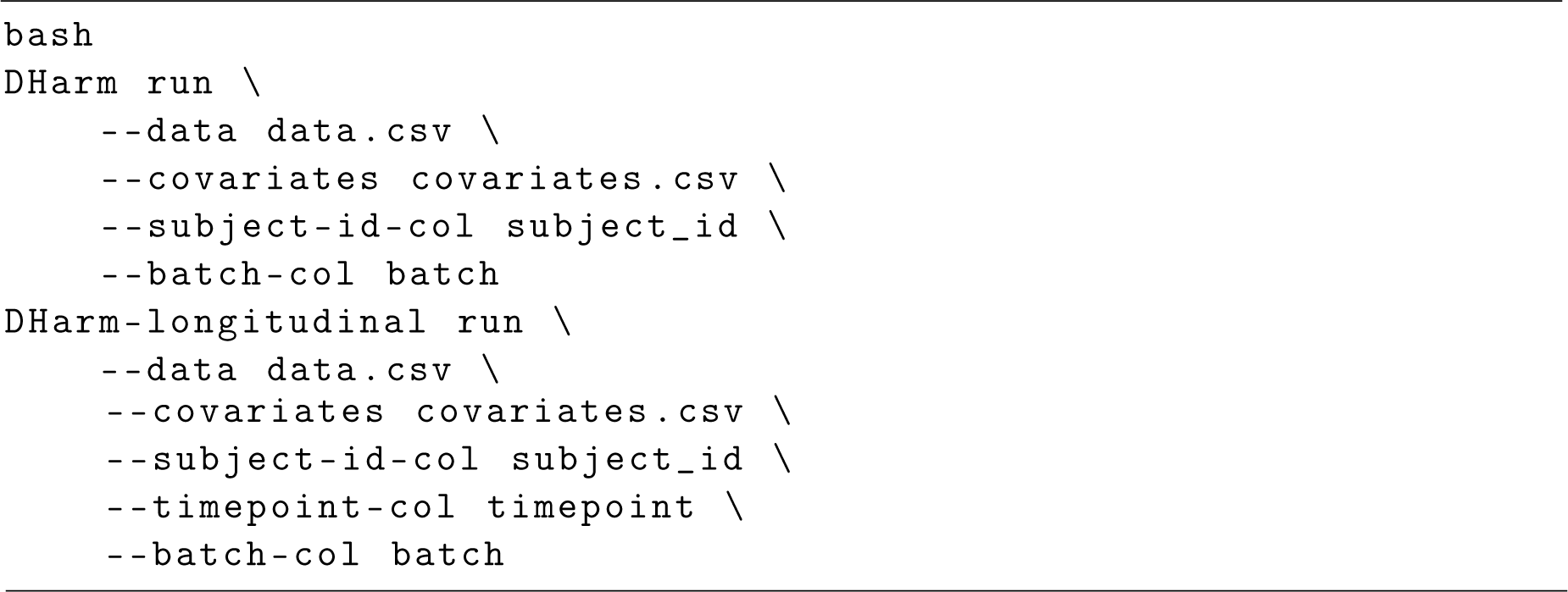

And finally, through the use of an interactive GUI which can be accessed through the terminal. This version allows columns from the called CSV file to be manually selected and inspected prior to running the reporting tool. Both the command-line-only and GUI versions have automatic detection of the batch columns built in, but also allow the name of the column to be specified. In each case, a subject ID column should also be provided. The GUI is shown in Figure 2 and can be launched through the terminal as:

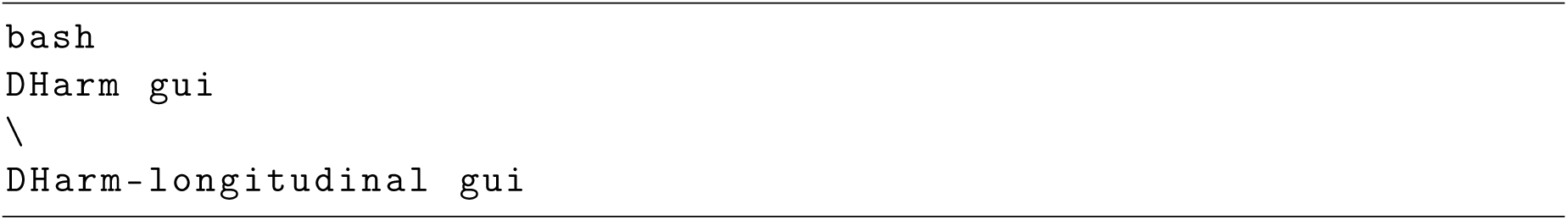

**Figure 2:**
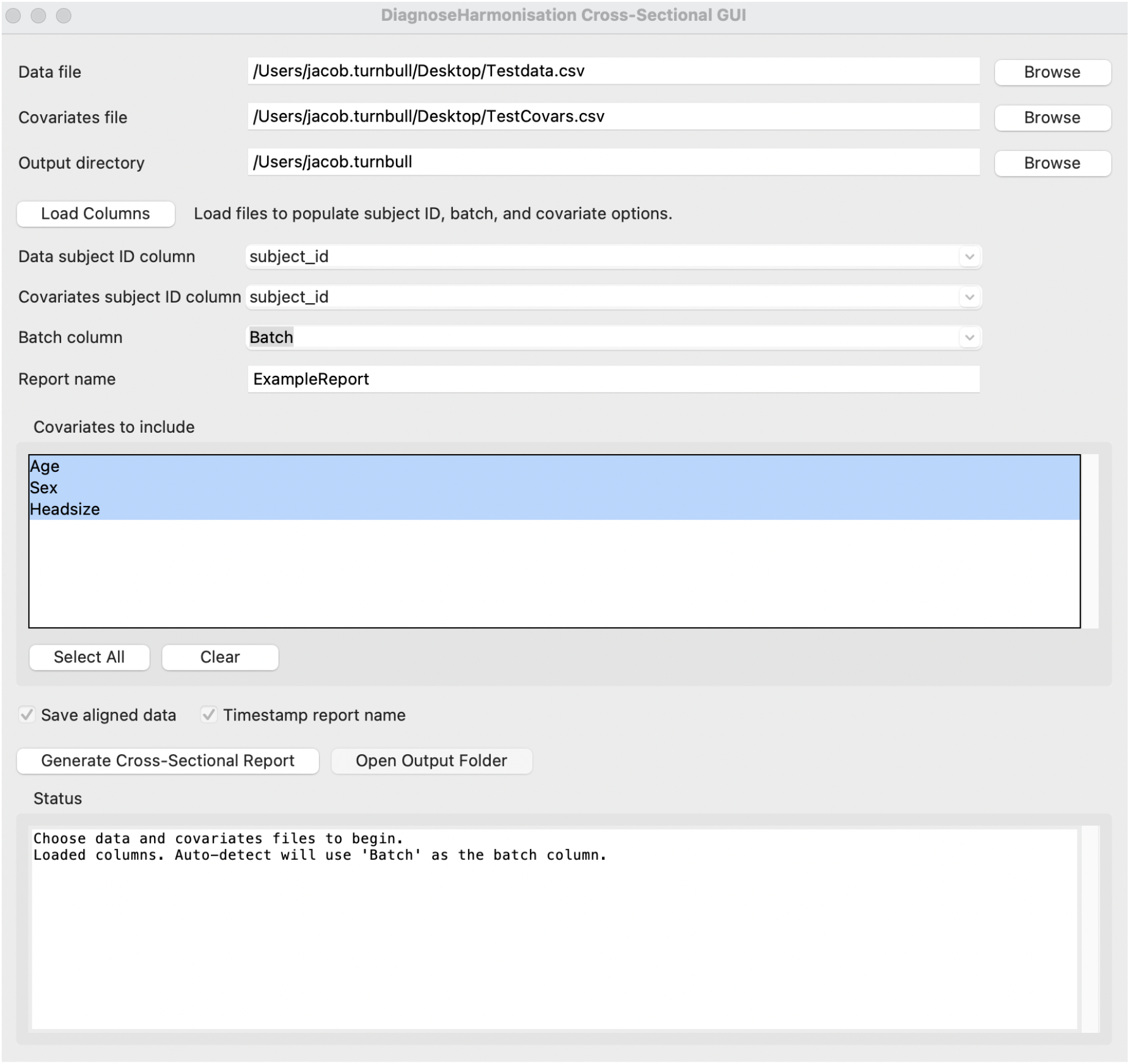
Example GUI called from the terminal with dummy data as the input CSV files. The desired covariates can be selected from the list shown, and the CSV files are matched through a “subject_id” column which is autodetected or selected manually by the user in the case of unmatched headers. The batch column is expected to be provided in the covariate CSV file.

After initial assessment, users can use any harmonisation methods available in the tool by loading the module containing the harmonisation methods within a Python script. The harmonisation methods currently provided in the tool are described in detail within the supplementary materials, and a summary of each, as well as their original publications, repository location and the major differences between the original implementation and the one provided by DHARM, are described briefly in Table 1. Depending on the method chosen, there may be slight differences between the implementation used in DHARM and the original implementation. This may be due to the original language being different, different libraries being used, or different dependencies.

**Table 1:**
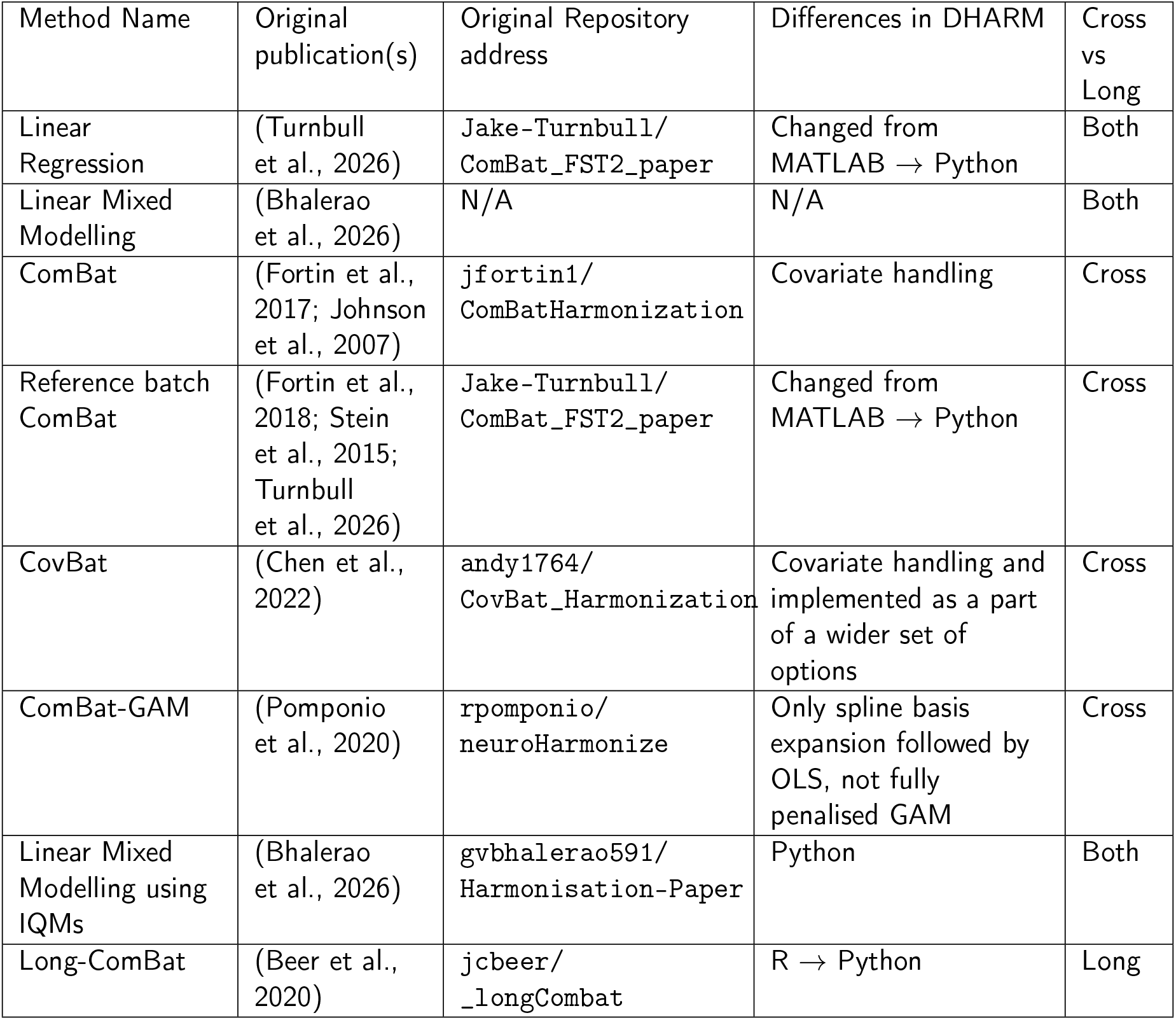
Harmonisation methods included in the DHARM package. Details include original publication(s), the original repository locations, the major changes between the original implementation and the one provided in DHARM and whether the method is repeated-measures or cross-sectional. A full description of each method, how to call them from within the package and the differences between the original implementations are provided in the supplementary materials and within the online DHARM documentation.

Like the reporting tool, the harmonisation methods will require the data array and a categorical batch variable as the minimum inputs, with the other arguments being additional biological covariates and different configurations and settings for a given method, described in detail within the documentation. Support for a terminal and GUI-based harmonisation may be provided in later updates to the tool.

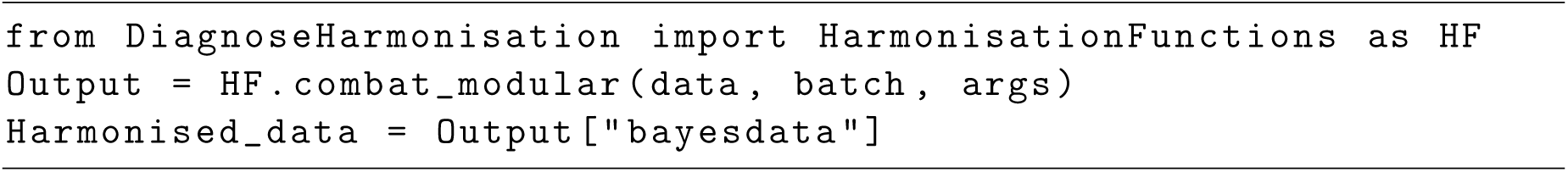

The user may then use the reporting tool on the harmonised dataset and compare the harmonised data report to that of the raw dataset to see how well a given method has done, or directly compare one or more harmonised datasets using the comparison reporting tool which directly compares each method, providing summary plots and rankings of how well each method did by a specific metric and type of measure.

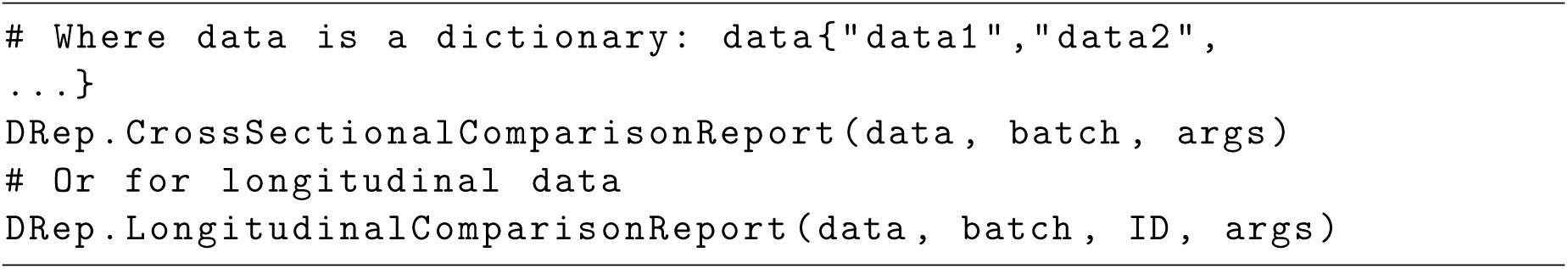

#### 2.2.1 Data preparation and diagnostic context

The tool takes three main inputs: the data matrix of derived features, a vector denoting the batch that the participant belongs to and an optional covariate matrix which can contain variables of interest or confounding variables that the user wants to co-model with the batch variable.

Let 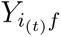 denote an *n × p* matrix of neuroimaging measures from any modality, where rows (*i*) correspond to participants, columns (*f* ) correspond to derived features, and *t* represents the observations’ time point. For cross-sectional data, each subject contributes a single observation, and the t index can be omitted. For repeated-measures (e.g., longitudinal, test-retest), subjects may contribute two or more observations, and as such, a subject identifier (ID) as well as a vector identifying the time point of each observation are additionally required as separate inputs. Each observation *i* must be given a batch label, which can relate to technical differences between the sources of the data, such as scanner, site, vendor, field strength, acquisition protocol or processing pipeline. This is given to the tool as a vector of numerical or string identifiers of the same number of rows as the provided data matrix 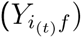.

Finally, while not required for the tool to run, we recommend including an additional matrix of covariates (*C_ic_*), where (*c*) indexes measured sources of variation that should be modelled separately from the batch variable. Covariates are not restricted to effects that should be preserved during harmonisation: they may represent variation that is biologically meaningful and should be retained (e.g., age or sex) or variation that is considered nuisance and may ultimately be removed or controlled for (e.g., signal-to-noise ratio or other measures of data quality). In either case, including these variables allows variation attributable to the measured covariates to be distinguished from variation attributable specifically to batch. Depending on the covariate type (continuous, categorical, or binary), a covariate may correspond to one or multiple columns of (*C*). Not including relevant covariates may lead some methods to attribute covariate-associated variation to batch, particularly when covariates are imbalanced across batches, resulting in an overestimation of batch effects and making it more difficult to assess whether harmonisation appropriately preserves biological information.

The general forms of the covariate-adjusted data is given by Equation 1b and Equation 1d and is used frequently throughout this manuscript. The full equations and descriptions for all methods of analysis are available in the supplementary materials. The unadjusted distributions represent the total observed variation in the imaging features and therefore reflect the combined influence of technical batch effects, biological covariates, covariate imbalance, and residual noise. The covariate-adjusted distributions are designed to assess whether systematic batch differences remain after accounting for measured covariates. Because batch and covariates may be partially collinear in multi-site neuroimaging studies, batch is included in the model used to estimate covariate effects; however, here, the fitted batch contribution is retained rather than removed.

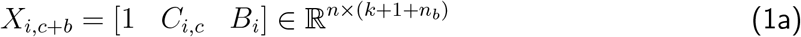

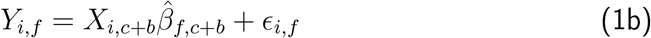

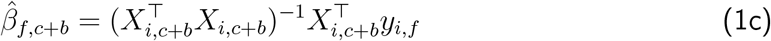

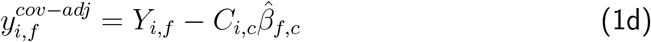

### 2.3 Statistical inference and multiple comparisons

Where diagnostic measures involve statistical hypothesis testing across multiple imaging features or batch comparisons, DHARM reports both uncorrected p-values and p-values adjusted using the Benjamini–Hochberg false discovery rate (FDR) procedure. Correction is performed separately within each diagnostic family rather than across all outputs generated by the report. For omnibus feature-wise tests, FDR correction is performed across features. For diagnostics involving pairwise comparisons between batches, correction is performed across the set of feature-by-batch comparisons. Statistical significance in summary visualisations is based on FDR-adjusted *p_F DR_ <* 0.05 unless otherwise specified. Descriptive statistics and effect-size measures, including Cohen’s *d*, Mahalanobis distance, variance ratios, *R*^2^, ICC and covariance distances, are reported in addition to statistical significance.

### 2.4 Global visual screening of batch differences

#### Visualising distributions

For the first visualisation, each feature is transformed to a robust standardised scale using the Median Absolute Deviation (MAD). This allows measurements with different units and dynamic ranges to be pooled into a common visual summary, while limiting the influence of outlying observations. If the user provides a covariate matrix, we generate two plots, one for covariate-unadjusted and one for covariate-adjusted MAD distributions. Comparing the two visualisations helps determine whether apparent batch differences are largely explained by covariate imbalance or instead persist as residual technical structure.

For repeated-measures data, covariate adjustment additionally accounts for within-subject dependence using a mixed-effects model, with batch and covariates included as fixed effects and subject included as a random intercept. Covariate contributions and the estimated subject-specific random intercept are removed while retaining batch-associated variation. Covariate- and subject-unadjusted and adjusted MAD distributions are provided separately, allowing apparent batch differences to be assessed before and after accounting for measured covariates and subject-specific differences.

#### Covariate imbalances and potential confounding with batch

In addition to visualising the covariate-adjusted and -unadjusted distributions, the tool evaluates the extent to which covariates are associated with batch membership. Differences in the distribution of biological covariates between batches can make biological and technical differences difficult to distinguish, and may lead to apparent batch effects that are partly attributable to differences in the sampled populations both before and after harmonisation.

For each continuous covariate, batch is treated as a categorical predictor in a linear model, and the coefficient of determination (*R*^2^) is used to quantify the proportion of covariate variance associated with batch membership. For categorical covariates, association with batch is quantified using Cramér’s *V* . Both measures range from 0 to 1, with larger values indicating a stronger association between the covariate and batch. These measures should therefore be interpreted as indicators of potential confounding or covariate imbalance, rather than evidence that batch and biological effects are necessarily confounded within the imaging measures themselves.

The report additionally visualises the distribution of each covariate across batches using box plots for continuous covariates and bar charts showing the relative proportion of a categorical covariate by batch membership, allowing the direction of any observed imbalance to be inspected directly. These diagnostics provide context for the subsequent batch-effect analyses and are particularly important when interpreting whether apparent differences between batches are likely to represent technical variation, underlying population differences, or a combination of both.

### 2.5 Analysis on the effect on the mean

Effects on the mean, often referred to as ‘location effects’, occur when measurements from one batch are systematically shifted relative to another. In structural neuroimaging methods, for example, this could appear as a consistent decrease in one batch’s regional volumes or thickness. However, it is also important to test whether these differences could be driven by underlying biology. A summary of the methods used can be seen in Table 2

**Table 2:**
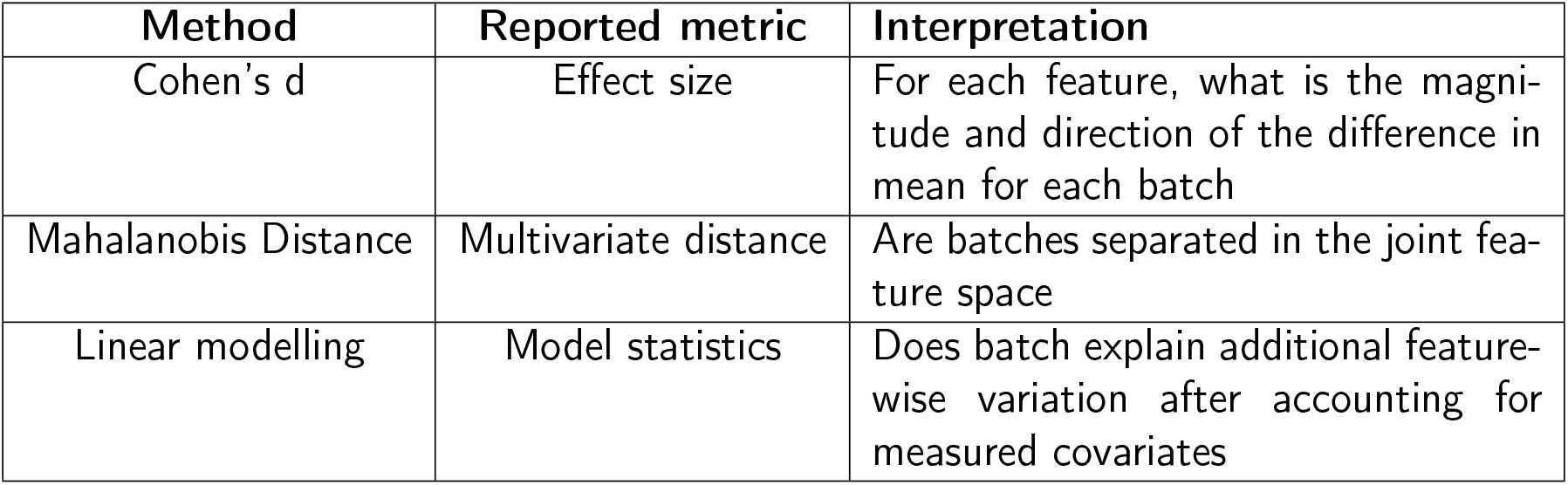
Three methods for analysing the difference in the mean attributable to batch.

| **Method** | **Reported metric** | **Interpretation** |
|---|---|---|
| Cohen’s d | Effect size | For each feature, what is the magni-tude and direction of the difference in mean for each batch |
| Mahalanobis Distance | Multivariate distance | Are batches separated in the joint fea-ture space |
| Linear modelling | Model statistics | Does batch explain additional feature-wise variation after accounting for measured covariates |

#### 2.5.1 Feature-wise mean differences: Cohen’s d

Cohen’s d is used to quantify standardised differences in the mean between batches for each derived imaging measure. This provides an interpretable feature-wise estimate of additive batch effects, expressed in units of pooled standard deviation. Unlike significance testing, the use of effect sizes allows batch differences to be assessed in terms of magnitude rather than sample-size-dependent statistical evidence. Additionally, as different batches are likely to have different covariate distributions (e.g., different ages between batches of data), we first transform the data to be covariate-adjusted, as shown in Equation 1d.

For repeated-measures data, conventional Cohen’s d calculated across observations would treat repeat scans from the same participant as independent and may underestimate uncertainty; as such, we do not use it here and extend the method described in section 2.5.3 for univariate additive batch effect analysis.

#### 2.5.2 Multivariate mean differences: Mahalanobis distance

Neuroimaging-derived measures are often correlated across features. Therefore, small feature-wise shifts that appear modest individually may combine to produce substantial (or at least statistically significant) multivariate separation between batches. Mahalanobis distance captures this joint displacement by scaling differences in mean vectors by the covariance structure of the data. It provides a global measure of batch separation in the joint feature space, complementing univariate effect-size estimates from Cohen’s d. Like Cohen’s d, we also calculate Mahalanobis distance on the covariate-adjusted data, showing the Mahalanobis distance between batches, as well as between each batch and the global centroid of the data for both the adjusted and unadjusted measures.

For the repeated-measures workflow, the Mahalanobis distance framework is adapted to account for the longitudinal structure of the data, with the specific implementation determined by the within-subject batch configuration. When batch assignment remains constant within subjects, Mahalanobis distances are calculated using within-subject changes in feature values between consecutive time points. This approach evaluates batch-associated multivariate separation based on longitudinal changes within individuals, thereby reducing the influence of stable between-subject heterogeneity. In contrast, when batch assignment varies within subjects, each observation is subject-centred by subtracting the subject-specific mean for each feature prior to calculating Mahalanobis distances between batches. This transformation removes subject-specific mean differences and enables batch separation to be evaluated relative to each individual’s average multivariate feature profile. Mahalanobis distance is bounded below by zero but has no fixed upper bound. A distance of zero indicates coincident batch centroids in the covariance-adjusted multivariate space, with progressively larger values indicating greater separation; consequently, no universal threshold defines acceptable batch separation. Lower Mahalanobis distances following harmonisation indicate reduced covariance-adjusted multivariate separation between batches, but do not imply the absence of other forms of batch-related variation. Mahalanobis distance should therefore be interpreted as a measure of multivariate batch separation and considered alongside complementary diagnostics assessing other batch effects and the preservation of biological and longitudinal variation.

#### 2.5.3 Model fit comparison using fixed and mixed-effects models

##### Linear modelling in the cross-sectional tool

The cross-sectional reporting tool fits three complementary statistical models in order to assess the contribution of batch and covariates to the overall variance of each feature: (i) A covariate-only fixed-effects model; (ii) A covariate and batch fixed-effects model; (iii) A linear mixed-effects model with batch as the random grouping variable. Respectively, the models are used to calculate the variance explained by measured covariates, the additional variance explained by batch after adjustment for those covariates, and the extent to which observations remain clustered by batch.

In each case, the models are fitted feature-wise, and several informative statistics using each model’s outputs are made. A linear regression model is first fitted using only the supplied biological and technical covariates. From this, we derive a covariate-only *R*^2^ value by comparing the residual and total sum of squares as shown in the supplementary materials. A second linear regression includes both the measured covariates and batch as fixed effects. By similarly deriving the total *R*^2^ for this model, we can compare the covariate-and-batch to the covariate-only *R*^2^, to assess the unique contribution of batch to the model fit. Covariatespecific partial *R*^2^ values are reported to quantify the proportion of feature variance associated with each supplied covariate after accounting for other model terms. Comparison before and after harmonisation provides an indication of how covariate-associated variation changes following correction. Preservation of similar partial *R*^2^ values is consistent with retention of covariate-associated structure, whereas increases may reflect greater relative prominence or detectability of these associations following removal of technical variation. However, increases in partial *R*^2^ should not in isolation be interpreted as improved preservation, as harmonisation may alter both residual variance and the relative variance attributable to the model term. Finally, a random-intercept linear mixed-effects model is fitted using batch-specific random intercepts. Unlike the fixed-effects regression, this model estimates the variance attributable to between-batch differences rather than estimating a separate coefficient for each batch. The random-intercept mixed-effects model is used to estimate the Intraclass Correlation Coefficient (batch-ICC) when batch is included as a random effect. The batch-ICC estimates the proportion of unexplained variance attributable to clustering within batches, with larger values indicating stronger batch-dependent clustering.

##### Linear modelling in the longitudinal tool

For the repeated-measures workflow, additive batch effects are tested using a mixed-effects model in which batch and covariates are included as fixed effects and the subject is included as a random effect (Beer et al., 2020). A full model including batch is compared with a reduced model excluding batch using a Kenward–Roger F-test, yielding estimates of the significance of additive batch effects. This model comparison yields an omnibus test of whether batch explains additional variance in the features beyond biological covariates. While the omnibus model comparison provides a global assessment of additive batch effects, it may obscure site-specific mean differences between individual batches. To assess batch effects at the level of individual site pairs, we perform pairwise comparisons using linear contrasts of the estimated fixed effects. For each pair of sites, we evaluate the statistical significance of the model-estimated difference in features. Additionally, we assess the preservation of meaningful variation after harmonisation. We partition the variance into between-subject variance, reflecting stable inter-individual differences, and residual variance, capturing within-subject variability, measurement noise, and other unexplained sources of variation. From these variance components, we compute the subject ICC as the ratio of between-subject variance to total variance. An increase in ICC after harmonisation indicates greater relative prominence of between-subject variation. To assess the preservation of biological variability, a linear mixed-effects model is fitted with user-specified biological and/or technical covariates as fixed effects and a subject-specific random intercept to account for repeated measurements. Feature-wise effect size estimates, 95% confidence intervals, and corresponding p-values are reported for the specified covariates, enabling evaluation of whether known biological or technical associations are preserved following harmonisation. These estimates are particularly informative in settings where little or no systematic change is expected. For example, in short-interval test–retest data, the emergence or amplification of substantial timepoint effects after harmonisation may indicate the introduction of spurious longitudinal variability rather than genuine biological change.

### 2.6 Batch effect on data variance and distributions

Although batch effects are often discussed in terms of shifts in the mean, differences in scale can be equally important. Even when batch-specific means have been aligned, differences in variance can affect downstream inference, the stability of machine-learning models, and the calibration of normative models. For example, a feature with inflated within-batch variance may appear less reliable, contribute unevenly to model fitting, and distort estimates of atypicality or uncertainty. For these reasons, assessing scale differences between batches is a necessary complement to evaluating mean differences, particularly in workflows intended to preserve biologically meaningful variation while removing technical bias.

To assess differences in variance between batches, the cross-sectional workflow uses the F-test and the Brown-Forsythe test for the homogeneity of variance between groups as well as a two-sample Kolmogorov-Smirnov (KS) test, which is sensitive to not only differences in data variance across empirical distributions but also differences in the distributions themselves. In the repeated-measures workflow, the Fligner–Killeen test is used as a robust, non-parametric test of homogeneity of variance across batches, providing an assessment of residual differences in scale that is less sensitive to departures from normality (Beer et al., 2020). The methods are summarised in Table 3.

**Table 3:** Three methods for analysing differences in the variance and distribution between batches.

| Method | Reported Metric | Description |
|---|---|---|
| F-test | Ratio of Variance | The ratio of observed within-batch variance compared to whole-sample variance |
| KS-test | KS statistic and p-value | The difference in the empirical distri-butions of each batch's data |
| Fligner-Killeen test | ChiSq and p-value | Test of variance equivalence that is ro-bust to non-normal data and extreme outliers |

#### 2.6.1 F-test of variance differences

The F-test provides a direct comparison of the ratio of variances between two batches and is therefore useful under the assumption of approximate normality. As with Cohen’s d, this is performed on the covariate-adjusted data as opposed to the unadjusted data. The results of the F-test are displayed separately for each batch as feature-wise estimates of the ratio of variance (*V_r_*) between that batch and all other batches by the single dataset reporting tool. For the comparison tool, the results are instead shown as the feature-wise average absolute log values of all calculated 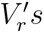. We express it this way such that a single plot can be given for each harmonisation method tested and so that variance ratios below one are expressed similarly to those above one. In each case, deviations from one and from zero, the single dataset and comparison tools, respectively, indicate differences in scale between batches and would be expected to improve after harmonisation.

#### 2.6.2 Fligner-Killeen test of variance homogeneity

Because variance-based tests can be sensitive to deviations from normality, we additionally use the Fligner-Killeen test, an inferential statistic used to assess the homogeneity of the variance between one or more groups. The Fligner-Killeen test is an alternative way of applying Levene’s test that uses the median as opposed to the mean when estimating variance and uses normalised ranked deviations rather than absolute deviations, making it more robust when applied to potentially non-normal data. The Fligner-Killeen test is applied to each feature and unique batch combination independently for both the raw data and the residuals after accounting for covariate effects. In repeated-measures data, variance differences must be distinguished from stable between-subject heterogeneity. The variance diagnostic is therefore applied to residuals obtained after accounting for temporal effects and subject-level dependence in addition to biological covariates.

#### 2.6.3 Two-sample Kolmogorov-Smirnov test of empirical distributions

The two-sample KS test compares the empirical cumulative distributions of two batches. It calculates the absolute difference between the two distributions across the range of observed feature values and uses the largest observed difference as the KS statistic. The test does not assume a specific distribution and is sensitive to both mean and scale differences. A significant result indicates differences in the underlying distribution of each batch. As differences in the distributions can come from underlying population differences, this test is performed on the residuals of the data, with a significant result indicating residual batch-associated differences in the distributions.

### 2.7 Covariance analysis and dimensionality reduction

While feature-wise analyses can effectively quantify bias in individual derived measures, they may not fully characterise the overall impact of batch differences on a dataset. For example, when differences are spatially heterogeneous (e.g., those related to field inhomogeneities), the magnitude, direction, and nature of batch effects are also likely to vary across regions. Therefore, a comprehensive assessment of batch effects should also consider multivariate approaches that take into account relationships between features (Table 4. Such methods enable the assessment of whether biological and anatomical dependencies between derived measures are preserved across batches, providing a more holistic evaluation of harmonisation need and effectiveness.

**Table 4:**
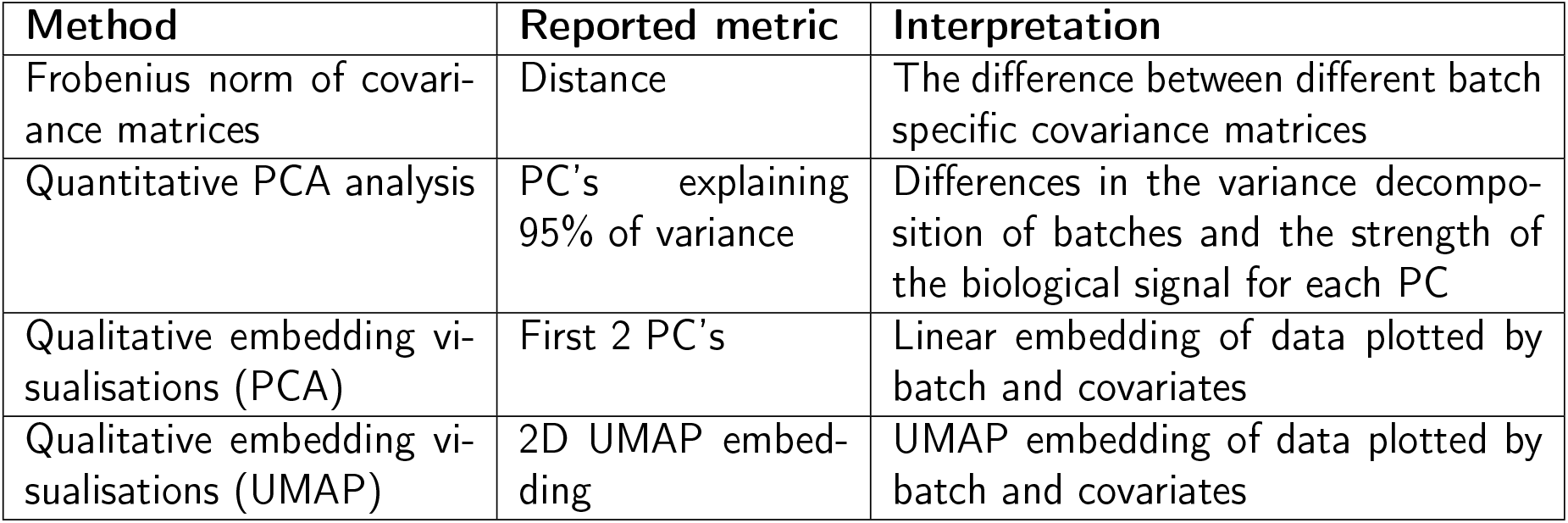
The four approaches (qualitative and quantitative) for analysing differences in the covariance between batches, visualising data structure and assessing batch and covariate contributions.

| Method | Reported metric | Interpretation |
|---|---|---|
| Frobenius norm of covari-ance matrices | Distance | The difference between different batch specific covariance matrices |
| Quantitative PCA analysis | PC’s explaining 95% of variance | Differences in the variance decompo-sition of batches and the strength of the biological signal for each PC |
| Qualitative embedding vi-sualisations (PCA) | First 2 PC’s | Linear embedding of data plotted by batch and covariates |
| Qualitative embedding vi-sualisations (UMAP) | 2D UMAP embed-ding | UMAP embedding of data plotted by batch and covariates |

#### 2.7.1 Analysis of covariance using the Frobenius norm

Batch effects may influence not only the mean and variance of individual features but also the relationships between features. In neuroimaging datasets, many derived measures are biologically correlated due to shared anatomical, developmental, or pathological processes. Technical effects that differentially affect specific regions or measurements may therefore alter the covariance structure of the dataset even when feature-wise means appear similar. This is of particular concern in harmonisation scenarios where there are significant differences between the scanners themselves, such as magnetic field strength, *B*0 inhomogeneities or pulse sequences, which will have a technical bias that is spatially dependent.

To assess whether batches differ in their covariance structure, the covariance between each of the features (columns) in the data is calculated separately for each batch. Pairwise differences between the resulting batch covariance matrices are then calculated using the Frobenius norm. The Frobenius norm between the covariance matrices provides a single metric measuring the differences in covariance structure between batches. Larger Frobenius distances indicate greater disagreement in the covariance structure between batches and may suggest the presence of complex technical effects that alter relationships between imaging features. Such effects are unlikely to be detected using feature-wise analyses alone and may persist even after mean and variance differences have been removed through harmonisation. To improve robustness, covariance comparisons should be restricted to features with sufficient observations in each batch. Results should be interpreted cautiously when batch sample sizes are small relative to the number of features.

#### 2.7.2 Principal Component Analysis

Principal Component Analysis (PCA) transforms the feature matrix into an orthogonal set of latent variables (Principal Components, PCs) which are ordered by the proportion of total variance explained. By transforming the data in this way, we are able to assess how batch labels and covariates contribute to the overall variance structure of the dataset. We first provide visualisations using scatter plots of PC1 and PC2, colouring observations by batch before plotting again for each covariate. Prior to harmonisation, distinct clusters may be visible by batch, and covariate trends or groupings, in the case of categorical covariates, may be less obvious. This is true for both cross-sectional and repeated-measures data, but in the later setting, distinct clusters by subject ID may also be desired.

Additionally, to provide a quantitative measure of batch and covariate associations with the principal components, an omnibus coefficient of determination (*R*^2^) is calculated separately for all covariates and the batch label for each principal component. Continuous covariates are included as continuous predictors, while categorical variables, including batch, are represented using categorical indicator variables. The resulting *R*^2^ therefore quantifies the proportion of variation in each principal component associated with a given variable and, for categorical variables, is invariant to the numerical or lexical coding of the variable, so the order of categories does not matter. These values are visualised as a heatmap for the first five principal components. High *R*^2^values for batch indicate that batch membership is strongly associated with dominant variation in the feature space, while associations with biological covariates indicate biologically relevant organisation of the data. Where batch and biological covariates are both strongly associated with the same principal components, the resulting associations should be interpreted cautiously because they may reflect confounding between technical and biological variation. For continuous or binary variables, Pearson or point-biserial correlations may alternatively be used; however, we report omnibus *R*^2^ because correlations are not appropriate for nominal variables with multiple categories, such as batch, whose numerical encoding is arbitrary. Per-batch scree and cumulative variance curves are also generated to compare the variance explained by each principal component across batches, with differences indicating potential batch-related differences in multivariate structure.

#### 2.7.3 Cluster analysis using dimensionality reduction/embedding

While PCA is a good approach for the analysis and visualisation of linear sources of variation, batch effects may also manifest as non-linear multivariate structure within the data. To capture this, we use Uniform Manifold Approximation (UMAP) for visualising non-linear effects (McInnes et al., 2020). UMAP constructs a weighted nearest-neighbour graph in the original feature space and seeks a low-dimensional embedding that preserves local neighbourhood relationships. A full explanation of the underlying topological assumptions and derivation of the algorithm can be found in McInnes et al. (2020).

While UMAP embeddings can be used as inputs to downstream analyses such as clustering or machine learning models, within DHARM they are primarily intended as a visual diagnostic tool for exploring non-linear relationships between imaging features, batch membership, and biological covariates. UMAP is stochastic and depends on several parameters. These are set to sensible defaults by the tool but can also be specified by the user. For ease of use and visualisation, the number of components is set by default to 2, and the distance metric is set to Euclidean. As with PCA, the resulting embeddings are visualised as scatter plots coloured according to batch membership or covariate values.

### 2.8 Evaluation metrics for test–retest and travelling-head settings

Test–retest metrics provide a controlled framework for quantifying technical variability and measurement reproducibility, as substantial biological changes are generally not expected over the relatively short interval between repeated acquisitions. With test–retest and travelling-head datasets increasingly incorporated into the design and validation of clinical and research studies (Markiewicz et al., 2026; Warrington et al., 2025), these metrics provide quantitative measures of repeat-measure consistency and facilitate systematic characterisation of measurement stability and technical sources of variation.

#### 2.8.1 Within-subject variability

Within-subject variability is quantified using pairwise relative percent difference (RPD) across repeated measurements. RPD is used as a symmetric measure of relative disagreement that can be applied consistently to datasets with two or more, potentially unbalanced, repeated observations. For each participant and measurement, RPD is calculated for each pair of available repeated measurements as the absolute difference between the measurements divided by their mean, expressed as a percentage. Where more than two repeated measurements are available, the median pairwise RPD is used to summarise within-subject variability for each measurement. Lower RPD values indicate greater consistency across repeated measurements, whereas higher values indicate greater within-subject variability. The tool reports participant-and feature-specific RPD values, allowing participants with unusually high overall variability and features with relatively high or low measurement variability to be identified. In test-retest settings, a reduction in RPD following harmonisation can therefore be interpreted as improved measurement consistency and reduced unwanted non-biological variability.

#### 2.8.2 Subject order consistency

Subject-order consistency is quantified using Spearman rank correlation between matched participants across repeated acquisition pairs, following the rank-consistency principle used in travelling-head studies such as ON-Harmony (Warrington et al., 2023). Each acquisition pair is evaluated using participants available in both acquisitions, allowing the metric to accommodate unbalanced repeated-measures designs. Spearman’s rho (*ρ*) is calculated separately for each measurement, with higher values indicating greater preservation of the relative ordering of participants across repeated acquisitions. Statistical significance is assessed using permutation testing, in which measurements from the second acquisition are randomly permuted across matched participants to generate a null distribution of correlation values. The tool reports feature-specific correlations across acquisition pairs, allowing features with relatively high or low rank consistency to be identified. In test-retest and longitudinal settings, where participant-specific ordering is expected to remain relatively stable, an increase in Spearman’s rho following harmonisation can therefore be interpreted as improved preservation of subject-order structure and reduced influence of unwanted technical variability.

### 2.9 Summary of methods

The outputs from each of the methods used by the reporting part of the tool are organised into a single structured HTML file. The report contains descriptions of each method, plots showing the results and a brief summary of the highlighted results. The tool also returns the results for each test as CSV files. We additionally offer a reduced version of the cross-sectional report which runs a less extensive test suite. The comparison versions of the cross-sectional and longitudinal reports, respectively, use the same tests as their single dataset counterparts. A table showing which tests are used in each of the reporting tools is shown in Table 5

**Table 5:** Table indicating whether a given test is used in each of the reporting tools. Metrics suitable for test-retest settings are marked with *.

| Evaluation metric | cross-sectional min | cross-sectional full | Repeated-measures |
|---|---|---|---|
| Cohen's d | ✗ | ✓ | ✓ |
| Mahalanobis distance | ✗ | ✓ | ✓ |
| Linear mixed and fixed-effect modelling | ✓ | ✓ | ✓ |
| F-Test | ✓ | ✓ | ✗ |
| Fligner-Killeen | ✗ | ✓ | ✓ |
| KS-test | ✗ | ✓ | ✗ |
| Frobenius norm | ✗ | ✓ | ✗ |
| PCA | ✗ | ✓ | ✓ |
| UMAP | ✗ | ✓ | ✗ |
| Subject order consistency* | ✗ | ✗ | ✓ |
| Within-subject variability* | ✗ | ✗ | ✓ |

### 2.10 Demonstration of DHARM

We demonstrate DHARM in both cross-sectional and repeated-measures settings using a selection of the harmonisation methods implemented within the tool. In each setting, DHARM is first applied to the unharmonised data to characterise the magnitude and nature of the baseline batch effects, and is then applied again following harmonisation. Comparing the same diagnostic metrics before and after harmonisation allows the extent to which technical variation has been reduced to be assessed alongside the preservation of biologically meaningful variation. Where multiple harmonisation approaches are considered, their outputs can be evaluated using the same set of diagnostic criteria, providing a consistent framework for comparing their relative performance and identifying method-specific strengths and limitations. Here, we show the outputs of the comparison reports for both the cross-sectional and repeated-measures demonstrations, running the reporting tool using both the unharmonised and all harmonised datasets.

#### 2.10.1 Cross-sectional demonstration

We demonstrate DHARM using structural measurements from the four UK Biobank sites (Stockport, Newcastle, Reading and Bristol) (Alfaro-Almagro et al., 2018; Miller et al., 2016), the Cambridge Centre for Ageing and Neuroscience neuroimaging dataset (Cam-CAN) (Shafto et al., 2014; Taylor et al., 2017) and the Oxford UK Biobank Aligned Controls dataset (OxBAC) (Griffanti et al., 2025), a subset of healthy, age-matched controls from the Oxford Brain Health Clinic study (Blane et al., 2025; O’Donoghue et al., 2023). These datasets were all acquired at different sites with similar scanners and acquisition parameters (for the T1 and FLAIR data) but very different demographics. We show here the data for 244 cortical thickness measures derived using the FreeSurfer implementation in the UKB pipeline from Alfaro-Almagro et al. (2018) (Fischl, 2012). The specific features are provided in the supplementary materials. A summary of the datasets, including their demographic and imaging acquisition characteristics, is provided in Table 6.

**Table 6:** Table showing age, sex, sample size and some scanner information for the five datasets used in the cross-sectional demonstration of DHARM, as well as original references.

| Dataset | Sample size used | Age range (years) | Percentage Female | Scanner and dataset references |
|---|---|---|---|---|
| UKB: Stockport | 500 | 47-83 | 54% | Siemens Skyra 3T (Alfaro-Almagro, 2021; Miller et al., 2016) |
| UKB: New-castle | 500 | 50-84 | 56.6% | Siemens Skyra 3T(Alfaro-Almagro, 2021; Miller et al., 2016) |
| UKB: Reading | 500 | 50-82 | 57.6% | Siemens Skyra 3T(Alfaro-Almagro, 2021; Miller et al., 2016) |
| UKB: Bris-tol | 500 | 50 - 84 | 54.6% | Siemens Skyra 3T(Alfaro-Almagro, 2021; Miller et al., 2016) |
| Cam-CAN | 474 | 18 - 89 | 50% | Siemens TIM Trio (Shafto et al., 2014) |
| OxBAC | 96 | 60 - 92 | 54.1% | Siemens Prisma scanner (Blane et al., 2025; Griffanti et al., 2025) |

#### 2.10.2 Harmonisation approaches

We apply five different harmonisation approaches here, each with three biological covariates included to be preserved in the model: Age, sex and head-size scaling, derived using SIENAX as part of the FSL software package (Jenkinson et al., 2012; Smith et al., 2002). All harmonisation was performed using DHARM. 1. Linear regression (OLS) — A simple ordinary least squares regression was used to model and remove batch effects while accounting for covariate effects. 2. ComBat — An empirical Bayes harmonisation method that models and removes additive (mean) and multiplicative (variance) batch effects at the feature level, using information pooled across features to improve parameter estimation while preserving covariates of interest(Fortin et al., 2017; Johnson et al., 2007). 3. CovBat — An extension of ComBat that additionally corrects for batch effects present in the covariance structure of the data, by applying ComBat-style correction to the principal component scores of the ComBat residuals (Chen et al., 2022). 4. ComBat with B-splines — A variant of ComBat in which continuous covariates (e.g., age) are modelled using spline basis expansion rather than assuming a strictly linear relationship, allowing for non-linear covariate effects during harmonisation (Pomponio et al., 2020). 5. CovBat with a reference batch — CovBat harmonisation performed with a specified batch (UKB Stockport) set as the reference, so that all other batches were harmonised to match the reference batch rather than pooled/global average (Turnbull et al., 2026).

### 2.11 Repeated-measures evaluation

The longitudinal implementation of DHARM is demonstrated using the ON-Harmony multisite, travelling-head dataset (Warrington et al., 2023, 2025), which provides a setting in which biological change is expected to be minimal, allowing technical variation due to batch effects to be examined more clearly. The dataset comprised 20 healthy participants, each scanned on six 3T scanners across five sites and three major MRI vendors, with data acquired across five imaging modalities. In addition to the between-scanner repeated acquisitions, nine participants also underwent five additional within-scanner acquisitions, providing a within-scanner repeatability reference against which between-scanner variability could be evaluated. This combination of repeated measurements across and within scanners makes ON-Harmony a particularly informative test case for DHARM, enabling evaluation of test–retest metrics while also testing methods that explicitly account for within-subject dependence. For this demonstration, we analysed atlas-based cortical GM volumes (total 138) from T1w scans as in (Warrington et al., 2025). The list of imaging-derived phenotypes is provided in the supplementary materials. These measures were derived using an adapted version of the UK Biobank image processing pipeline and openly available on OpenNeuro.

#### 2.11.1 Harmonisation approaches

We applied three harmonisation approaches: ComBat, CovBat, and Longitudinal ComBat (LongComBat), with age and sex included as biological covariates to be preserved. ComBat and CovBat, described in the cross-sectional evaluation above, were retained as reference approaches that do not explicitly model within-subject dependence. Longitudinal ComBat (LongComBat) (Beer et al., 2020) extends ComBat by explicitly modelling the correlation between repeated measurements from the same participant. Although originally developed for longitudinal multi-scanner data, its inclusion allowed us to examine whether explicitly accounting for repeated-measures dependence provides additional benefit over approaches that treat observations as independent. All harmonisation methods were applied using DHARM. Raw and harmonised data were evaluated using the test–retest metrics described above, alongside the broader DHARM metrics developed for repeated-measures observations.

## 3 Results

### 3.1 Cross-sectional demonstration

Cortical-thickness measures exhibited the largest apparent differences between batches and are presented here as a representative example of comparing different harmonisation approaches with DHARM. The figures presented were produced by applying the cross-sectional comparison tool after applying each harmonisation method. All figures are taken from the comparison report produced by the tool, covering shifts in feature location, differences in scale and empirical distributions, preservation of covariate-associated variation, and residual multivariate structure. The full report is provided in the supplementary materials; here, we focus on a representative subset of diagnostics. In the standard single dataset reports, Cohen’s d, F-tests, Fligner-Killeen tests and the results for the two-sample KS test are displayed per feature separately for each batch. In the comparison report, this is instead shown as a summary of the results for all batch-wise results. This is done to make the reports more readable when many different batches are used. However, as with the single dataset reporting tool, the outputs of each test are still returned as CSV files.

#### Global visual screening of batch differences

The report visualises the overall distribution of different batches of data as histograms of all median-centred Z-scores (Figure 3). The initial batch-wise distribution plots showed clear differences between the raw datasets, with visible displacement and differences in the overall shape of the pooled robust-standardised cortical-thickness distributions. Following harmonisation, the distributions showed substantially greater overlap, indicating that all approaches reduced the most visually apparent between-batch differences. The degree of overlap nevertheless varied between approaches, demonstrating that apparently similar correction of the global distributions does not necessarily imply equivalent harmonisation at the individual-feature level.

**Figure 3:**
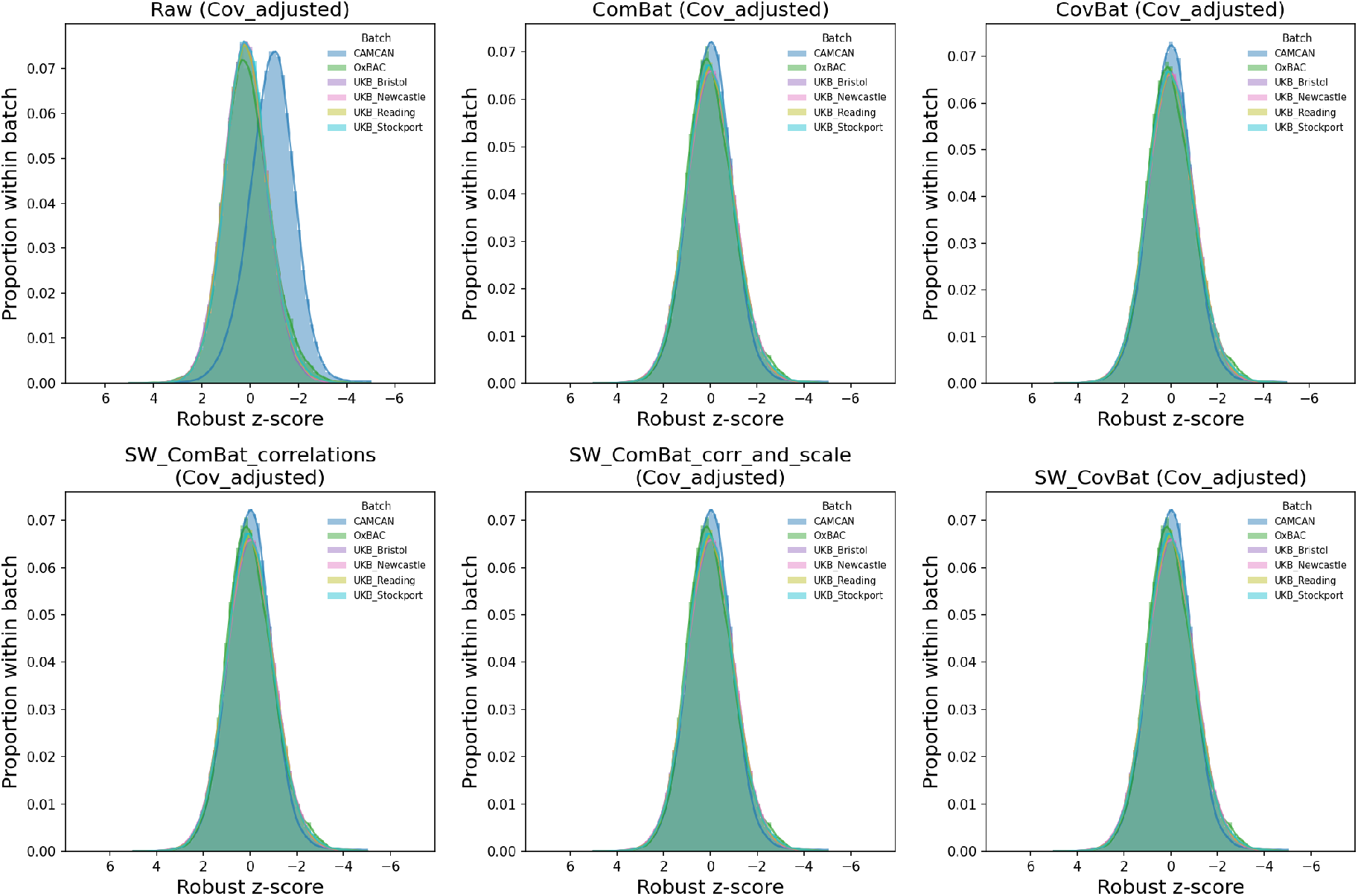
Distribution of feature-wise Z-scores across all features for each harmonisation method and the raw data after adjusting for covariates. In the full report, the covariate-adjusted and covariate-unadjusted distributions are plotted alongside each other.

The analysis of differing covariate distributions showed large differences in the ages of the chosen batches, with Cam-CAN having the youngest average population but also the widest age range. The UKB sites all had approximately identical age distributions, and the OxBAC participants were older on average with a much narrower age range. There were very small differences between each batch in terms of the sex split and head size distributions.

#### Additive batch effect analysis

Feature-wise, Cohen’s d provided a more sensitive assessment of residual additive effects (Figure 5). In the raw data, standardised mean differences were distributed across a broad range of values, indicating that the magnitude of the batch effect differed considerably between cortical-thickness measures. Following harmonisation, these effects were markedly reduced. Regression-based correction produced a particularly strong reduction in feature-wise mean differences, as expected for a procedure that directly removes linear batch-associated offsets. ComBat and CovBat likewise compressed the distribution of effect sizes towards zero, while the ComBat-GAM/B-spline implementation retained somewhat greater residual variation for some features.

Using the goodness of fit and relative contribution of a linear model with batch and biological covariates allows the assessment for the reduction of batch effects but also the preservation of biology after harmonisation. Figure 6 shows the goodness of fit of two models using the total *R*^2^ of the model, the first model (shown in blue) uses just the provided biological covariates, and the second (shown in orange) uses biological covariates and the batch variables. A large difference in the two models indicates batch contributes to a large portion of the total variance for a given feature. Figure 6 shows that there were large differences between the two models’ *R*^2^ for almost all features and that this difference was zero post-harmonisation, indicating batch contributed very little to the overall model fit.

**Figure 4:**
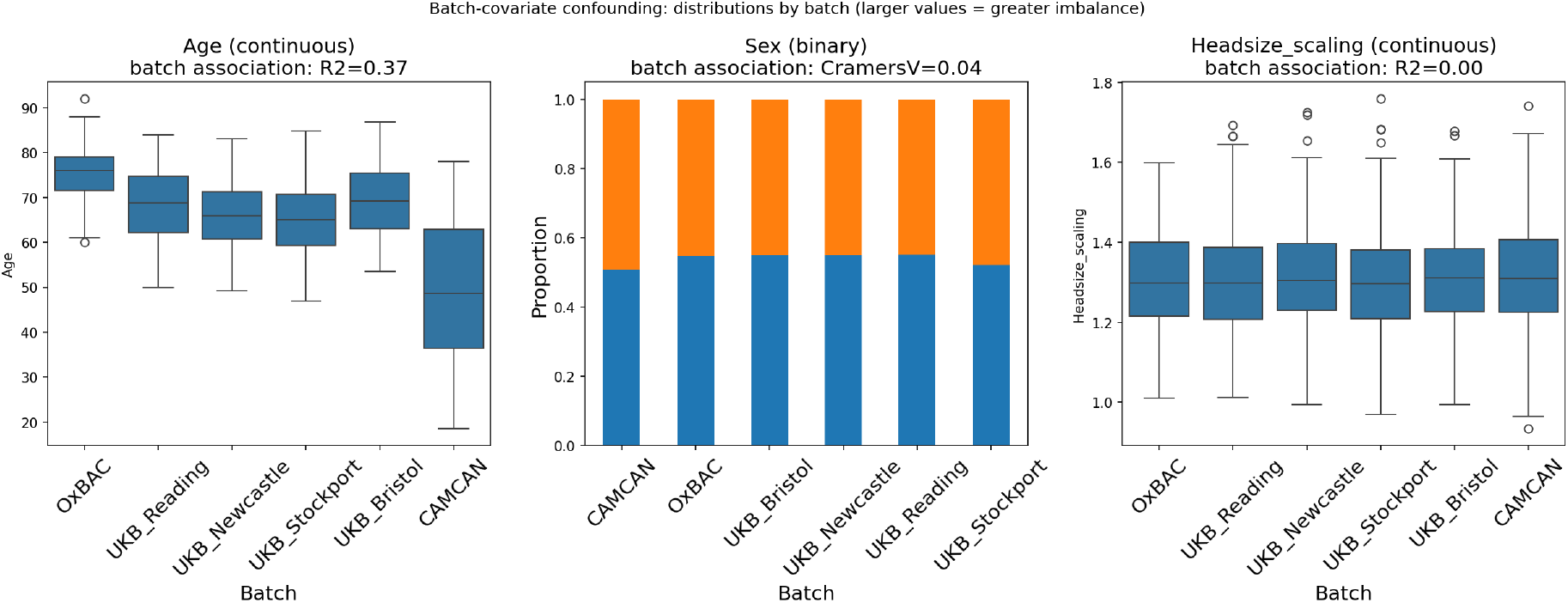
Three plots showing the covariate distribution by batch. Continuous covariates are plotted as box plots, and categorical and binary covariates as bar charts showing relative proportion.

**Figure 5:**
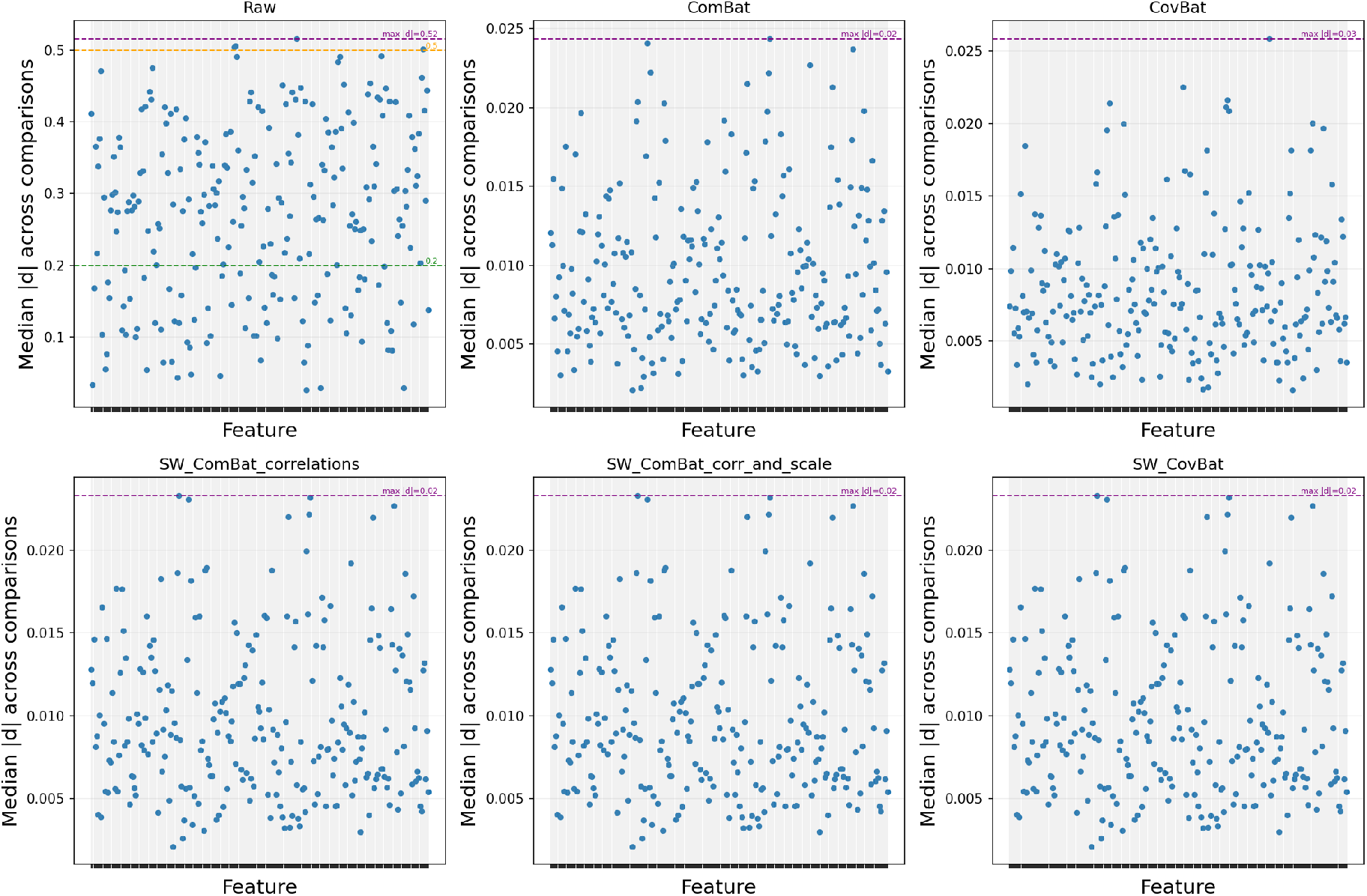
Magnitude of the average feature-wise Cohen’s d score (of the residual batch effect) for each method. Cohen’s d was averaged as the magnitude of the score for each batch against all the other batches, with a larger score showing greater disagreements in the means (a greater effect size) by batch on average for a given feature. The maximum average Cohen’s d score for a given method is shown by the purple horizontal line.

Additionally, the overall *R*^2^ of the covariate-only model improved after the application of each harmonisation method. The summary measures in Figure 7 provide a complementary view of this behaviour. The partial *R*^2^ associated with age increased substantially following harmonisation, whereas values associated with sex and head size were comparatively stable. This indicates that age-associated variation accounted for a greater proportion of the remaining feature variance after harmonisation.

**Figure 6:**
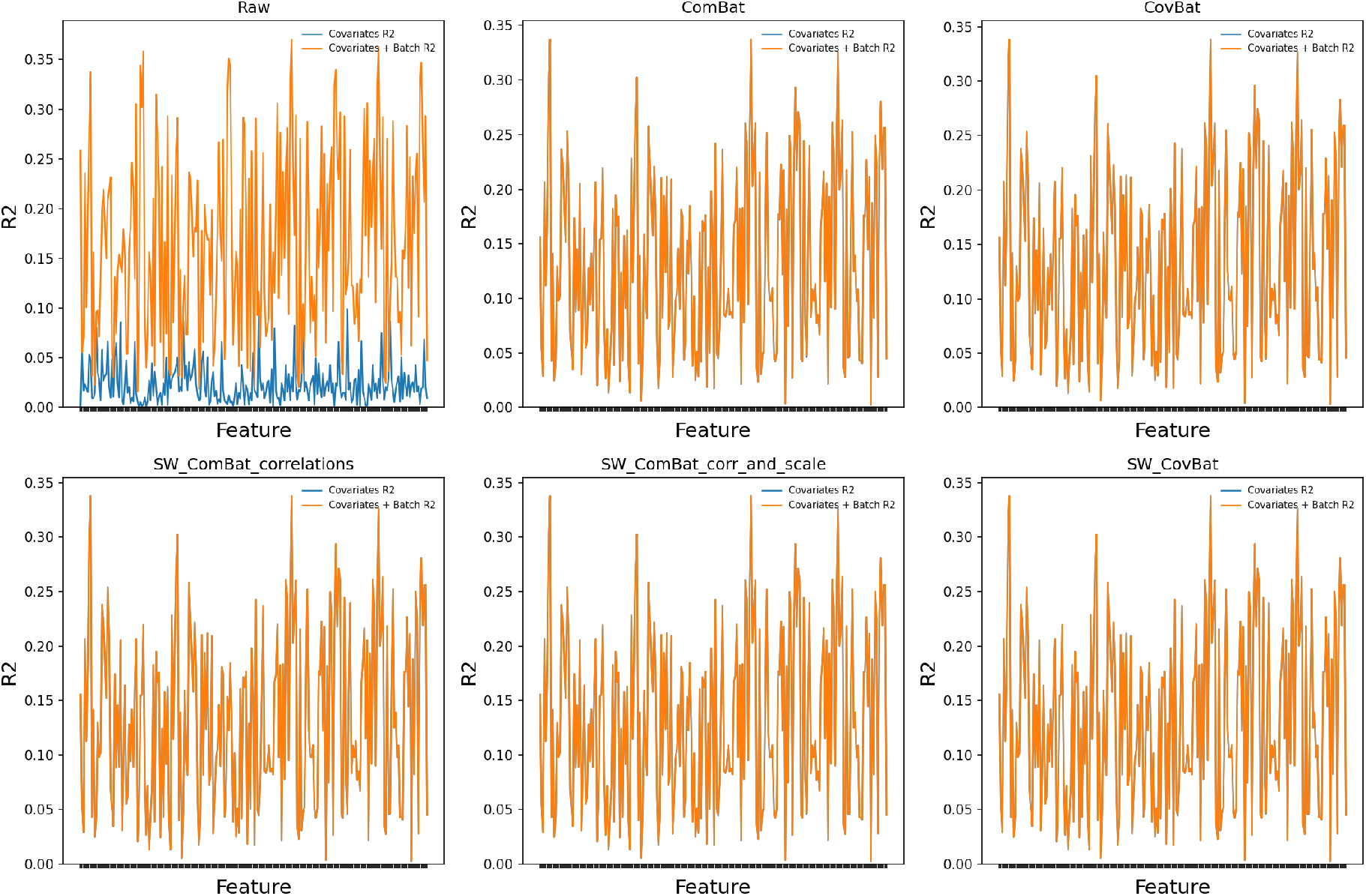
Feature-wise *R*^2^ of the two fixed-effect models, covariate only (blue) and covariate with batch (orange). *R*^2^ provides a measure of the goodness of fit of the model, with values closer to 1 indicating a higher proportion of explained variance for a given feature.

#### Batch effect on data variance and distributions

Differences in feature variance showed a more heterogeneous pattern than the additive effects (Figure 8). In the raw data, the variance-ratio profile varied substantially across cortical-thickness features, indicating that the batches differed not only in their average measurements but also in their scale. The size of these differences was strongly feature dependent, consistent with a technical effect that does not influence all cortical regions uniformly. The report calculates these quantities by comparing the variance within each batch with that of the remaining data and displaying the feature-wise result. Importantly, the harmonisation methods differed more clearly for this diagnostic than for the mean-effect analysis. Some approaches produced considerably more homogeneous variance profiles, whereas residual feature-specific peaks remained after others. This distinction is particularly informative when comparing simple regression with location-and-scale harmonisation approaches (i.e., ComBat). Removal of a batch-associated mean does not, by itself, guarantee that batch-specific variance has also been corrected, whereas ComBat-family methods explicitly allow adjustment of scale effects. The results therefore demonstrate that a dataset may appear well harmonised according to additive-effect metrics while continuing to contain measurable multiplicative batch structure.

**Figure 7:**
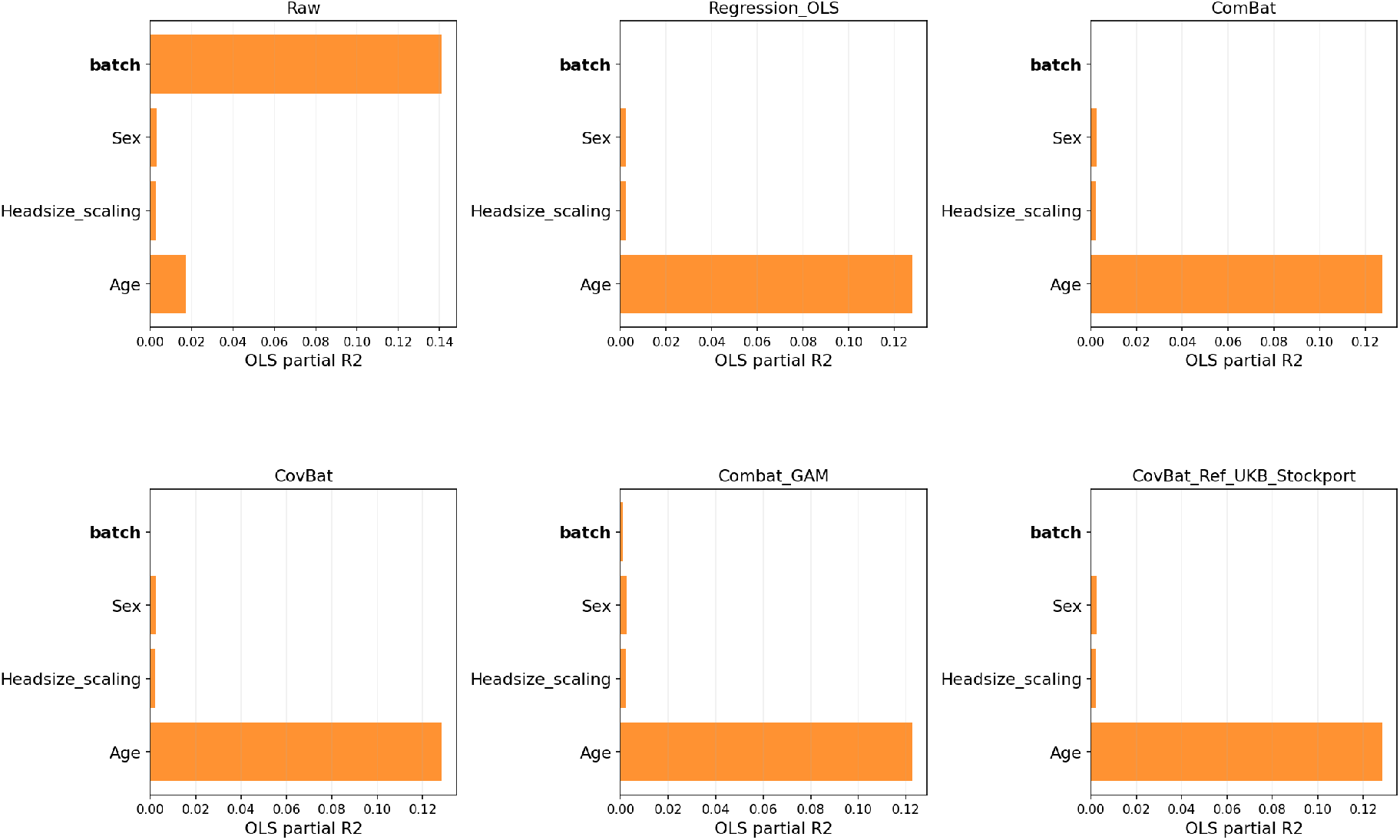
Magnitude of the partial *R*^2^ for covariates and batch, averaged across features. Covariate-specific partial *R*^2^ values indicate the proportion of variance attributable to each biological covariate after accounting for batch effects, and the batch-specific partial *R*^2^ summarises the strength of the batch effect across features.

**Figure 8:**
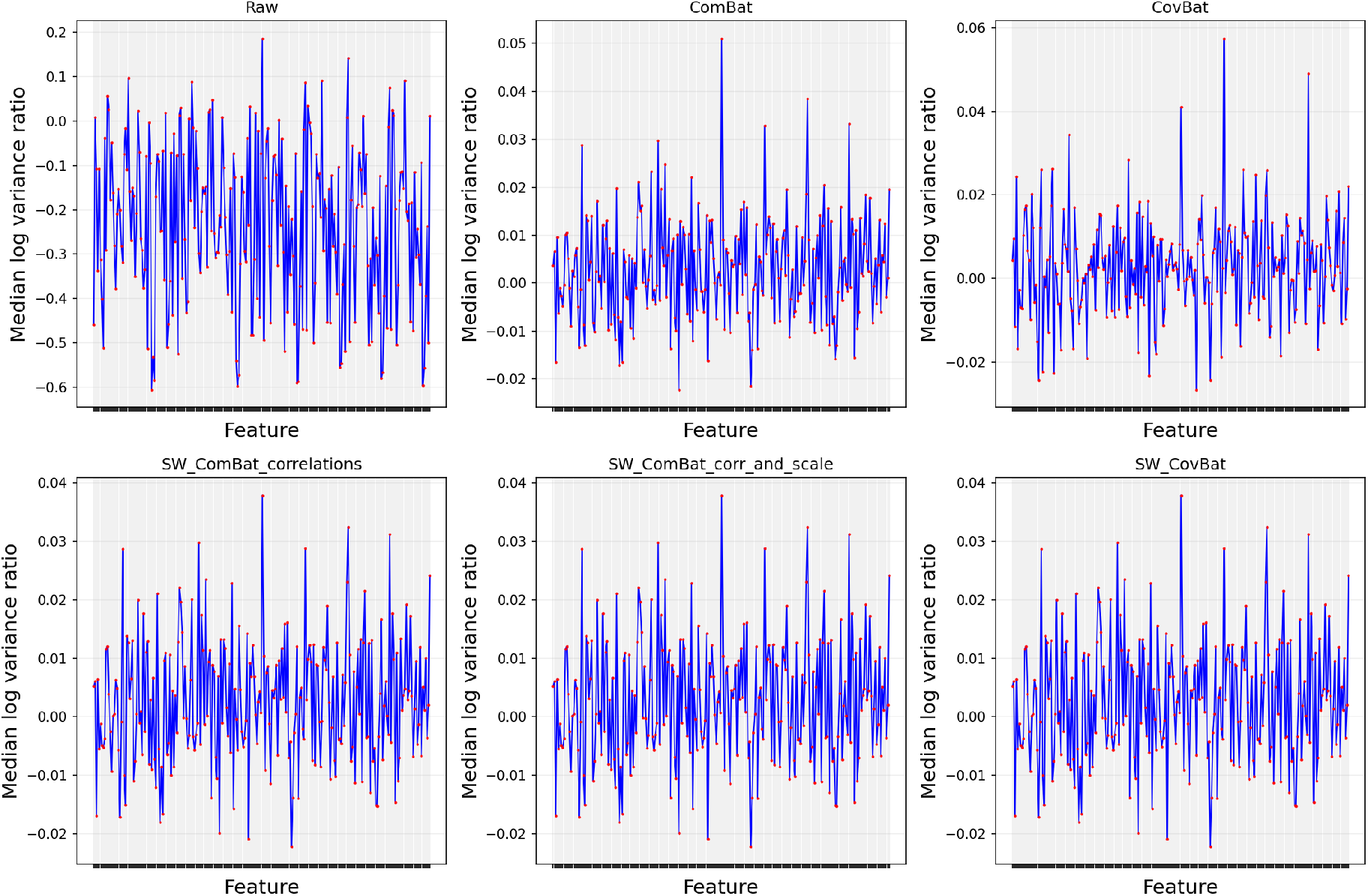
Comparison of variance ratio across methods. Median absolute log variance ratios summarise the overall deviation from perfect variance preservation, where values closer to zero indicate better alignment.

The two-sample Kolmogorov-Smirnov analysis provided a broader test of residual distributional differences by assessing each cortical-thickness feature independently (Figure 9). Whereas the previous diagnostics separately emphasise differences in feature location or scale, the KS statistic is sensitive to disagreement anywhere within the empirical distributions. The raw data contained a large number of features showing evidence of significant between-batch distributional differences. Following harmonisation, the number and magnitude of these differences were generally reduced, although the extent of reduction differed between methods. Overall, the KS analysis showed that removal of additive batch offsets did not automatically result in complete equality of the empirical feature distributions, reinforcing the value of evaluating distributional effects separately from differences in mean and variance.

**Figure 9:**
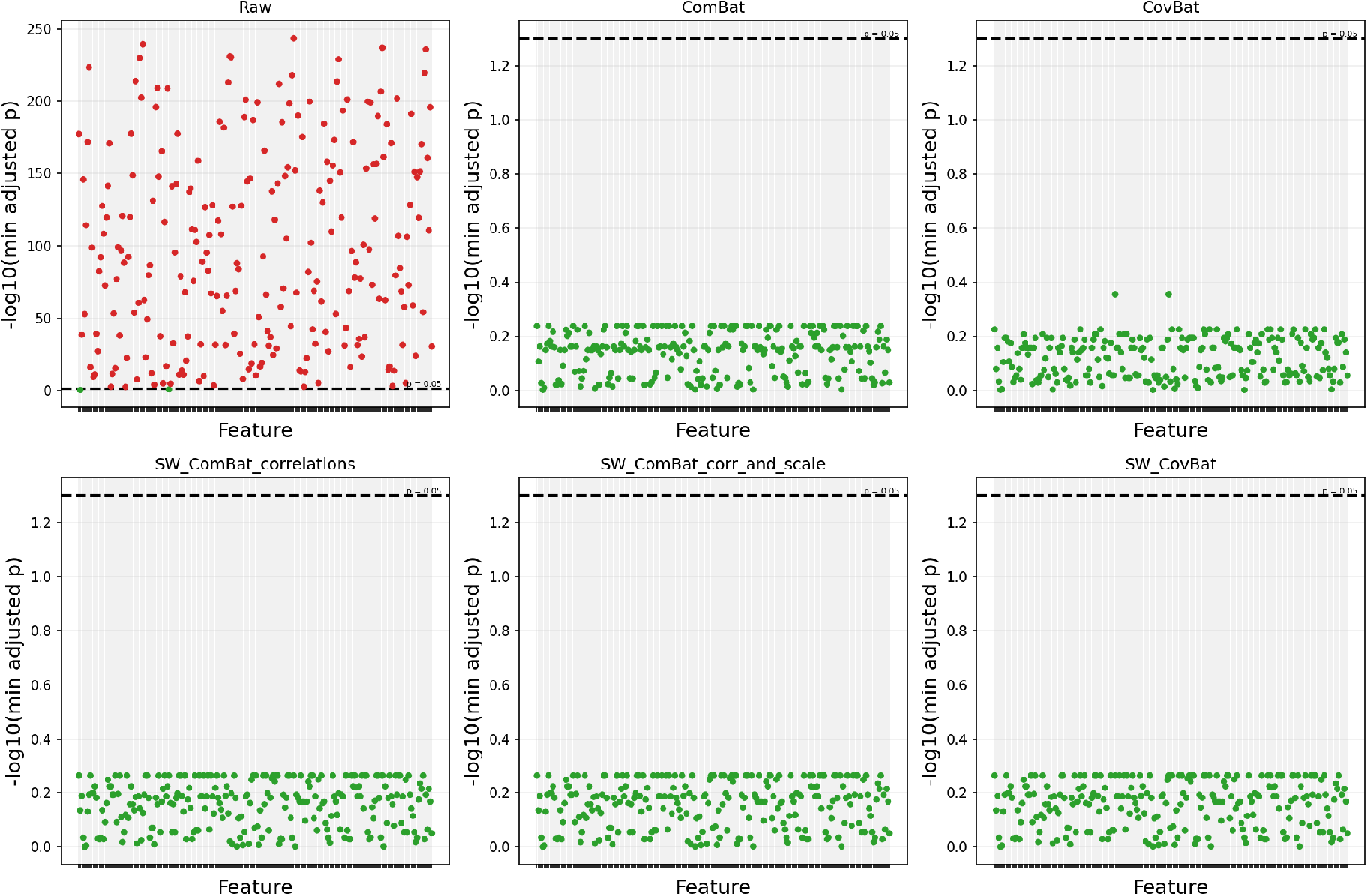
Feature-wise average KS score between each batch and all other batches. The black dotted line indicates the significance boundary. If a given marker is red, that indicates that one or more batches showed significant differences after correcting for multiple comparisons for this feature.

#### Covariance analysis and dimensionality reduction

PCA and UMAP were used to assess multivariate structure across all cortical-thickness features, with particular attention to the influence of batch and biological covariates before and after harmonisation. The omnibus *R*^2^ associations between batch, the supplied biological covariates, and the first five principal components are shown in Figure 10. Following harmonisation, the association with batch decreased slightly for the first principal component and was reduced to zero for the remaining components. In contrast, age-related associations strengthened, while those with sex and head size remained relatively stable.

**Figure 10:**
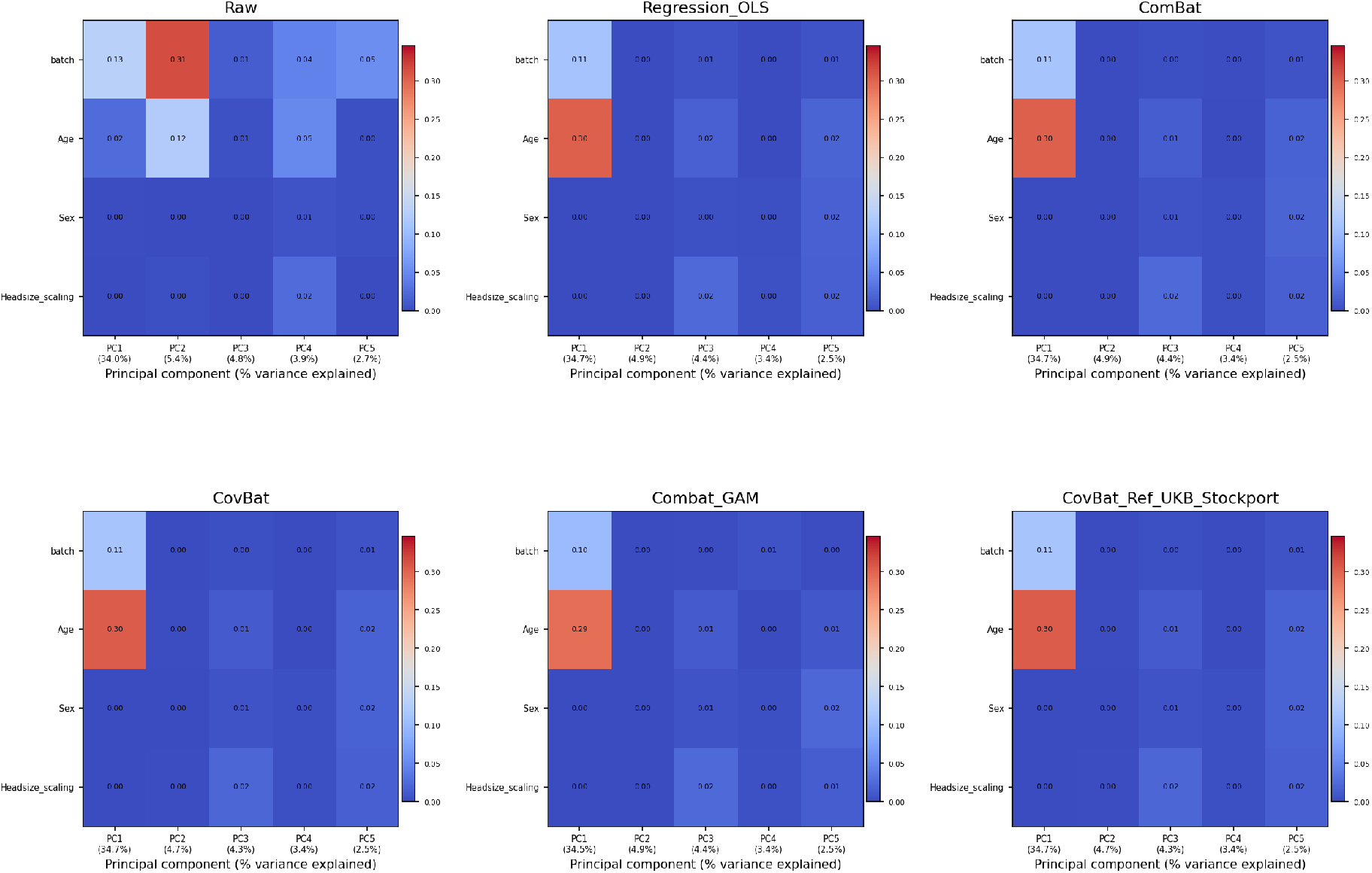
Heat map showing the omnibus *R*^2^ association of each supplied covariate and batch with each of the first five principal components.

UMAP provided a complementary visual assessment of these changes. In the raw embedding, observations exhibited visible batch-associated clustering, with certain regions disproportionately represented by individual batches (Figure 11). Following harmonisation, batches became substantially more intermixed across all approaches, although the geometry of the embeddings varied between methods. When coloured by age, the harmonised embeddings exhibited a more pronounced age-related gradient (Figure 12), consistent with the strengthened age associations observed in PCA. As UMAP coordinates have no direct physical interpretation, these findings are best understood in terms of changes in clustering and local organisation rather than absolute embedding positions. Together, the PCA and UMAP results suggest that harmonisation reduced technical structure while preserving, and potentially enhancing the visibility of, biologically meaningful variation.

**Figure 11:**
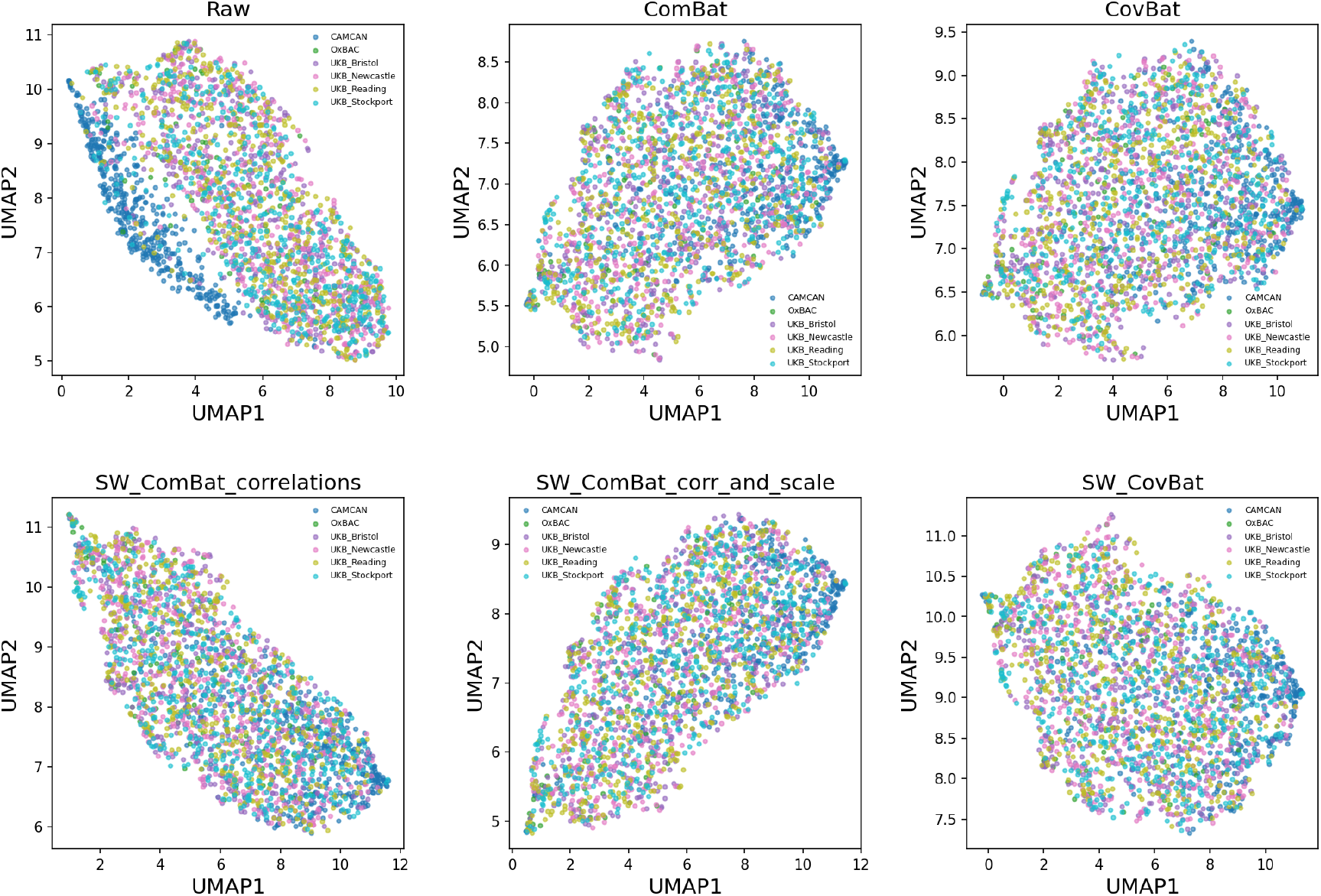
UMAP embedding results in two dimensions plotted by batch for each method of harmonisation. In the raw data, distinct batch clusters are clearly visible, with this being most obvious for the Cam-CAN data.

**Figure 12:**
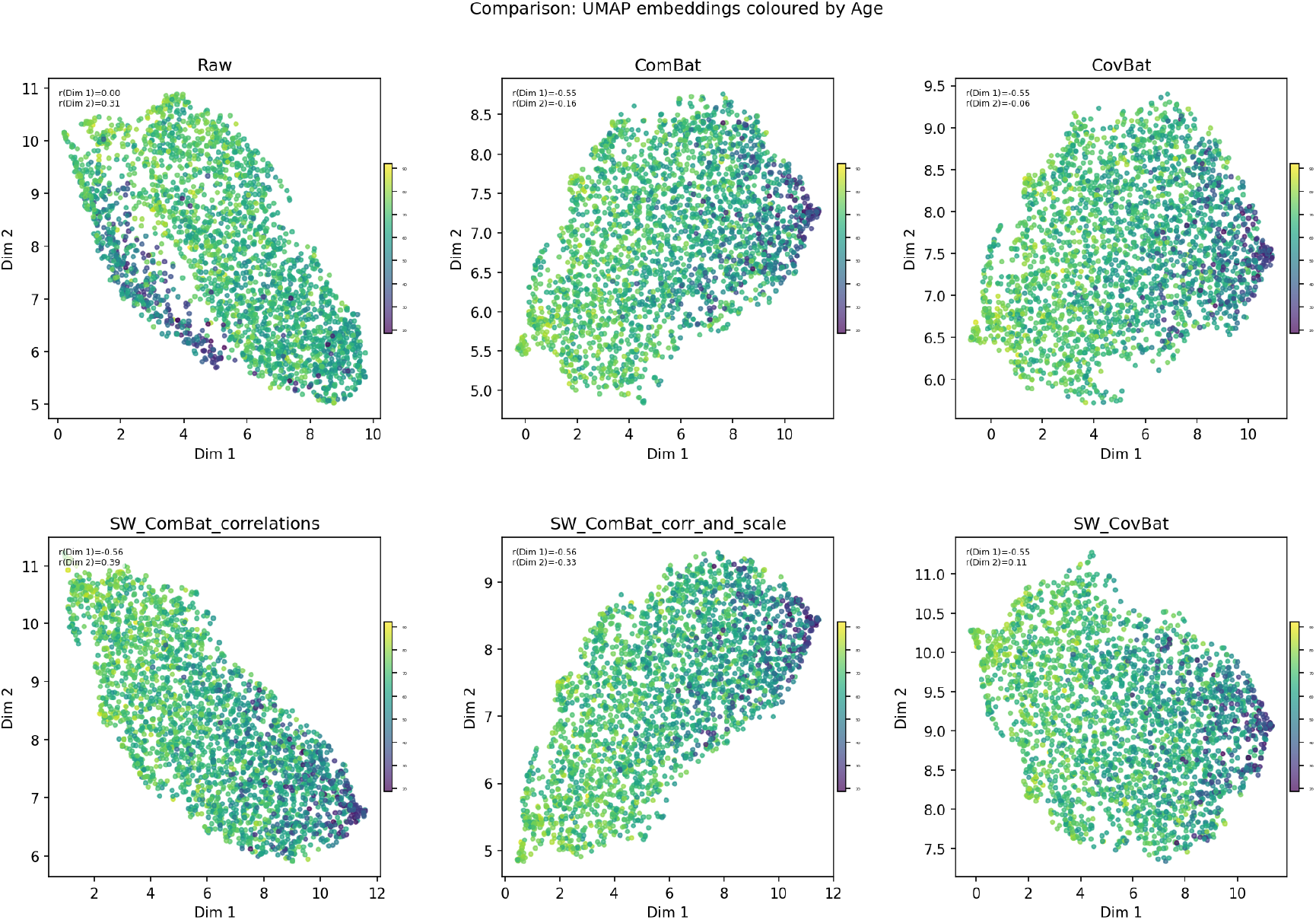
UMAP embedding results in two dimensions plotted by age for each of the methods of harmonisation, with the darker colours being younger in age. The correlation of age with each of the UMAP dimensions is shown for each method in a legend in the top left for each subplot.

### 3.2 Repeated-measures demonstration

DHARM generates a comprehensive diagnostic report that begins with dataset-level summaries, including the total number of observations, batch identifiers, batch sizes, and covariates (provided by the user), and then provides an overview of the evaluation workflow, along with a brief description of the methodology underlying each metric and general guidance on how to interpret the corresponding results. In the sections below, we show the outputs of the repeated measures implementation of DHARM comparing raw and harmonised volumes using ComBat, CovBat, and LongComBat, and outline how the metrics and visualisations can be used to evaluate harmonisation efficacy.

#### Global visual screening of batch effects

Before converting to robust Z-scores, a linear mixed effect model is fitted to the data which includes covariates and batch as fixed effects, with subject ID as a random intercept term. Batch effects are preserved and all other estimated effects are removed. The distribution of robust Z-scores showed substantial batch separation in both the mean and spread of the unharmonised/Raw data (Figure 13). Residual between-batch differences remained visually apparent following ComBat and CovBat, whereas the distributions following LongComBat showed greater overlap across batches. These distributional visualisations were interpreted alongside the quantitative batch-effect and repeated-measures consistency metrics.

**Figure 13:**
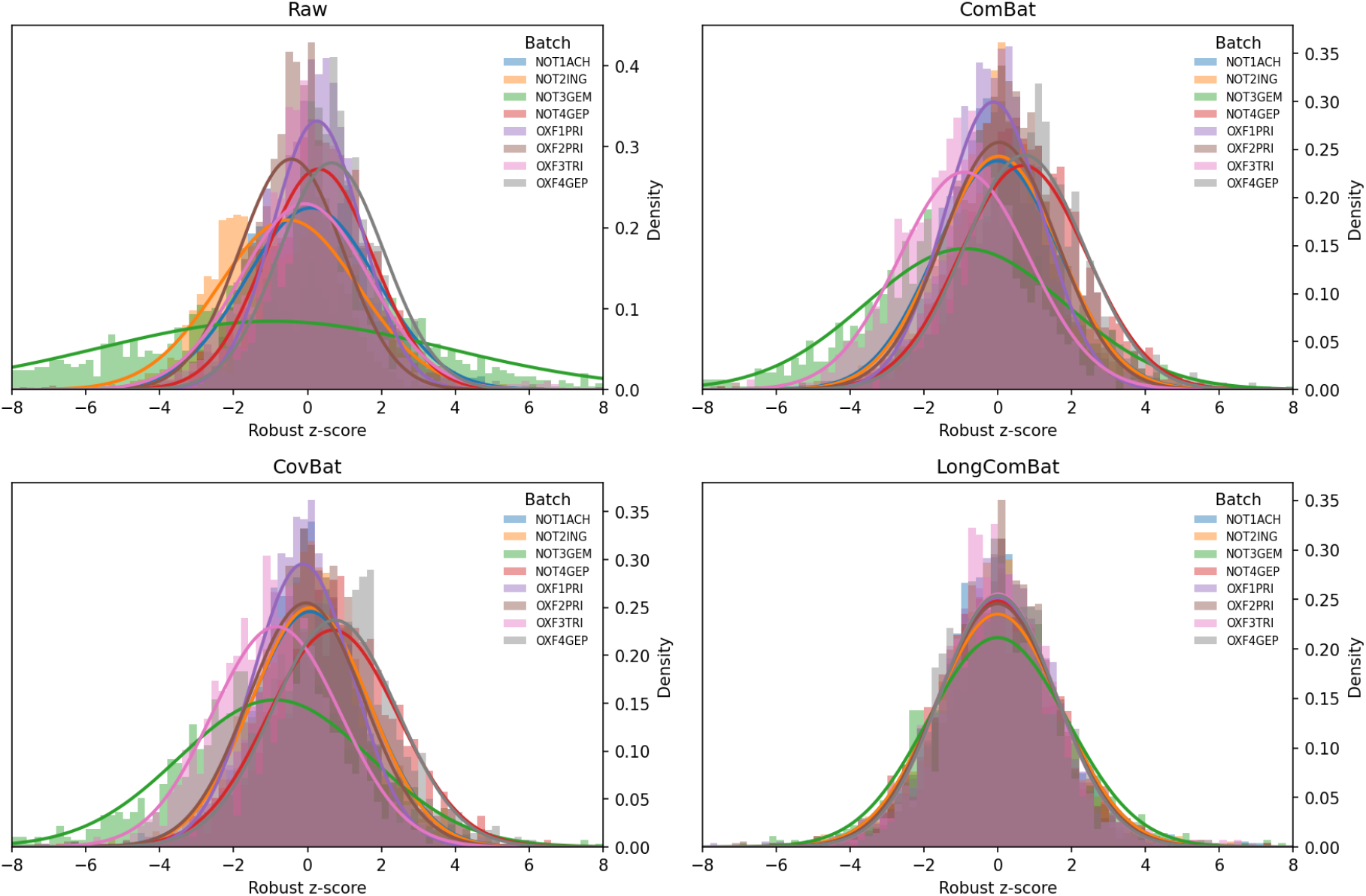
Robust Z-score distributions across scanners are shown for the raw data and following ComBat, CovBat, and LongComBat harmonisation. Histograms represent the density of pooled feature-wise robust Z-scores within each scanner, after accounting for covariate effects and subject-specific random intercepts. Greater overlap indicates greater similarity of feature distributions across scanners.

#### Within-subject variability and Subject order consistency

Within-subject relative percentage difference (RPD) is calculated for all pairs of repeated measurements. The RPD metric implemented in DHARM provides a general measure of pairwise within-subject variability across the repeated observations supplied by the user. To demonstrate how this output can be further interrogated according to the specific study design of ON-Harmony, RPD values from within-scanner repeats were additionally used as a baseline and compared with those from between-scanner repeats. This illustrates how the metric can be used to characterise different sources of measurement variability. For each feature, the median RPD across the relevant pairs was first calculated separately for each subject, followed by the median across subjects, resulting in one within-scanner and one between-scanner RPD value per feature.

As shown in Figure 14, before harmonisation, between-scanner variability in the raw data was greater than the baseline within-scanner variability, with a median RPD of 5.2% (IQR: 3.8–12.6%) compared with 1.8% (IQR: 1.3–3.2%) for within-scanner measurements. Harmonisation reduced the median and spread of between-scanner RPD across features, approaching the baseline within-scanner variability. Median between-scanner RPD was 4.3% (IQR: 3.4–5.8%) following ComBat, 4.4% (IQR: 3.5–5.7%) following CovBat, and 4.4% (IQR: 3.4–6.7%) following LongComBat.

**Figure 14:**
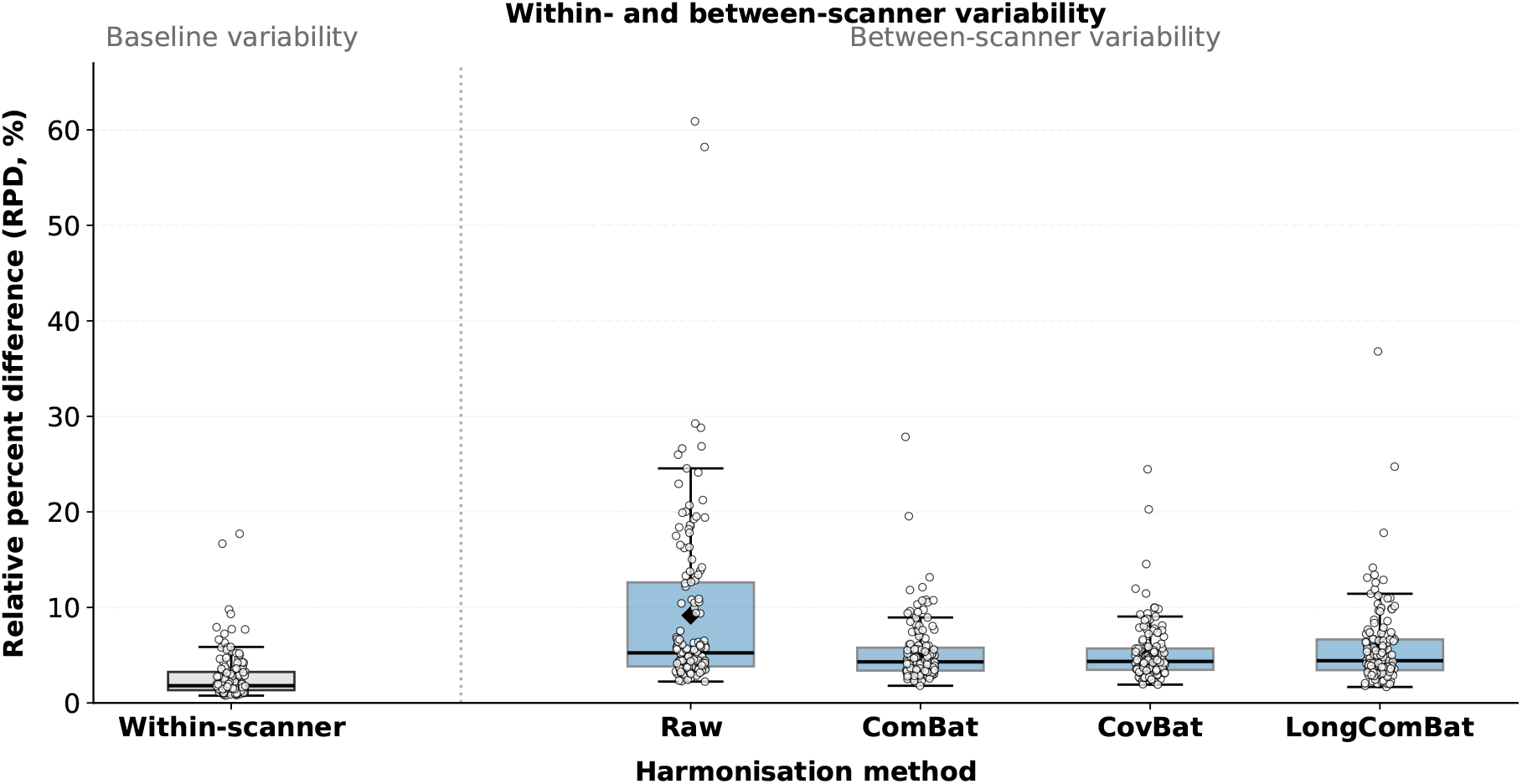
Relative percent difference (RPD) in atlas-based GM volumes for within-scanner group as baseline and between-scanner repeated measurements for the raw data and following ComBat, CovBat, and LongComBat harmonisation. Each point represents a feature, with RPD summarised first across measurement pairs within each participant and then across participants using the median.

For each feature, Spearman’s correlation coefficients were calculated across matched participants for each available pair of repeats and summarised using the median, resulting in one between-scanner subject-order consistency estimate per feature. As shown in Figure 15, subject-order consistency was high and broadly comparable across methods, with a median *ρ* of 0.9 (IQR: 0.8–0.9) for the raw data, ComBat, CovBat, and LongComBat. LongComBat nevertheless showed slightly lower correlations for some individual features, as reflected by the greater spread towards lower values.

**Figure 15:**
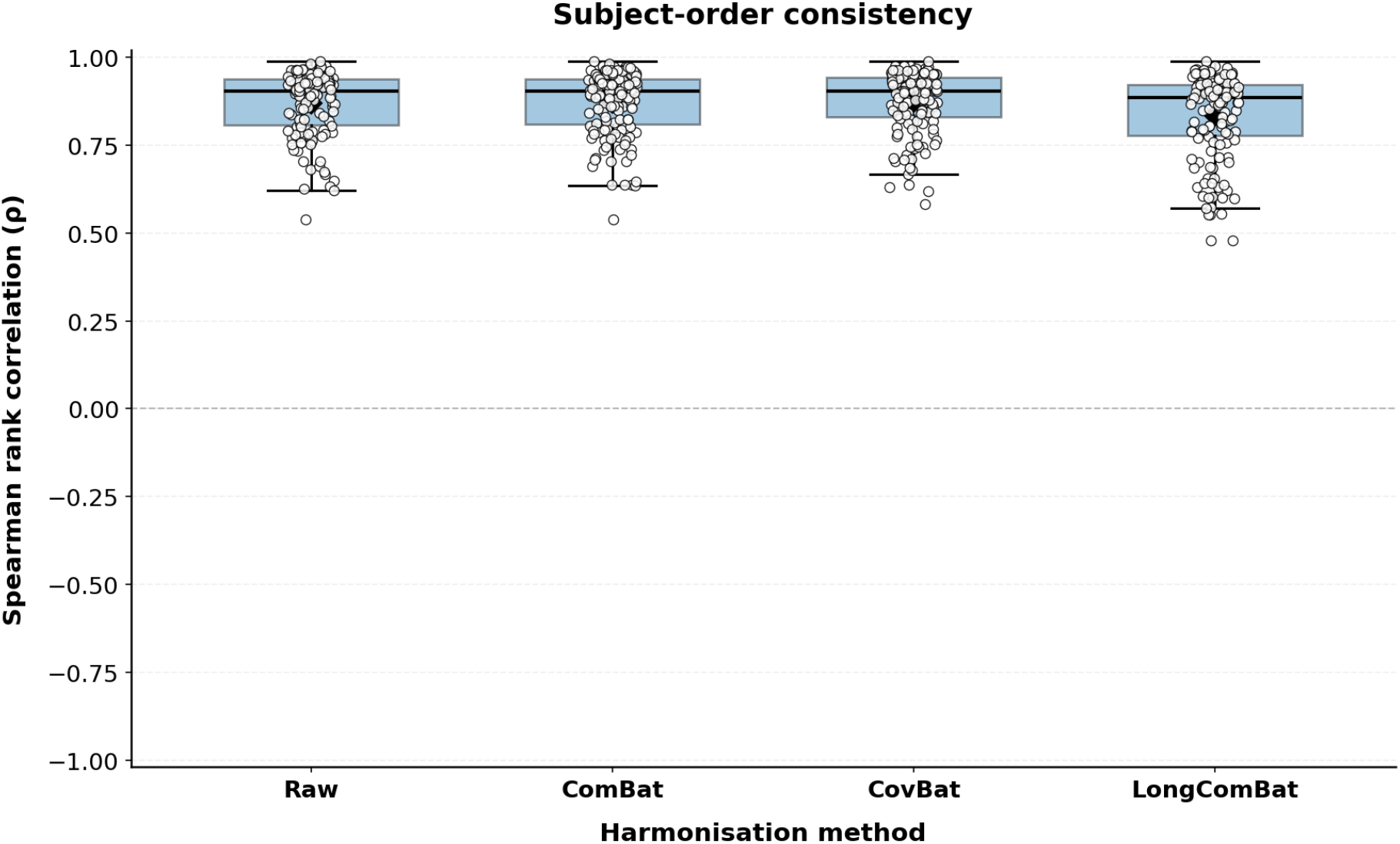
Between-scanner subject-order consistency before and after harmonisation. Spearman rank correlation (*ρ*) quantifies preservation of subject ordering of atlas-based GM volumes for between-scanner repeated measurements in the raw data and following Com-Bat, CovBat, and LongComBat harmonisation. Each point represents the median Spearman correlation for an individual feature.

#### Batch effect analysis (additive and multiplicative)

The batch-effect significance plots summarise additive (Figure 16) and multiplicative (Figure 17) batch effects as feature-wise plots of *−* log_10_(*p*_FDR_), where *p*_FDR_ denotes the FDR-corrected *p*-value. The dashed line indicates the significance threshold of *p*_FDR_ = 0.05, with features above this threshold showing a significant batch effect after FDR correction. Overall, the number of features showing significant additive and multiplicative batch effects was reduced following harmonisation. As shown in Figure 16, before harmonisation, significant additive batch effects were detected in 136 out of 138 features (98.6%). Following harmonisation, significant additive batch effects remained in 72/138 (52.2%) features following ComBat and 58/138 (42.0%) following CovBat. In contrast, no features showed a significant additive batch effect following LongComBat (0%). As shown in Figure 17, before harmonisation, significant multiplicative batch effects were detected in 25/138 (18.1%) features. Following harmonisation, significant multiplicative batch effects remained in 2/138 (1.4%) features following ComBat and 3/138 (2.2%) following CovBat. Following Long-ComBat, none of the features showed a significant multiplicative batch effect.

**Figure 16:**
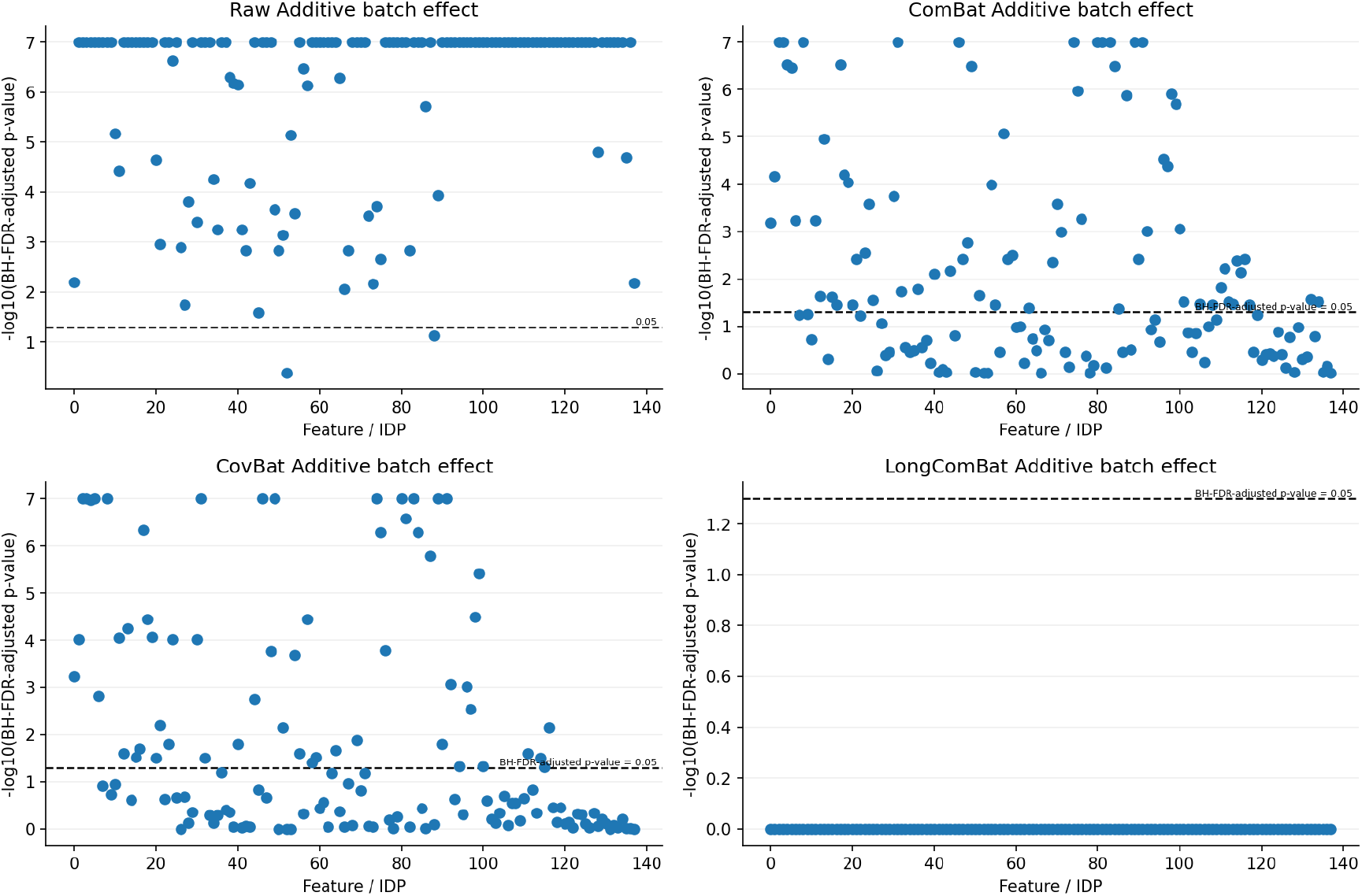
Additive batch effects before and after harmonisation. FDR-adjusted p-values are shown for raw data and following ComBat, CovBat, and LongComBat. Each point represents a feature (atlas-based GM volume), displayed as *−log*10(*p_F DR_*). Points above the dashed horizontal line (*p_F DR_* = 0.05) indicate a significant effect.

**Figure 17:**
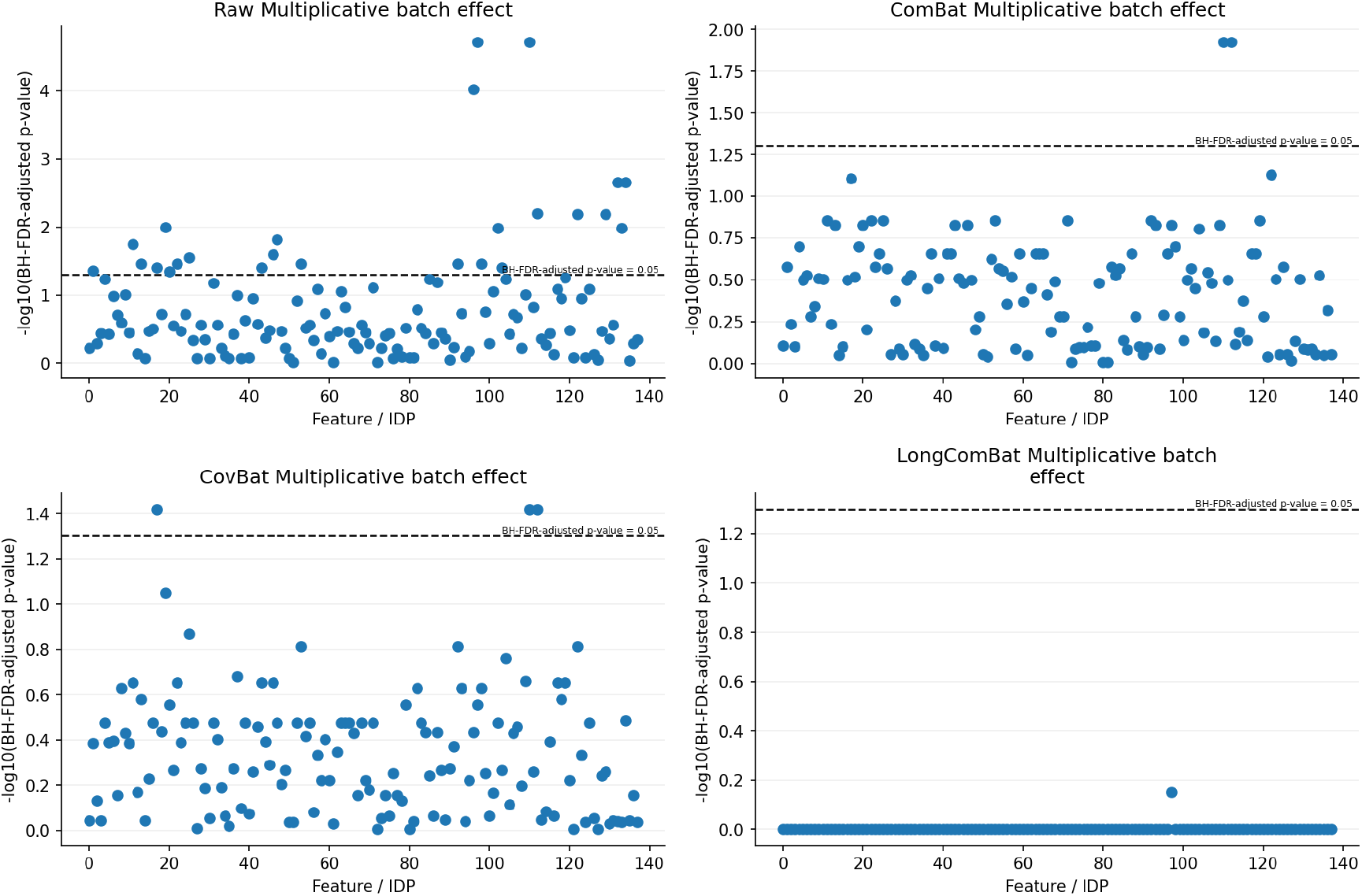
Multiplicative batch effects before and after harmonisation. FDR-adjusted p-values are shown for raw data and following ComBat, CovBat, and LongComBat. Each point represents a feature (atlas-based GM volume), displayed as *−log*10(*p_F DR_*). Points above the dashed horizontal line (*p_F DR_* = 0.05) indicate a significant effect.

As shown in Figure 17, before harmonisation, significant multiplicative batch effects were detected in 25/138 (18.1%) features. Following harmonisation, significant multiplicative batch effects remained in 2/138 (1.4%) features following ComBat and 3/138 (2.2%) following CovBat. Following LongComBat, none of the features showed a significant multiplicative batch effect.

#### Preservation of biology

Figure 18 shows subject intraclass correlation coefficients (ICC, calculated as ratio of between-subject variance and total variance) across features. Before harmonisation, the median ICC was 0.8 (IQR: 0.6–0.9). Following harmonisation, the median ICC increased to 0.9 (IQR: 0.8–0.9) for all the methods, indicating that a greater proportion of the remaining variance was attributable to stable between-subject differences rather than residual variability.

**Figure 18:**
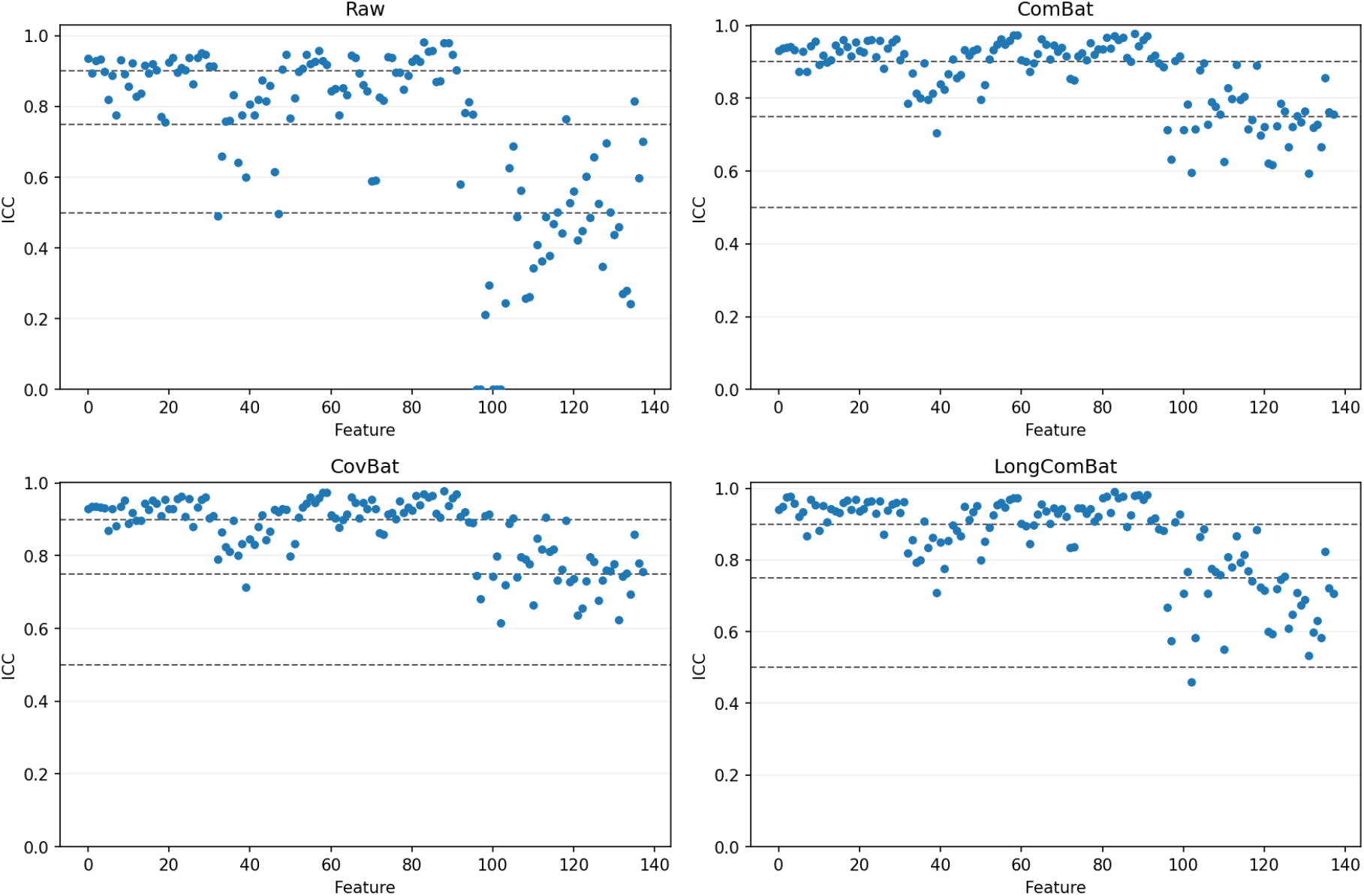
Subject Intraclass correlation coefficients (ICCs) are shown across atlas-based GM volumes (features) for the raw data and following ComBat, CovBat, and LongComBat harmonisation. Each point represents an individual feature. The relatively lower ICC values in the right of each subplot correspond to subcortical regions (*∼* 30% ) and from within the cerebellum (*∼* 50%).

Biological variability was assessed using feature-wise fixed effects estimated from the mixed-effects models. For each covariate, the tool reports the effect estimate (*β*), 95% confidence interval, and corresponding *p*-value for each feature. The *p*-values were additionally FDR-corrected across features for each covariate and *p*_FDR_ *<* 0.05 considered statistically significant.

As shown in Figure 19, age-related effect estimates showed a similar distribution across the raw and harmonised data, indicating that the overall pattern of age associations was largely preserved following harmonisation. After FDR correction, only 1 of 138 GM features (namely, the superior region of the lh-Lateral Occipital-Cortex) showed a significant negative age effect, consistently across the raw data, ComBat, CovBat, and LongComBat.

**Figure 19:**
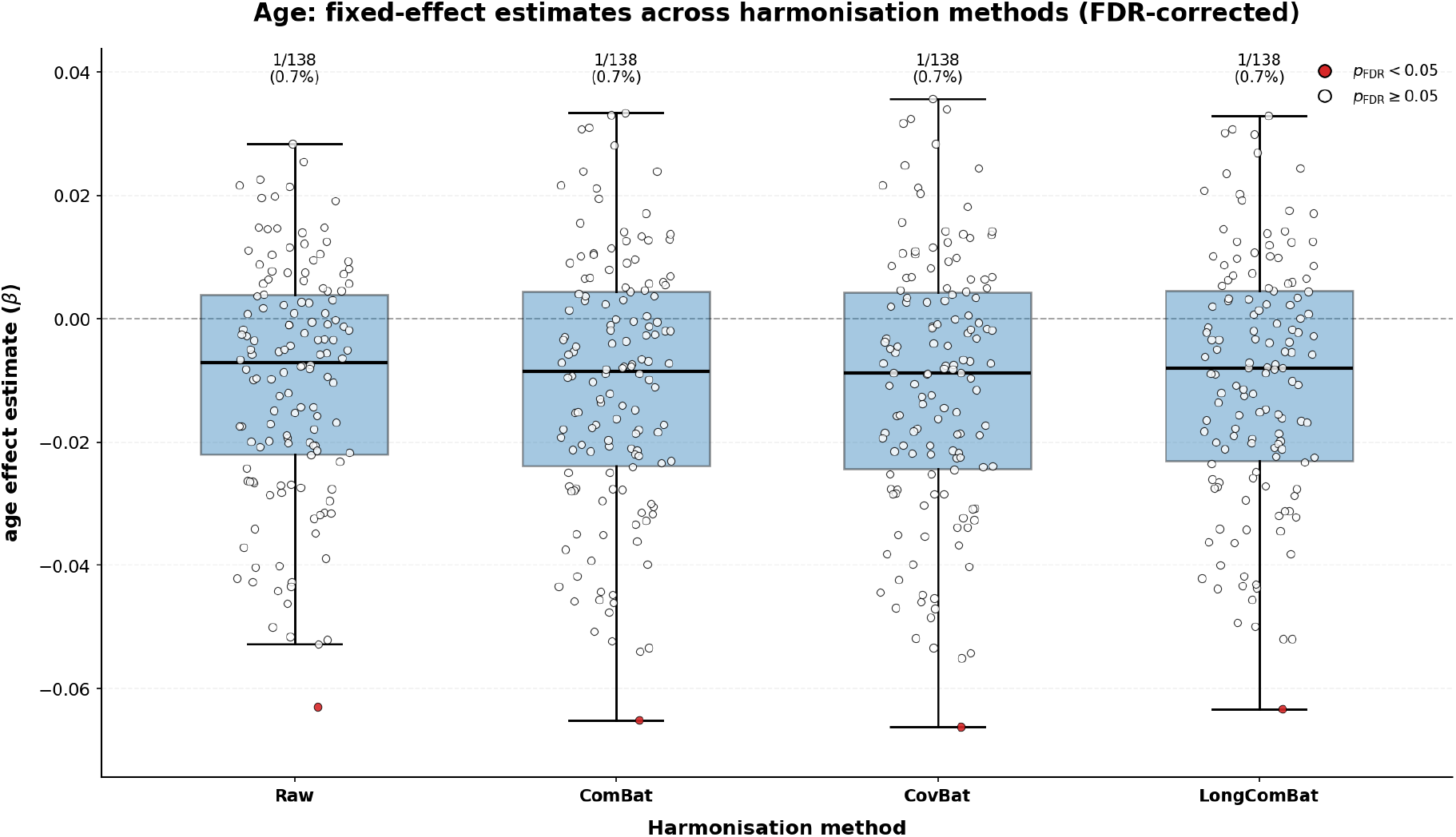
Age-related fixed-effect estimates before and after harmonisation. Age-related fixed-effect estimates (*β*) are shown across atlas-based GM volumes for the raw data and following ComBat, CovBat, and LongComBat harmonisation. Each point represents an individual feature, with filled red points indicating significant age associations (*p*_FDR_ *<* 0.05). The number and proportion of GM volumes with significant age associations are reported within each panel. All features were standardised, whereas age was retained in its original scale; therefore, *β* represents the change in the volume, in standard-deviation units, associated with a one-year increase in age.

## 4 Discussion

The development of Diagnose Harmonisation (DHARM) was motivated by two related challenges in the field: 1) the fragmented availability of harmonisation methods and 2) the lack of consistency in how batch effect and harmonisation performance are assessed and reported across multisite neuroimaging studies. Although a growing number of harmonisation methods and their implementations are available, these methods are not readily accessible within a common framework, and their evaluation is usually performed using different datasets, metrics, and criteria, limiting systematic comparison across studies and the subsequent selection of the most appropriate method for a given study. DHARM addresses these challenges by bringing a broad range of harmonisation methods and complementary diagnostic measures together within a single workflow. Rather than defining successful harmonisation by a single statistical test or visualisation, our framework evaluates changes in feature means, variance, empirical distributions, covariance, and multivariate structure, while also examining whether biologically meaningful variation has been retained.

The cross-sectional demonstrations illustrate why such multi-metric approaches are necessary. Across the cortical-thickness measures examined here, all tested harmonisation approaches reduced some of the most visually apparent differences between batches. Feature-wise Cohen’s d values were also substantially reduced following harmonisation, demonstrating attenuation of additive batch-associated shifts. However, the variance and distributional analyses showed that improvement in mean alignment did not necessarily imply equivalent correction of other forms of technical variation. In particular, differences remained between methods in their ability to align feature-specific scale and empirical distributions. A method may therefore appear effective when judged according to mean differences alone while retaining measurable multiplicative or distributional batch effects. This distinction is particularly important when comparing regression-based approaches, which primarily target additive differences, with methods such as ComBat that explicitly model both location and scale effects.

PCA provides a deterministic linear representation of the dominant directions of variation, but technical effects that are weak, non-linear, or distributed across lower-order components may not be obvious from inspection of the first two principal components. Conversely, visible separation of batches in a PCA representation may partly reflect biological differences if covariates are unevenly distributed between sites. UMAP is useful for examining non-linear local structure, but the appearance of an embedding depends on parameter choices and stochastic optimisation, and the orientation and absolute coordinates of an embedding have no intrinsic meaning. Consequently, these plots should primarily be used to identify relative clustering, gradients and neighbourhood structure, rather than as quantitative evidence in isolation. Their interpretation is strengthened when they are considered alongside feature-wise and model-based diagnostics, providing a complementary perspective. In the application shown, observations in the raw data showed visible organisation according to batch membership, which was generally reduced after harmonisation. At the same time, biologically relevant organisation, particularly that associated with age, improved after harmonisation. These findings illustrate an important principle underlying the DHARM framework: removal of detectable batch structure alone is not sufficient evidence of successful harmonisation. Methods should reduce technical differences while preserving or strengthening relationships attributable to biological covariates. The relative preservation of covariate-associated variance and multivariate biological structure in the analyses therefore provides information that would not be available from measures of residual batch association alone.

The repeated-measures demonstration further highlights that the interpretation of evaluation metrics should consider the underlying study design. This is particularly important for repeated-measures data, where technical batch variability can directly influence estimates of within-subject change. DHARM extends harmonisation evaluation by providing complementary metrics suited to different study designs. These include within-subject variability (via RPD) and subject-order consistency, which are particularly informative in test–retest and travelling-head studies, where no biological change is expected. Other metrics assess residual batch effects and preservation of biological associations, and can also be applied to longitudinal studies. We demonstrated these metrics using ON-Harmony, a multisite travelling-head dataset containing both within-and between-scanner repeated acquisitions. The RPD results showed that harmonisation reduced within-subject variability in the between-scanner repeats, bringing them closer to the within-scanner test–retest reference variability. Subject-order consistency was also largely maintained between the raw and harmonised data. Of note, the original ON-Harmony study (Warrington et al., 2023) used coefficient of variation (CoV) to assess within- and between-scanner variability. DHARM instead uses RPD as a pairwise measure of within-subject variability, allowing it to accommodate different numbers of repeated observations and repeated-measures structures, including subjects with only two observations. The reduced between-scanner variability and preservation of subject ordering were consistent with the findings of the original ON-Harmony study, while demonstrating how DHARM can characterise complementary aspects of measurement variability.

When evaluating different harmonisation methods on the ON-Harmony dataset, LongCom-Bat, which explicitly accounts for repeated observations through subject-specific random effects, showed no significant residual additive or multiplicative batch effects for the evaluated features (GM volumes). However, this finding should not be interpreted as independent evidence of successful harmonisation, as LongComBat explicitly estimates and removes these effects by design. Therefore, these diagnostics are of limited value for evaluating Long-ComBat’s effectiveness, although they remain useful for assessing residual batch effects following other harmonisation approaches. Importantly, LongComBat also showed small increases in within-subject variability and reduced subject-order consistency for some volumes relative to ComBat and CovBat. These findings illustrate that removing modelled batch effects does not necessarily translate into improved measurement consistency, highlighting the importance of complementary evaluation metrics that assess aspects of harmonisation performance beyond those directly targeted by the correction method.

A central objective of DHARM is therefore to provide a step towards a more unified and transparent harmonisation workflow rather than to prescribe a single definition of successful harmonisation. The same diagnostic framework can be applied to raw data and to the outputs of multiple correction approaches, allowing users to identify which aspects of batch variation are improved by each method and whether these improvements occur at the expense of biologically meaningful signal. This is particularly important because the optimal harmonisation strategy depends on the characteristics of the dataset and the intended downstream analyses. A method that performs well when site differences are predominantly additive may not be optimal when differences in scale or covariance dominate, while more flexible approaches may offer advantages when biological effects are non-linear or strongly confounded with batch. Harmonisation should therefore be considered a datasetand objective-specific problem rather than one for which a universally optimal method can be identified.

### Practical considerations

Several practical considerations of the present framework should also be recognised. First, the quality of the diagnostic conclusions depends on the information supplied to the model. If biologically important covariates are omitted, or if batch and biological variables are strongly confounded, technical and biological effects may be difficult or impossible to distinguish reliably. Apparent reduction of a batch effect in such circumstances may correspond partly to removal of genuine biological variation. Additionally, application of the repeated-measures metrics requires appropriate modelling of expected biological change and study-specific sources of variation. The current framework models batch, timepoint, and biological covariates (e.g., age and sex) as separate fixed effects, with subject as a random effect, providing a practical starting point for longitudinal assessment. More complex study designs may require adaptation to incorporate interactions among these factors and more flexible subject-specific trajectories, depending on the sampling design and expected biological dynamics. Careful inspection of covariate distributions and consideration of the acquisition context remain essential before applying and interpreting any harmonisation method.

Second, DHARM is designed primarily for tabular imaging-derived phenotypes rather than direct manipulation of the underlying images. Consequently, spatially localised image artefacts or acquisition differences that do not propagate strongly into the selected derived measures may not be identified by the current diagnostics. This does not prevent DHARM from being used to evaluate image-level harmonisation methods: derived phenotypes produced before and after an image-level correction can still be passed through the diagnostic workflow to determine whether downstream batch effects have been reduced and biological associations preserved. Nevertheless, direct evaluation of image appearance, spatial anatomy and image quality remains complementary to the feature-level assessments provided here.

Finally, statistical tests included within the framework remain sensitive to sample size and data dimensionality. Very large datasets may produce statistically significant differences that are negligible in practical magnitude, whereas small batches may provide insufficient power to identify important technical effects. Covariance-based and multivariate measures can additionally become unstable when the dimensionality of the feature space approaches or exceeds the number of available observations. For this reason, the report deliberately combines statistical significance with effect-size measures, visualisations and model-based quantities. Interpretation should focus on the overall pattern of evidence and the practical magnitude of remaining differences rather than on whether individual p-values cross a fixed significance threshold.

## 5 Conclusion

In this manuscript, we introduce DHARM, a toolbox for diagnosing and characterising batch effects in a dataset, applying harmonisation methods, and assessing harmonisation performance. Rather than proposing a single metric for detecting batch effects, it integrates multiple complementary perspectives within one reproducible diagnostic workflow. The analyses described in this paper illustrate the importance of this approach: harmonisation can substantially reduce batch effects in one type of measure while producing different results in another. DHARM provides a practical framework for selecting and validating harmonisation strategies according to the characteristics of the datasets and scientific objectives of the study. This approach moves harmonisation assessment away from reliance on a single measure of success and towards a more transparent, evidence-based evaluation of whether technical differences have been reduced without compromising the biological information contained within the data.

## Supporting information

Full Supplemental Materials

## Authors’ Contributions

**Conceptualisation:** Jacob Turnbull, Gaurav Bhalerao, Stephen M. Smith, Ludovica Griffanti

**Data Curation:** Jacob Turnbull, Gaurav Bhalerao, Shaun Warrington, Stamatios N. Sotiropoulos, Fidel Alfaro-Almagro, Grace Gillis, Brian Zhaoyi Mo, Stephen M. Smith, Ludovica Griffanti

**Formal analysis:** Jacob Turnbull, Gaurav Bhalerao

**Funding acquisition:** Stephen M. Smith, Ludovica Griffanti

**Methodology:** Jacob Turnbull, Gaurav Bhalerao, Thomas E. Nichols, Stephen M. Smith, Ludovica Griffanti

**Project administration:** Jacob Turnbull, Gaurav Bhalerao, Ludovica Griffanti

**Resources:** Shaun Warrington, Stamatios N. Sotiropoulos, Fidel Alfaro-Almagro, Grace Gillis, Jasmine Blane, Brian Zhaoyi Mo

**Software:** Jacob Turnbull, Gaurav Bhalerao, Paul McCarthy, Renato Besenczi

**Supervision:** Stephen M. Smith, Ludovica Griffanti

**Visualisation:** Jacob Turnbull, Gaurav Bhalerao

**Writing – original draft:** Jacob Turnbull, Gaurav Bhalerao

Writing – review & editing: All authors

## Use of AI

AI and/or other tools were used to check spelling and grammar. AI was used in some parts to reword draft text. AI was not used to generate any original text for this paper. AI was not used in generating any figures. ChatGPT 5.5 and 5.6-sol were used under the educational agreement between the University of Oxford and OpenAI.

## Ethics

This study involved secondary analysis of existing datasets obtained through the relevant data access procedures. Each dataset was originally collected under the applicable ethical approvals and governance procedures; further details regarding ethical approval and informed consent for the original data collection are provided in the corresponding publications cited in the Methods. No new participant data were collected as part of the present study, and no additional ethical approval was required for the secondary analyses reported here.

This research was conducted in part using data from UK Biobank under approved application number 8107. UK Biobank has ethical approval from the Northwest Multicentre Research Ethics Committee as a research tissue bank, and all participants provided informed consent for their de-identified data to be used for health-related research in the public interest. This study involved secondary analysis of de-identified UK Biobank data and was conducted in accordance with the UK Biobank Access Procedures.

The present study involved secondary analysis of anonymised, openly available data from the ON-Harmony dataset (OpenNeuro accession ds004712, version 2.0.1). The dataset was collected in two phases. For Phase A, data collection at the University of Nottingham was conducted under ethical approvals FMHS-36-1220-03 and H14082014/47, while technical development at the University of Oxford was approved by the Oxford University Clinical Trials and Research Governance office. For Phase B, data collection was approved by the University of Nottingham Research Ethics Committee (FMHS-452-0122) and the University of Oxford Medical Sciences Research Ethics Committee (R79669/RE001).

## Acknowledgements

We are grateful to all participants in the studies included in this work who generously donated their time and contributed their data to make these research resources possible. This research has been conducted in part using the UK Biobank Resource under Application Number 8107. We are grateful to Sameera Shabir and Luciana Maffei for their major contribution to the data collection of the OxBAC dataset. We are grateful to the members of OxCIN’s (Oxford University Centre for Integrative Neuroimaging) Harmonisation Working Group for the helpful discussions and feedback on the tool.

## Data and code availability statement

UK Biobank data used in this study are available to approved researchers through the UK Biobank Access Management System, subject to UK Biobank’s standard access procedures.

Cam-CAN data used in this study are available through the Cam-CAN repository: https://cam-can.mrc-cbu.cam.ac.uk/dataset/

OxBAC data used in this study are available to approved researchers upon request to the Oxford Brain Health Clinic (OBHC, https://www.psych.ox.ac.uk/research/research-groups/translational-neuroimaging-group/team/oxford-brain-health-clinic), subject to an application process, review by the OBHC Data Access Group, and completion of the relevant data sharing agreement.

The ON-Harmony data used in this study are openly available through OpenNeuro (dataset ds004712, version 2.0.1; https://openneuro.org/datasets/ds004712/versions/2.0.1).

The DHARM tool is available on GitHub: https://github.com/Jake-Turnbull/HarmonisationDiagnostics

