## Supplementary material for "Diagnose Harmonisation (DHARM): An open-source tool for the harmonisation and analysis of batch effects in MRI-derived measures": Full Supplemental Materials

#### 1 Preamble

These supplementary materials accompany the corresponding manuscript "Diagnose Harmonisation (DHARM): An open-source tool for the harmonisation and analysis of batch effects in MRI-derived measures".

They provide additional methodological and practical information supporting the analyses presented in the manuscript. First, we describe the mathematical background and implementation of the diagnostic metrics available within DHARM, including their assumptions and considerations for interpretation. We also provide an overview of the harmonisation methods implemented within the framework. Detailed instructions for using DHARM and its associated diagnostic and harmonisation functions are available in the Online Documentation. The remainder of the supplement provides additional information for the cross-sectional and longitudinal demonstrations presented in the manuscript. This includes details of the datasets and imaging-derived phenotypes (IDPs) used, the code and configurations used for harmonisation and report generation, and the complete diagnostic reports generated by DHARM. These reports are included to provide a transparent record of the tool outputs underlying the results and figures presented in the manuscript.

#### 2 Supplementary Methods: Diagnostic Metrics

##### 2.1 Notation and covariate-adjustment

The general form of the covariate-adjusted data is given by equations 1b and 1d and is used frequently throughout this manuscript. The full equations and descriptions for all methods of analysis are available in this supplement. The unadjusted distributions represent the total observed variation in the imaging features and therefore reflect the combined influence of technical batch effects, biological covariates, covariate imbalance, and residual noise. The covariate-adjusted distributions are designed to assess whether systematic batch differences remain after accounting for measured covariates. Because batch and covariates may be partially collinear in multi-site neuroimaging studies, batch is included in the model used to estimate covariate effects; however, here, the fitted batch contribution is retained rather than removed.

In each case, the fitted covariates are automatically sorted into categorical, binary and continuous depending on the number of unique observations relative to sample size. In the case of categorical covariates, these variables are dummy expanded into $N_{cat} - 1$ columns within the covariate matrix $C_{i,c}$.

$$X_{i,c+b} = [1 \quad C_{i,c} \quad B_i] \in \mathbb{R}^{n \times (k+1+n_b)} \tag{1a}$$

$$Y_{i,f} = X_{i,c+b}\hat{\beta}_{f,c+b} + \epsilon_{i,f} \tag{1b}$$

$$\hat{\beta}_{f,c+b} = (X_{i,c+b}^{\top} X_{i,c+b})^{-1} X_{i,c+b}^{\top} y_{i,f} \tag{1c}$$

$$y_{i,f}^{cov-adj} = Y_{i,f} - C_{i,c}\hat{\beta}_{f,c} \tag{1d}$$

##### 2.2 Histogram comparisons

We compare histograms of the Median Absolute Deviations (MADs) of all features. We present these in two ways: one for covariate-unadjusted data and one for covariate-adjusted data. Batch differences in the covariate-unadjusted data may be simply due to different covariate distributions, therefore, we show both the adjusted and unadjusted data for clarity. The equation for the resulting data after converting to MADs is given by equation 2a, the form of the covariate-adjusted data, which then substitutes for $y_{if}$ in equation 2a is given by equation 2b.

$$MAD_f = median(|x_{if} - median_f(x_{if})|), \tag{2a}$$

$$y_{ij}^{MAD's} = \frac{(y_{ij} - median_f(x_{if}))}{1.4826 \times MAD_f} \tag{2b}$$

##### 2.3 Cohen's d

Additionally, as different batches are likely to have different covariate distributions (i.e., different ages between batches of data), we first convert the data to covariate-adjusted data, as shown in equation 1d. Using the residuals, we then calculate the feature-wise Cohen's d effect size for an individual batch against all other batches, as given by equation 3, where $\mu_b$ is the mean and $\sigma_b$ the standard deviation of the data denoted by $b$.

$$d = \frac{\mu_{batch=b} - \mu_{batch \neq b}}{\sqrt{(1/2)(\sigma^2_{batch=b} + \sigma^2_{batch \neq b})}} \tag{3}$$

##### 2.4 Mahalanobis distance

Small feature-wise shifts that appear modest individually may combine to produce substantial (or at least statistically significant) multivariate separation between batches. Mahalanobis distance captures this joint displacement by scaling differences in mean vectors by the covariance structure of the data.

Like Cohen's d, we also calculate Mahalanobis distance on the covariate-adjusted data, showing the Mahalanobis distance between batches, as well as between each batch and the global centroid of the data for both the adjusted and unadjusted measures. For the longitudinal workflow, the Mahalanobis distance framework is adapted to account for the repeated-measures structure of the data, with the specific implementation determined by the within-subject batch configuration.

Let $\mu_b$ denote the mean feature vector of batch b and $\mu_{ref}$ be the mean vector of a second batch, or the mean vector of the entire dataset excluding the batch $b$. Finally, let $\Sigma$ be the pooled covariance matrix of all data used in calculating $\mu_b$ and $\mu_{ref}$. The Mahalanobis distance for batch b is then defined as

$$D_M(b, ref) = \sqrt{(\mu_b - \mu_{ref})^{\top} \Sigma^{-1} (\mu_b - \mu_{ref})}$$

We include the Mahalanobis distance to assess whether batches differed in their multivariate location after accounting for the covariance among imaging features. This provides a global measure of batch separation in the joint feature space, complementing univariate effect-size estimates from Cohen's d. Like Cohen's d, we also calculate Mahalanobis distance on the covariate-adjusted data, showing the Mahalanobis distance between batches, as well as between each batch and the global centroid of the data for both the adjusted and unadjusted measures. This provides insight into the unique contributions to the effect on the mean by batch. In some cases, the batches could get further away from each other or from the global centroid after removing covariate effects, making this a useful sanity check of batch effects separate from population differences.

However, Mahalanobis distance can be sensitive to covariance estimation, high dimensionality, and collinearity among features. Additionally, it lacks the directionality of the difference in means that Cohen's d provides. It should therefore be interpreted alongside univariate metrics and covariance-structure diagnostics. Differences between the cross-sectional and longitudinal implementations of the Mahalanobis distance are described in the manuscript.

#### 2.5 Model fit comparison using linear modelling

Three different linear models are fitted, each answering a different question. How much variation is explained by the measured covariates? How much additional variation is explained by batch after accounting for covariates? How strongly are observations clustered within batches? The three models are given, respectively, by equations 4a, 4b and 4c.

$$y_{if} = \beta_0 + X_{cov}\beta_{cov} + \epsilon \tag{4a}$$

$$y_{if} = \beta_0 + X_{cov}\beta_{cov} + X_{batch}\beta_{batch} + \epsilon \tag{4b}$$

$$y_{if} = \beta_0 + X_{cov}\beta_{cov} + u_{batch} + \epsilon \tag{4c}$$

Complementary statistics are calculated and reported, $R^2_{cov}$, $R^2_{cov+batch}$, $\Delta R^2_{batch}$ and the feature-wise ICC. The feature-specific $R^2$ for a given model is calculated as given in Equation 5a and Equation 5b, with the difference attributable to batch being given by Equation 5c

$$R^2_{cov} = 1 - \frac{RSS_{cov}}{TSS} \tag{5a}$$

$$R^2_{cov+batch} = 1 - \frac{RSS_{cov+batch}}{TSS} \tag{5b}$$

$$\Delta R^2_{batch} = R^2_{cov+batch} - R^2_{cov} \tag{5c}$$

The random-intercept mixed-effects model is used to estimate the batch Intraclass Correlation Coefficient (batch-ICC):

$$ICC = \frac{\sigma^2_{batch}}{\sigma^2_{batch} + \sigma^2_{residual}} \tag{6}$$

Where $\sigma^2_{batch}$ is the estimated between-batch variance and $\sigma^2_{residual}$ is the residual variance. The batch-ICC estimates the proportion of unexplained variance attributable to clustering within batches, with larger values indicating stronger batch-dependent clustering.

For completeness, the marginal and conditional coefficients of determination for the mixed-effects model may also be reported. These describe the proportion of variance explained by the fixed effects alone and by both the fixed and random effects combined, respectively, but are presented as supplementary mixed-model variance decomposition measures rather than the primary batch-effect statistics.

Finally, to assess whether inclusion of the batch random intercept significantly improves model fit, the mixed-effects model is compared with the corresponding covariate-only fixed-effects regression using a likelihood-ratio test. Likelihood-ratio statistics and associated p-values are calculated for each feature using maximum-likelihood model fits. Multiple optimisation algorithms are attempted sequentially to improve robustness (Lindstrom & Bates, 1988; Seabold et al., 2010). If batch sizes fall below a user-specified threshold or the mixed-effects model fails to converge after all optimisation attempts, only the fixed-effects models are retained and mixed-model statistics are reported as unavailable.

#### 2.6 The F-test of variance differences

This is performed on the covariate-adjusted data as opposed to the unadjusted data to ensure minimal contribution from factors other than batch. The data for the result of the F-test are displayed as a feature-wise estimate $V_r$, with the calculation being shown in equation 7b, where $\sigma_{batch=b}$ is the standard deviation of batch $b$.

$$\sigma^2_{batch=b} = \frac{\sum_{i=1}(y_i - \bar{y}_{batch=b})^2}{N_b} \tag{7a}$$

$$V_r = \frac{\sigma^2_{batch=b}}{\sigma^2_{batch\neq b}} \tag{7b}$$

#### 2.7 Fligner-Killeen test of homogeneity

The tool offers three different alternatives for testing the homogeneity of variance between batches: Levene's test, the Brown-Forsythe test and the Fligner-Killeen test. The form of each test is shown in equation 8, with the only variation between them being the transformed response variables $z_{ib}$ for participant $i$ in the batch $b$. For Levene's test, $z_{ib}$ is obtained by subtracting the mean of batch $b$. For the Brown-Forsythe and Fligner-Killeen tests, this is instead done by subtracting the median. For the Fligner-Killeen test, the response variables are then additionally ranked and the deviations of the ranks and not the transformed variable used (Brown & Forsythe, 1974).

$$W = \frac{N-p}{p-1} \frac{\sum_{b=1}^{p} n_b(z_{.b} - \tilde{z}_{..})^2}{\sum_{j=1}^{p}\sum_{i=1}^{n_b}(z_{ib} - \tilde{z}_{.b})^2} \tag{8}$$

The Fligner-Killeen test is applied to each feature and unique batch combination independently for both the raw data and the residuals after accounting for covariate effects. As such, the form of equation 8 has degrees of freedom $d_1 = 1$ and $d_2 = N - 2$ where $N$ is the total number of samples in both batches for a given comparison. This test can also be applied to all batches at once, which, in cases where batches are small or when total sample heterogeneity is considered more important than group-wise comparisons, may be beneficial over applying the test between batch pairs.

#### 2.8 The two-sample Kolmogorov-Smirnov test of empirical distributions

The two-sample KS test is used to calculate the difference between the empirical distributions between two batches using a supremum function across the entire distribution for both batches, taking the maximum value as given in Equation 9a.

$$D_{m,n} = max_x|F(x) - G(x)| \tag{9a}$$

$$D_{m,n} > c(\alpha)\sqrt{\frac{m+n}{m \cdot n}} \tag{9b}$$

#### 2.9 Principal Component Analysis

For a given feature matrix $X$, PCA decomposes the covariance matrix as shown in equation 10 with subject-specific score $z_k = Xv_k$. Here, $V$ contains the principal loading vectors, $\Lambda$ the diagonal matrix of eigenvalues representing explained variance.

$$X^\top X = V\Lambda V^\top \tag{10}$$

#### 2.10 Cluster analysis and principal component variance spectra

To further characterise batch-related structure differences in the multivariate feature space, we compute the variance captured by each principal component within each batch. Here, let $\mathbf{Z}$ be the principal component matrix of real values and let $Z_{ij}$ be the score of participant i in principal component j. For each unique batch, $b$, the sample variance along component $j$ is calculated as

$$s_{bj}^2 = \frac{1}{n_b - 1} \sum_{i \in b} (Z_{ij} - \bar{Z}_{bj})^2 \tag{11}$$

Here, $n_b$ is the number of participants in batch $b$ and $\bar{Z}_{bj}$ is the mean score for batch $b$ of component $j$. The proportion of within-batch variance explained by each component is then defined $f_{bj}$ and the cumulative fraction of variance explained up to component $m$ as shown in equation 12

$$f_{bj} = \frac{s_{bj}^2}{\sum_{l=1}^{k} s_{bl}^2}, F_{bm} = \sum_{j=1}^{m} f_{bj} \tag{12}$$

These per-batch scree and cumulative variance curves provide a compact summary of how dispersion is distributed across principal components within each batch. Differences in these curves between batches indicate that technical effects may alter the dominant directions of variation, even when the overall means appear similar.

#### 2.11 Covariance structure difference using the Frobenius norm

To assess whether batches differ in their covariance structure, a covariance matrix was estimated separately for each batch using the original feature matrix. For batch b, the covariance matrix is given by $\Sigma_b = Cov\mathbf{Y}_b$, where $\mathbf{Y}_b$ denotes the subset of observations in batch $b$. Pairwise differences between batch covariances can then be quantified by using the Frobenius/Euclidean norm between the matrices in each case:

$$D_{bc} = ||\Sigma_b - \Sigma_c||_F = \sqrt{\sum_{i=1}^{p} \sum_{j=1}^{p} (\Sigma_{b,ij} - \Sigma_{c,ij})^2} \tag{13}$$

Here, $p$ is the total number of features. The Frobenius norm provides a single summary metric of the overall difference between two covariance matrices, incorporating changes in both variance and correlations. To facilitate the comparison of each batch with each other, we normalise the result by the Frobenius norm of the pooled covariance matrix,

$$D_{bc}^* = \frac{||\Sigma_b - \Sigma_c||_F}{||\Sigma_{pooled}||F} \tag{14}$$

The resulting pairwise distance matrix is visualised as a heatmap, together with summary statistics describing the maximum covariance difference associated with each batch.

#### 2.12 Cluster analysis using UMAP

A full explanation of the underlying topological assumptions and derivation of the algorithm can be found in McInnes et al. (2020). Given observations $x_i$ and $x_j$, edge weights are computed from local connectivity estimates and represented as a fuzzy simplicial set. The embedding coordinates $y_i$ are then obtained by minimising the cross-entropy objective.

$$C = \sum_{i \neq j} [w_{ij} \log(\frac{w_{ij}}{\hat{w}ij}) + (1 - w_{ij}) \log(\frac{1 - w_{ij}}{\hat{w}_i j})] \tag{15}$$

Here, $w_{ij}$ and $\hat{w}_{ij}$ denote graph edge weights in the original and embedded space, respectively.

Several parameters influence the resulting embedding, including the minimum distance between embedded points ($min_{dist}$), the number of neighbours ($n_{neighbours}$), the number of embedding dimensions, and the distance metric used to construct the neighbourhood graph. By default, embeddings are generated in two dimensions to facilitate visualisation, and Euclidean distance is used as the similarity metric. The default settings for UMAP are 15 neighbours and a minimum distance of 0.1, with a random seed of 42 for reproducibility. This can be changed by the user and is explained further in the online documentation.

The $n_{neighbors}$ parameter controls the balance between local and global structure preservation, while $min_{dist}$ determines how tightly observations may be packed together in the low-dimensional representation. Consequently, different parameter choices can emphasise different aspects of the data structure. Additionally, UMAP is a stochastic dimensionality reduction method, and repeated runs may produce slightly different embeddings. Furthermore, the absolute coordinates, orientation, and scale of a UMAP embedding are arbitrary; interpretation should therefore focus on relative neighbourhood relationships and clustering patterns rather than the precise location of points in the embedded space.

When optimisation of UMAP parameters is desired, an efficient strategy is to evaluate multiple combinations of $n_{neighbours}$ and $min_{dist}$ on representative subsets of the data while preserving the relative proportions of each batch. For each parameter combination, clustering methods such as k-means may be applied to the resulting embedding, and agreement between cluster assignments and known batch labels can be quantified using measures such as the Adjusted Rand Index (ARI) or Normalised Mutual Information (NMI). Parameter values producing the most consistent recovery of the known batch membership structure can then be applied to the full dataset.

#### 2.13 Longitudinal, test-retest metrics

Full descriptions of the test-retest-specific metrics can be found in Bhalerao et al. (2026). Additionally, we provide some brief descriptions below:

##### 2.13.1 Within-Subject Variability

To quantify within-subject variability, we calculated the Relative Percent Difference (RPD) for each feature between pairs of measurements from the same subject acquired at different sites. Lower RPD values indicate greater measurement consistency and reduced site-related variability. For each data point and feature, RPD was calculated as:

$$\text{RPD} = \frac{|x_1 - x_2|}{(x_1 + x_2)/2} \times 100 \tag{16}$$

where $x_1$ and $x_2$ denote the feature values for the same subject at the two sites.

##### 2.13.2 Subject order consistency

In test-retest settings, given the relatively short acquisition interval, substantial changes in the relative ordering of subjects were considered unlikely. We therefore assess subject ranking consistency using Spearman's rank correlation ($\rho$) between each acquisition pair for each feature (Warrington et al., 2023). Higher $\rho$ indicates greater preservation of subject ranking across pairs of sites. Statistical significance was assessed using permutation tests. For each feature and acquisition pair, a null distribution of Spearman's rank correlation coefficients was generated by randomly permuting subject labels at one site ($n$ = 10,000) times while keeping the subject ordering at the other site fixed. The permutation $p$-value was calculated as:

$$p = \frac{1 + \sum_{i=1}^{N} \mathbb{I}\left(\rho_i^{\text{perm}} \geq \rho_{\text{obs}}\right)}{N + 1}, \tag{17}$$

where $\rho_{\text{obs}}$ is the observed Spearman correlation, $\rho_i^{\text{perm}}$ is the correlation from permutation $i$, and $N$ = 10,000. A significant result ($p < 0.05$) indicates greater rank consistency than expected under random subject pairing.

###### 2.13.3 Additive batch effect

Additive batch effects were assessed for each feature using linear mixed-effects models. The full model included age, sex, and batch (site) as fixed effects, with a subject-specific random intercept to account for repeated measurements:

$$Y_i = \beta_0 + \beta_{age} Age_{ij} + \beta_{sex} Sex_{ij} + \beta_{batch} Batch_{ij} + u_i + \epsilon_{ij}, \tag{18}$$

where $Y_{ij}$ is the feature for subject $i$ at observation $j$, $\beta_0$ is the fixed intercept, $\beta_{age}$, $\beta_{sex}$, and $\beta_{batch}$ are the fixed-effect coefficients for age and batch, respectively, $u_i$ is the subject-specific random intercept, and $\epsilon_{ij}$ is the residual error.

The full model was compared with a reduced model excluding the batch effect:

$$Y_{ij} = \beta_0 + \beta_{age} Age_{ij} + \beta_{sex} Sex_{ij} + u_i + \epsilon_{ij}. \tag{19}$$

The models were compared using a Kenward-Roger F-test, providing an omnibus test of whether batch explained additional variation in the feature beyond the covariates (e.g., age, sex) in the above case.

###### 2.13.4 Multiplicative batch effects

Multiplicative batch effects were assessed for each feature using a linear mixed-effects model including covariates (e.g., age, sex) and batch (site) as fixed effects, with a subject-specific random intercept:

$$Y_{ij} = \beta_0 + \beta_{age} Age_{ij} + \beta_{sex} Sex_{ij} + \beta_{batch} Batch_{ij} + u_i + \epsilon_{ij}, \tag{20}$$

where $Y_{ij}$ is the feature for subject $i$ at observation $j$, $\beta_0$ is the fixed intercept, $\beta_{age}$, $\beta_{sex}$, and $\beta_{batch}$ are the fixed-effect coefficients for age, sex, and batch, respectively, $u_i$ is the subject-specific random intercept, and $\epsilon_{ij}$ is the residual error.

Residuals from the fitted model were tested for heterogeneity of variance across batches using the Fligner-Killeen test. This provides a test of whether residual variability differs systematically between batches, indicating a multiplicative batch effect.

###### 2.13.5 Site-pairwise batch effects

To identify site-specific batch effects that may be obscured by the omnibus test (additive batch effect), we assess pairwise comparisons across the sites/batches. For each feature, we fit a linear mixed-effects model. For example, if age and timepoint were provided as biological covariates:

$$Y_{ij} = \beta_0 + \beta_{\text{age}} \text{Age}_{ij} + \beta_{\text{sex}} \text{Sex}_{ij} + \beta_{\text{batch}} \text{Batch}_{ij} + u_i + \epsilon_{ij}, \tag{21}$$

where $Y_{ij}$ is the raw or harmonised IDP for subject $i$ at observation $j$; $\beta_0$ is the fixed intercept; $\beta_{\text{age}}$and $\beta_{\text{batch}}$ are the fixed-effect coefficients for age, timepoint, and batch, respectively; $\text{Age}_{ij}$, $\text{Time}_{ij}$, and $\text{Batch}_{ij}$ denote the corresponding covariates; $u_i$ is the subject-specific random intercept; and $\epsilon_{ij}$ is the residual error.

Pairwise site differences were assessed using linear contrasts of the estimated site effects. Statistical significance was determined using Bonferroni correction across the 28 pairwise site comparisons for each feature (adjusted $p < 0.05$).

###### 2.13.6 Between-subject variability

To assess preservation of between-subject variability following harmonisation, we fitted a linear mixed-effects model for each raw and harmonised feature, with biological covariates as fixed effects and a subject-specific random intercept:

$$Y_{ij} = \mu + u_i + \epsilon_{ij}, \tag{22}$$

where $\mu$ is the fixed intercept, $u_i \sim \mathcal{N}(0, \sigma^2_{\text{subject}})$ is the subject-specific random intercept with variance $\sigma^2_{\text{subject}}$, representing between-subject variability, and $\epsilon_{ij} \sim \mathcal{N}(0, \sigma^2_{\text{residual}})$ is the residual error with variance $\sigma^2_{\text{residual}}$.

The proportion of total variance attributable to between-subject differences was quantified using the intraclass correlation coefficient (ICC):

$$\text{ICC} = \frac{\sigma^2_{\text{subject}}}{\sigma^2_{\text{subject}} + \sigma^2_{\text{residual}}}. \tag{23}$$

###### 2.13.7 Biological variability

To assess whether harmonisation preserved biological variability, we evaluate covariate effects before and after harmonisation. For each raw and harmonised feature, we fit a linear mixed-effects model with covariates as fixed effects and a subject-specific random intercept. For example, if age and timepoint were provided as biological covariates:

$$Y_{ij} = \beta_0 + \beta_{age} Age_{ij} + \beta_{sex} Sex_{ij} + u_i + \epsilon_{ij} \tag{24}$$

where $Y_{ij}$ is the feature for subject $i$ at observation $j$, $\beta_0$ is the fixed intercept, $\beta_{age}$ and $\beta_{time}$ are the fixed-effect coefficients for age and sex, respectively, $u_i$ is the subject-specific random intercept, and $\epsilon_{ij}$ is the residual error.

### 3 Harmonisation methods implemented in DHARM

DHARM implements several popular harmonisation methods into its framework. How to call each method and the required arguments are described in detail in the tool's online documentation.

#### 3.1 Linear Regression

The simplest harmonisation method implemented in DHARM is removal of batch effects through the linear modelling of fixed effects. In this regime, the batch effect is co-modelled alongside the covariates in the same format as described in equations 1b and 1d. We keep the same notation here, but instead of removing the covariate effects, batch effects are removed as shown in equation 25. Ordinary least squares (OLS) was fitted independently for each feature. Harmonisation was performed by subtracting the estimated batch contribution while preserving the fitted covariate effects.

$$y^{Harmonised}_{i,f} = Y_{i,f} - B_i \hat{\beta}_{f,b} \tag{25}$$

#### 3.2 Linear mixed modelling

Similarly to the simple linear regression case, we also allow harmonisation through the use of linear mixed modelling. Linear mixed-effects models were fitted independently to each feature. When the batch was specified as random,

$$y_{ij} = \beta_0 + \mathbf{X}_{cov} \boldsymbol{\beta_{cov}} + u_{b(i)} + \varepsilon_{ij}, with : u_b \sim \mathcal{N}(0, \sigma^2_b). \tag{26}$$

Alternatively, for repeated-measures data, the subject was included as the random intercept and the batch as a fixed effect. Harmonisation removed the estimated batch contribution while retaining other modelled effects.

#### 3.3 ComBat

ComBat models each standardised feature using batch-specific location and scale parameters. Full descriptions of ComBat are given in Fortin et al. (2017) and Johnson et al. (2007).

$$z_{ij} = \gamma_{b(i),j} + \delta_{b(i),j}\varepsilon_{ij}. \tag{27}$$

Raw batch parameters $\hat{\gamma}_{bj}$ and $\hat{\delta}_{bj}$ are estimated and shrunk towards batch-level priors using parametric empirical Bayes. Harmonised values are obtained as shown in Equation 28, followed by transformation back to the original feature scale, with covariate effects preserved by default. As described in Turnbull et al. (2026), covariate effects can be optionally removed as well.

$$z^*_{ij} = \frac{z_{ij} - \gamma^*_{b(i),j}}{\sqrt{\delta^*_{b(i),j}}} \tag{28}$$

#### 3.4 Reference-batch ComBat

Reference-batch ComBat uses the same empirical-Bayes framework but defines standardisation relative to a selected reference batch. The reference-batch mean and residual variance are used in place of the global mean and pooled variance, and its correction parameters are fixed, thereby leaving the reference batch unchanged while mapping other batches onto its scale (equation 29).

$$\gamma^*_{\text{ref},j} = 0, \delta^*_{\text{ref},j} = 1, \tag{29}$$

#### 3.5 CovBat

CovBat first applies ComBat location-scale harmonisation and then targets residual covariance differences. The ComBat-adjusted data are standardised and decomposed using principal component analysis (PCA). The smallest set of components explaining more than 95% of the cumulative variance was selected, their scores were harmonised across batches using ComBat, and the adjusted scores were projected back into the original feature space.

#### 3.6 ComBat-GAM/B-spline implementation

The nonlinear ComBat implementation extended the ComBat mean model by representing sufficiently continuous covariates using cubic B-spline basis functions. With $K = 6$ basis degrees of freedom and spline degree of 3 by default. The resulting spline-expanded design was fitted by OLS and subsequently passed through the standard ComBat empirical-Bayes location-scale correction. Thus, this implementation corresponds to spline-basis ComBat rather than a fully penalised GAM as done in Pomponio et al. (2020).

$$f(x) = \sum_{k=1}^{K} \beta_k B_k(x), \tag{30}$$

#### 3.7 Linear mixed modelling using IQMs

Image quality metrics (IQMs) can be incorporated as covariates within the linear mixed-effects harmonisation framework. For feature $j$, the data are modelled as shown in 31, where $\mathbf{q}_i$ denotes the IQM vector and $u_{b(i)}$ represents the batch-specific random effect. Harmonisation therefore removes the estimated batch component while conditioning on variation explained by the supplied IQMs. More information on the methodology is documented elsewhere (Bhalerao et al., 2026).

$$y_{ij} = \beta_0 + \mathbf{q}_i^\top \boldsymbol{\beta}_q + u_{b(i)} + \varepsilon_{ij}, \tag{31}$$

##### 3.8 Long-ComBat

Long-ComBat combines longitudinal mixed-effects modelling with ComBat empirical-Bayes correction. For each feature, a mixed model is fitted to the unstandardised measurements with subject-specific random intercepts, optional random time slopes, and fixed batch, timepoint, and covariate effects. Batch coefficients are converted to a weighted zero-sum parameterisation, and the resulting standardised batch location and scale estimates are shrunk using the ComBat empirical-Bayes procedure. The harmonised measurements are then reconstructed as shown in 32, thereby correcting longitudinal batch effects while retaining subject-specific and covariate-related variation.

$$y^*_{ij} = \frac{\sigma_j}{\sqrt{\delta^*_{b(i),j}}}\left(z_{ij} - \gamma^*_{b(i),j}\right) + \hat{y}_{ij} - \hat{\alpha}_{b(i),j}, \tag{32}$$

### 4 Supplementary information for the cross-sectional demonstration

#### 4.1 Dataset information

We demonstrate DHARM using structural measurements from the four UK Biobank sites (Stockport, Newcastle, Reading and Bristol) (Alfaro-Almagro et al., 2018; Miller et al., 2016), the Cambridge Centre for Ageing and Neuroscience neuroimaging dataset (CAMCAN) (Demetriou et al., 2025) and a subset of healthy, age-matched controls from the Oxford Brain Health Clinic study (OxBAC) (Griffanti et al., 2025). These datasets were all acquired at different sites with similar scanners and acquisition parameters (for the T1 and FLAIR data) but very different demographics. We apply the tool pre-harmonisation in order to calculate a baseline batch effect and again post-harmonisation, comparing metrics across different harmonisation methods in order to assess the performance of each method for different scenarios. We show here the data for 244 cortical thickness measures derived using the FreeSurfer implementation in the UKB pipeline from Alfaro-Almagro et al. (2018) (Fischl, 2012). The specific IDPs are provided in Table 3. A summary of the total sample size, age range, scanner type and acquisition parameters can be seen in Table 1.

| Dataset | Sample size used | Age range (years) | Percentage Female | Acquisition Information |
|---|---|---|---|---|
| UKB: Stockport | 500 | 47-83 | 54% | Siemens Skyra 3T (Alfaro-Almagro, 2021; Miller et al., 2016) |
| UKB: Newcastle | 500 | 50-84 | 56.6% | Siemens Skyra 3T(Alfaro-Almagro, 2021; Miller et al., 2016) |
| UKB: Reading | 500 | 50-82 | 57.6% | Siemens Skyra 3T(Alfaro-Almagro, 2021; Miller et al., 2016) |
| UKB: Bristol | 500 | 50 - 84 | 54.6% | Siemens Skyra 3T(Alfaro-Almagro, 2021; Miller et al., 2016) |
| CAMCAN | 474 | 18 - 89 | 50% | Siemens TIM Trio (Demetriou et al., 2025) |
| OxBAC | 95 | 60 - 92 | 54.1% | Siemens Prisma scanner (Griffanti et al., 2025) |

Table 1: Datasets used for the testing of different cross-sectional harmonisation methods

#### 4.2 List of Imaging-Derived Phenotypes - Cross-sectional demonstration

Table 2: Cortical thickness IDPs from the Destrieux atlas (aparc_a2009s) atlas. All listed IDPs were included for both left (LH) and right (RH) hemispheres. The common prefix `aparc_a2009s_lh_thickness` is omitted.

| **IDP list: aparc_a2009s_lh_thickness_** |
|---|
| g_cingul_post_dorsal |
| g_cingul_post_ventral |
| g_cuneus |
| g_front_inf_opercular |
| g_front_inf_orbital |
| g_front_inf_triangul |
| g_front_middle |
| g_front_sup |
| g_insular_short |
| g_oc_temp_lat_fusifor |
| g_oc_temp_med_lingual |
| g_oc_temp_med_parahip |
| g_occipital_middle |
| g_occipital_sup |
| g_orbital |
| g_pariet_inf_angular |
| g_pariet_inf_supramar |
| g_parietal_sup |
| g_postcentral |
| g_precentral |
| g_precuneus |
| g_rectus |
| g_subcallosal |
| g_temp_sup_g_t_transv |
| g_temp_sup_lateral |
| g_temp_sup_plan_polar |
| g_temp_sup_plan_tempo |
| g_temporal_inf |
| g_temporal_middle |
| lat_fis_ant_horizont |
| lat_fis_ant_vertical |
| lat_fis_post |
| pole_occipital |
| pole_temporal |
| s_calcarine |
| s_central |
| s_cingul_marginalis |
| s_circular_insula_ant |
| s_circular_insula_inf |
| s_circular_insula_sup |
| s_collat_transv_ant |
| s_collat_transv_post |
| s_front_inf |
| s_front_middle |
| s_front_sup |
| s_interm_prim_jensen |
| s_oc_temp_lat |
| s_occipital_ant |
| s_orbital_h_shaped |
| s_orbital_lateral |
| s_orbital_med_olfact |
| s_parieto_occipital |
| s_pericallosal |

| IDP: aparc_a2009s_lh_thickness_ |
|---|
| s_postcentral |
| s_precentral_inf_part |
| s_precentral_sup_part |
| s_suborbital |
| s_subparietal |
| s_temporal_inf |
| s_temporal_sup |
| s_temporal_transverse |

Table 3: Cortical thickness IDPs from the Desikan–Killiany atlas. All listed IDPs were included for both left (LH) and right (RH) hemispheres. The common prefix `aparc_desikan_[lh/rh]_thickness_` is omitted.

| IDP list: aparc_desikan_[lh/rh]_thickness_ |
|---|
| caudalanteriorcingulate |
| caudalmiddlefrontal |
| cuneus |
| entorhinal |
| frontalpole |
| fusiform |
| inferiorparietal |
| inferiortemporal |
| insula |
| isthmuscingulate |
| lateraloccipital |
| lateralorbitofrontal |
| lingual |
| medialorbitofrontal |
| middletemporal |
| paracentral |
| parsopercularis |
| parsorbitalis |
| parstriangularis |
| pericalcarine |
| postcentral |
| posteriorcingulate |
| precentral |
| precuneus |
| rostralanteriorcingulate |
| rostralmiddlefrontal |
| superiorfrontal |
| superiorparietal |
| superiortemporal |
| supramarginal |
| transversetemporal |

Table 4: Cortical thickness IDPs from the DKT atlas. All listed IDPs were included for both left (LH) and right (RH) hemispheres. The common prefix `aparc_dktatlas_lh_thickness_` is omitted.

| IDP list: aparc_dktatlas_lh_thickness_ |
|---|
| caudalanteriorcingulate |
| caudalmiddlefrontal |

| IDP list: aparc_dktatlas_lh_thickness_ |
|---|
| cuneus |
| entorhinal |
| fusiform |
| inferiorparietal |
| inferiortemporal |
| insula |
| isthmuscingulate |
| lateraloccipital |
| lateralorbitofrontal |
| lingual |
| medialorbitofrontal |
| middletemporal |
| paracentral |
| parsopercularis |
| parsorbitalis |
| parstriangularis |
| pericalcarine |
| postcentral |
| posteriorcingulate |
| precentral |
| precuneus |
| rostralanteriorcingulate |
| rostralmiddlefrontal |
| superiorfrontal |
| superiorparietal |
| superiortemporal |
| supramarginal |
| transversetemporal |

#### 4.3 List of Imaging-Derived Phenotypes - Repeated-measures demonstration

Table 5: T1-weighted grey matter volume imaging-derived phenotypes (IDPs) from the UK Biobank imaging-processing pipeline. Grey matter volumes were derived using atlas-based parcellation, including Harvard–Oxford cortical and subcortical regions and cerebellar regions. The common prefix `T1_GM_parcellation_` is omitted from the IDP names.

| IDP list: T1_GM_parcellation_ |
|---|
| L_Front_Pole_vol |
| R_Front_Pole_vol |
| L_Insula_vol |
| R_Insula_vol |
| L_Sup_Front_Gyr_vol |
| R_Sup_Front_Gyr_vol |
| L_Mid_Front_Gyr_vol |
| R_Mid_Front_Gyr_vol |
| L_Inf_Front_Gyr_pars_triangularis_vol |
| R_Inf_Front_Gyr_pars_triangularis_vol |
| L_Inf_Front_Gyr_pars_opercularis_vol |
| R_Inf_Front_Gyr_pars_opercularis_vol |
| L_Precentral_Gyr_vol |
| R_Precentral_Gyr_vol |
| L_Temp_Pole_vol |
| R_Temp_Pole_vol |

**IDP: T1_GM_parcellation_**

L_Sup_Temp_Gyr_ant_vol
R_Sup_Temp_Gyr_ant_vol
L_Sup_Temp_Gyr_post_vol
R_Sup_Temp_Gyr_post_vol
L_Mid_Temp_Gyr_ant_vol
R_Mid_Temp_Gyr_ant_vol
L_Mid_Temp_Gyr_post_vol
R_Mid_Temp_Gyr_post_vol
L_Mid_Temp_Gyr_temporoOcc_part_vol
R_Mid_Temp_Gyr_temporoOcc_part_vol
L_Inf_Temp_Gyr_ant_vol
R_Inf_Temp_Gyr_ant_vol
L_Inf_Temp_Gyr_post_vol
R_Inf_Temp_Gyr_post_vol
L_Inf_Temp_Gyr_temporoOcc_part_vol
R_Inf_Temp_Gyr_temporoOcc_part_vol
L_Postcentral_Gyr_vol
R_Postcentral_Gyr_vol
L_Sup_Parietal_Lobule_vol
R_Sup_Parietal_Lobule_vol
L_Supramarginal_Gyr_ant_vol
R_Supramarginal_Gyr_ant_vol
L_Supramarginal_Gyr_post_vol
R_Supramarginal_Gyr_post_vol
L_Angular_Gyr_vol
R_Angular_Gyr_vol
L_Lateral_Occ_Sup_vol
R_Lateral_Occ_Sup_vol
L_Lateral_Occ_Inf_vol
R_Lateral_Occ_Inf_vol
L_Intracalcarine_vol
R_Intracalcarine_vol
L_Front_Medial_vol
R_Front_Medial_vol
L_Juxtapositional_Lobule_vol
R_Juxtapositional_Lobule_vol
L_Subcallosal_vol
R_Subcallosal_vol
L_Paracingulate_Gyr_vol
R_Paracingulate_Gyr_vol
L_Cingulate_Gyr_ant_vol
R_Cingulate_Gyr_ant_vol
L_Cingulate_Gyr_post_vol
R_Cingulate_Gyr_post_vol
L_Precuneous_vol
R_Precuneous_vol
L_Cuneal_vol
R_Cuneal_vol
L_Front_Orbital_vol
R_Front_Orbital_vol
L_Parahippocampal_Gyr_ant_vol
R_Parahippocampal_Gyr_ant_vol
L_Parahippocampal_Gyr_post_vol
R_Parahippocampal_Gyr_post_vol
L_Lingual_Gyr_vol
R_Lingual_Gyr_vol
L_Temp_Fusiform_ant_vol

| IDP: T1_GM_parcellation_ |
|---|
| R_Temp_Fusiform_ant_vol |
| L_Temp_Fusiform_post_vol |
| R_Temp_Fusiform_post_vol |
| L_Temp_Occ_Fusiform_vol |
| R_Temp_Occ_Fusiform_vol |
| L_Occ_Fusiform_Gyr_vol |
| R_Occ_Fusiform_Gyr_vol |
| L_Front_Operculum_vol |
| R_Front_Operculum_vol |
| L_Central_Opercular_vol |
| R_Central_Opercular_vol |
| L_Parietal_Operculum_vol |
| R_Parietal_Operculum_vol |
| L_Planum_Polare_vol |
| R_Planum_Polare_vol |
| L_Heschl_Gyr_vol |
| R_Heschl_Gyr_vol |
| L_Planum_Tempe_vol |
| R_Planum_Tempe_vol |
| L_Supracalcarine_vol |
| R_Supracalcarine_vol |
| L_Occ_Pole_vol |
| R_Occ_Pole_vol |
| L_Thalamus_vol |
| R_Thalamus_vol |
| L_Caudate_vol |
| R_Caudate_vol |
| L_Putamen_vol |
| R_Putamen_vol |
| L_Pallidum_vol |
| R_Pallidum_vol |
| L_Hippocampus_vol |
| R_Hippocampus_vol |
| L_Amygdala_vol |
| R_Amygdala_vol |
| L_Vent_Striatum_vol |
| R_Vent_Striatum_vol |
| Brain_Stem_vol |
| L_I_IV_Cerebellum_vol |
| R_I_IV_Cerebellum_vol |
| L_V_Cerebellum_vol |
| R_V_Cerebellum_vol |
| L_VI_Cerebellum_vol |
| Vermis_VI_Cerebellum_vol |
| R_VI_Cerebellum_vol |
| L_Crus_I_Cerebellum_vol |
| R_Crus_I_Cerebellum_vol |
| L_Crus_II_Cerebellum_vol |
| Vermis_Crus_II_Cerebellum_vol |
| R_Crus_II_Cerebellum_vol |
| L_VIIb_Cerebellum_vol |
| Vermis_VIIb_Cerebellum_vol |
| R_VIIb_Cerebellum_vol |
| L_VIIIa_Cerebellum_vol |
| Vermis_VIIIa_Cerebellum_vol |
| R_VIIIa_Cerebellum_vol |
| L_VIIIb_Cerebellum_vol |

| IDP: T1_GM_parcellation_ |
|---|
| Vermis_VIIIb_Cerebellum_vol |
| R_VIIIb_Cerebellum_vol |
| L_IX_Cerebellum_vol |
| Vermis_IX_Cerebellum_vol |
| R_IX_Cerebellum_vol |
| L_X_Cerebellum_vol |
| Vermis_X_Cerebellum_vol |
| R_X_Cerebellum_vol |

##### 4.4 Harmonisation configurations

In this section, we present the full cross-sectional comparison report generated by the tool. All figures in the manuscript were generated and taken directly from the report. All commands used to generate the harmonised data are given below:

```python
def _extract_corrected_data(result_obj, method_name):
        if isinstance(result_obj, dict):
            if "bayesdata" not in result_obj:
                raise KeyError(
                    f"{method_name} returned a dict without "
                    f"'bayesdata'. Keys: {list(result_obj.keys())}"
                )
            return result_obj["bayesdata"]
        return result_obj
    results = {
        "Raw": raw_data.copy()
    }
    # OLS
    regression_results = HF.linearmodelling_harmonisation(
        data=raw_data,
        batch=batch_clean,
        covariates=mod,
        covariate_names=covariate_names,
        model_type="ols",
        batch_as_random=False,
    )
    results["Regression_OLS"] = regression_results["Residuals"]
    # ComBat
    combat_result = HF.combat_modular(
        raw_data,
        batch_clean.tolist(),
        mod=mod,
        covbat_mode=False,
    )
    results["ComBat"] = _extract_corrected_data(
        combat_result,
        "ComBat",
    )
    # CovBat
    covbat_result = HF.combat_modular(
        raw_data,
        batch_clean.tolist(),
        mod=mod,
        covbat_mode=True,
    )
    results["CovBat"] = _extract_corrected_data(
        covbat_result,
```

```
        "CovBat",
    )
    # ComBat GAM
    combat_gam_result = HF.combat_modular(
        raw_data,
        batch_clean.tolist(),
        mod=mod,
        covbat_mode=False,
        mean_model="gam",
    )
    results["Combat_GAM"] = _extract_corrected_data(
        combat_gam_result,
        "Combat_GAM",
    )
    # CovBat with UKB Stockport reference
    covbat_ref_result = HF.combat_modular(
        raw_data,
        batch_clean.tolist(),
        mod=mod,
        covbat_mode=True,
        mean_model="ols",
        ref_batch="UKB_Stockport",
    )
    results["CovBat_Ref_UKB_Stockport"] = _extract_corrected_data(
        covbat_ref_result,
        "CovBat_Ref_UKB_Stockport",
    )
```

All harmonised datasets were then combined into a single results dictionary, where each datasets key corresponds to the name used in the comparison report. The combined dictionary was then fed to the comparison reporting function, and the full report seen below was produced:

```
from DiagnoseHarmonisation import DiagnosticReport as DRep
DRep.CrossSectionalComparisonReport(
        results,
        batch=batch_list
        covariates=mod,
        report_name=report_name,
        covariate_names=covariate_names,
        save_data=True,
        timestamped_reports=False,
        save_dir=str(save_dir),
        shared_y=False,
    )
```

#### Cross-sectional Comparison Report including baseline

### Cross-sectional comparison of harmonisation methods

#### Contents


- Report preamble and overview
- Multi-method comparison overview
- Batch-covariate confounding diagnostics
- Diagnostics for Raw
- Diagnostics for Regression_OLS
- Diagnostics for ComBat
- Diagnostics for CovBat
- Diagnostics for Combat_GAM
- Diagnostics for CovBat_Ref_UKB_Stockport
- Method scorecard
- Best method summary


```
2026-09-16 12:08:37,307 — INFO — === Report session started ===
2026-09-16 12:08:37,307 — INFO — save_artifacts=False | save_dir=None
2026-09-16 12:08:37,311 — INFO — initialised HTML report at:
/vols/Scratch/hpn007/Project_2/BIRAX_UKB_CAMCAN/Thickness/Thickness_Baselines_subsampled_comparison_post_harmonisation.html
2026-09-16 12:08:37,311 — INFO — [TITLE] Cross-sectional comparison of harmonisation methods
```

#### Report preamble and overview

```
2026-09-16 12:08:37,311 — INFO — [SECTION] Report preamble and overview
This report compares multiple harmonisation methods using the same cross-sectional diagnostics and summarises best-performing methods by
The report is intended for side-by-side evaluation of raw and harmonised outputs but should be interpreted with caution.
As the best performing method by the metrics here may not be the best method for your downstream analysis.
For more information on usage, please see the documentation at https://jake-turnbull.github.io/HarmonisationDiagnostics/
```

#### Multi-method comparison overview

```
2026-09-16 12:08:37,311 — INFO — [SECTION] Multi-method comparison overview
This report compares multiple harmonisation methods using the same cross-sectional diagnostics and summarises best-performing methods by
Comparison dataset characteristics:
Validated methods: Raw, Regression_OLS, ComBat, CovBat, Combat_GAM, CovBat_Ref_UKB_Stockport
Total samples: 2569
Features: 244
Samples in each batch:
{np.str_('CAMCAN'): 474, np.str_('OxBAC'): 95, np.str_('UKB_Bristol'): 500, np.str_('UKB_Newcastle'): 500, np.str_('UKB_Reading'): 500, n
Missing data per method: Raw=0, Regression_OLS=0, ComBat=0, CovBat=0, Combat_GAM=0, CovBat_Ref_UKB_Stockport=0
Covariates: Age, Sex, Headsize_scaling
--------------------------------------------------------------------------------------------------------------------------------------------
```

#### Batch-covariate confounding diagnostics

```
2026-09-16 12:08:37,314 — INFO — [SECTION] Batch-covariate confounding diagnostics
Batch-covariate confounding quantifies potential imbalance of each covariate across batches,
Continuous covariates are summarised by the omnibus R2 from
covariate ~ batch; binary/categorical covariates are summarised by Cramer's V. Batch is always
treated as nominal categorical regardless of its input encoding. Larger values indicate greater
covariate imbalance across batches and therefore greater potential for confounding.

        Variable       Type Statistic  Value
             Age continuous        R2  0.370
             Sex     binary  CramersV  0.035
Headsize_scaling continuous        R2  0.003
--------------------------------------------------------------------------------------------------------------------------------------------
```

#### Diagnostics for Raw

```
2026-09-16 12:08:37,448 — INFO — [SECTION] Diagnostics for Raw
2026-09-16 12:08:37,448 — INFO — Running diagnostics for method: Raw
Omnibus R2 (variance associated with metadata) per PC for Raw:
                    PC1    PC2    PC3    PC4    PC5    PC6    PC7    PC8    PC9   PC10   PC11   PC12   PC13   PC14   PC15   PC16   PC17
batch             0.132  0.315  0.015  0.043  0.055  0.003  0.010  0.016  0.007  0.007  0.002  0.011  0.021  0.018  0.007  0.017  0.022
Age               0.025  0.124  0.005  0.049  0.003  0.006  0.002  0.040  0.000  0.001  0.004  0.012  0.026  0.001  0.006  0.001  0.021
Sex               0.002  0.000  0.001  0.006  0.004  0.025  0.053  0.005  0.037  0.003  0.008  0.004  0.000  0.004  0.003  0.001  0.026
```

```
Headsize_scaling  0.000  0.004  0.001  0.025  0.000  0.015  0.044  0.025  0.007  0.000  0.001  0.004  0.005  0.010  0.004  0.009  0.027
Summary metrics for Raw:
- median_abs_cohens_d: 0.2800524491552635
- prop_large_abs_cohens_d: 0.16120218579234974
- max_mahalanobis: 2.30901781104412
- median_r2_covariates: 0.01790266318079531
- median_r2_covariates_batch: 0.1551994115434031
- median_icc: 0.0
- prop_high_icc: 0.12295081967213115
- median_delta_r2_batch: 0.13002229912842833
- median_abs_log_variance_ratio: 0.21022686216238584
- prop_significant_levene: 0.10683060109289617
- prop_significant_ks: 0.7506830601092896
- max_frobenius_normalized: 0.44476285273668636
- max_abs_batch_pc_correlation: 0.397801280397195
- weighted_covariate_pc_correlation_top3: 0.07442622406236604
- prop_significant_icc: 0.13524590163934427
--------------------------------------------------------------------------------------------------------------------------------------------
```

#### Diagnostics for Regression_OLS

```
2026-09-16 12:08:58,664 — INFO — [SECTION] Diagnostics for Regression_OLS
2026-09-16 12:08:58,664 — INFO — Running diagnostics for method: Regression_OLS
Omnibus R2 (variance associated with metadata) per PC for Regression_OLS:
                    PC1    PC2    PC3    PC4    PC5    PC6    PC7    PC8    PC9   PC10   PC11   PC12   PC13   PC14   PC15   PC16   PC17
batch             0.112  0.000  0.006  0.000  0.007  0.008  0.007  0.005  0.000  0.000  0.003  0.001  0.008  0.004  0.002  0.004  0.000
Age               0.303  0.000  0.017  0.001  0.019  0.019  0.020  0.014  0.001  0.001  0.006  0.003  0.021  0.010  0.004  0.010  0.000
Sex               0.001  0.001  0.005  0.003  0.017  0.015  0.044  0.003  0.044  0.005  0.010  0.001  0.001  0.001  0.003  0.000  0.027
Headsize_scaling  0.001  0.000  0.023  0.004  0.015  0.001  0.038  0.035  0.022  0.002  0.001  0.000  0.000  0.002  0.005  0.001  0.030
Summary metrics for Regression_OLS:
- median_abs_cohens_d: 2.4252879584671243e-14
- prop_large_abs_cohens_d: 0.0
- max_mahalanobis: 1.8648220303774088e-12
- median_r2_covariates: 0.12491618516188863
- median_r2_covariates_batch: 0.12491618516188863
- median_icc: 0.0
- prop_high_icc: 0.0
- median_delta_r2_batch: 0.0
- median_abs_log_variance_ratio: 0.08355106546578853
- prop_significant_levene: 0.10683060109289617
- prop_significant_ks: 0.01092896174863388
- max_frobenius_normalized: 0.44820258965256377
- max_abs_batch_pc_correlation: 0.23571292042162037
- weighted_covariate_pc_correlation_top3: 0.17124498110578323
- prop_significant_icc: 0.0
--------------------------------------------------------------------------------------------------------------------------------------------
```

#### Diagnostics for ComBat

```
2026-09-16 12:09:17,596 — INFO — [SECTION] Diagnostics for ComBat
2026-09-16 12:09:17,597 — INFO — Running diagnostics for method: ComBat
Omnibus R2 (variance associated with metadata) per PC for ComBat:
                    PC1    PC2    PC3    PC4    PC5    PC6    PC7    PC8    PC9   PC10   PC11   PC12   PC13   PC14   PC15   PC16   PC17
batch             0.113  0.000  0.003  0.001  0.006  0.006  0.006  0.004  0.000  0.000  0.003  0.001  0.007  0.003  0.002  0.004  0.000
Age               0.302  0.000  0.014  0.001  0.018  0.017  0.021  0.013  0.000  0.001  0.008  0.002  0.017  0.009  0.005  0.012  0.000
Sex               0.001  0.001  0.006  0.003  0.019  0.016  0.041  0.002  0.044  0.006  0.011  0.001  0.000  0.001  0.002  0.000  0.027
Headsize_scaling  0.001  0.000  0.024  0.004  0.015  0.001  0.034  0.030  0.024  0.003  0.001  0.000  0.000  0.001  0.005  0.001  0.029
Summary metrics for ComBat:
- median_abs_cohens_d: 0.008788991113098398
- prop_large_abs_cohens_d: 0.0
- max_mahalanobis: 0.772764513136949
- median_r2_covariates: 0.1243646999519748
- median_r2_covariates_batch: 0.12443238951820068
- median_icc: 0.0
- prop_high_icc: 0.0
- median_delta_r2_batch: 6.164203340724983e-05
- median_abs_log_variance_ratio: 0.0264210411423831
- prop_significant_levene: 0.01912568306010929
- prop_significant_ks: 0.0
- max_frobenius_normalized: 0.3852060575414918
- max_abs_batch_pc_correlation: 0.23664419344560972
- weighted_covariate_pc_correlation_top3: 0.17201683590032243
- prop_significant_icc: 0.0
--------------------------------------------------------------------------------------------------------------------------------------------
```

#### Diagnostics for CovBat

```
2026-09-16 12:09:37,665 — INFO — [SECTION] Diagnostics for CovBat
2026-09-16 12:09:37,665 — INFO — Running diagnostics for method: CovBat
Omnibus R2 (variance associated with metadata) per PC for CovBat:
                    PC1    PC2    PC3    PC4    PC5    PC6    PC7    PC8    PC9   PC10   PC11   PC12   PC13   PC14   PC15   PC16   PC17
batch             0.112  0.000  0.003  0.001  0.006  0.006  0.006  0.004  0.000  0.000  0.003  0.001  0.007  0.004  0.002  0.003  0.000
Age               0.305  0.002  0.013  0.001  0.019  0.017  0.020  0.013  0.000  0.001  0.008  0.002  0.016  0.010  0.005  0.012  0.000
Sex               0.001  0.000  0.006  0.003  0.018  0.017  0.042  0.001  0.044  0.005  0.011  0.001  0.000  0.001  0.002  0.000  0.027
```

```
Headsize_scaling  0.001  0.002  0.022  0.004  0.015  0.001  0.035  0.029  0.025  0.003  0.001  0.000  0.000  0.001  0.005  0.001  0.028
Summary metrics for CovBat:
- median_abs_cohens_d: 0.007731128336254581
- prop_large_abs_cohens_d: 0.0
- max_mahalanobis: 0.7270184545627915
- median_r2_covariates: 0.1263927093229973
- median_r2_covariates_batch: 0.12648341646556782
- median_icc: 0.0
- prop_high_icc: 0.0
- median_delta_r2_batch: 5.381241299878825e-05
- median_abs_log_variance_ratio: 0.028457024724630527
- prop_significant_levene: 0.029508196721311476
- prop_significant_ks: 0.0
- max_frobenius_normalized: 0.36197337358050335
- max_abs_batch_pc_correlation: 0.23662414605602292
- weighted_covariate_pc_correlation_top3: 0.17431883803890427
- prop_significant_icc: 0.0
--------------------------------------------------------------------------------------------------------------------------------------------
```

#### Diagnostics for Combat_GAM

```
2026-09-16 12:09:57,831 — INFO — [SECTION] Diagnostics for Combat_GAM
2026-09-16 12:09:57,831 — INFO — Running diagnostics for method: Combat_GAM
Omnibus R2 (variance associated with metadata) per PC for Combat_GAM:
                    PC1    PC2    PC3    PC4    PC5    PC6    PC7    PC8    PC9   PC10   PC11   PC12   PC13   PC14   PC15   PC16   PC17
batch             0.101  0.000  0.002  0.006  0.002  0.005  0.006  0.002  0.001  0.001  0.003  0.000  0.004  0.003  0.002  0.005  0.000
Age               0.293  0.000  0.013  0.000  0.011  0.019  0.020  0.012  0.000  0.001  0.007  0.002  0.015  0.007  0.006  0.013  0.000
Sex               0.001  0.001  0.006  0.003  0.022  0.013  0.041  0.002  0.043  0.006  0.011  0.001  0.000  0.001  0.003  0.000  0.027
Headsize_scaling  0.001  0.000  0.024  0.004  0.015  0.000  0.035  0.030  0.022  0.003  0.001  0.000  0.000  0.001  0.005  0.001  0.029
Summary metrics for Combat_GAM:
- median_abs_cohens_d: 0.02472825279252342
- prop_large_abs_cohens_d: 0.0
- max_mahalanobis: 0.9916183697965305
- median_r2_covariates: 0.117604616054583
- median_r2_covariates_batch: 0.1179937692419149
- median_icc: 0.0
- prop_high_icc: 0.0
- median_delta_r2_batch: 0.0005425693455869962
- median_abs_log_variance_ratio: 0.029088539233093753
- prop_significant_levene: 0.016939890710382512
- prop_significant_ks: 0.01980874316939891
- max_frobenius_normalized: 0.3949308432730692
- max_abs_batch_pc_correlation: 0.22253287813889874
- weighted_covariate_pc_correlation_top3: 0.16915392152855233
- prop_significant_icc: 0.0
--------------------------------------------------------------------------------------------------------------------------------------------
```

#### Diagnostics for CovBat_Ref_UKB_Stockport

```
2026-09-16 12:10:17,982 — INFO — [SECTION] Diagnostics for CovBat_Ref_UKB_Stockport
2026-09-16 12:10:17,982 — INFO — Running diagnostics for method: CovBat_Ref_UKB_Stockport
Omnibus R2 (variance associated with metadata) per PC for CovBat_Ref_UKB_Stockport:
                    PC1    PC2    PC3    PC4    PC5    PC6    PC7    PC8    PC9   PC10   PC11   PC12   PC13   PC14   PC15   PC16   PC17
batch             0.112  0.000  0.003  0.001  0.006  0.006  0.006  0.004  0.000  0.000  0.003  0.001  0.007  0.004  0.002  0.003  0.000
Age               0.305  0.002  0.013  0.001  0.019  0.017  0.020  0.013  0.000  0.001  0.008  0.002  0.016  0.010  0.005  0.012  0.000
Sex               0.001  0.000  0.006  0.003  0.018  0.017  0.042  0.001  0.044  0.005  0.011  0.001  0.000  0.001  0.002  0.000  0.027
Headsize_scaling  0.001  0.002  0.022  0.004  0.015  0.001  0.035  0.029  0.025  0.003  0.001  0.000  0.000  0.001  0.005  0.001  0.028
Summary metrics for CovBat_Ref_UKB_Stockport:
- median_abs_cohens_d: 0.007731128336254581
- prop_large_abs_cohens_d: 0.0
- max_mahalanobis: 0.7270184545627915
- median_r2_covariates: 0.1263927093229973
- median_r2_covariates_batch: 0.12648341646556782
- median_icc: 0.0
- prop_high_icc: 0.0
- median_delta_r2_batch: 5.381241299878825e-05
- median_abs_log_variance_ratio: 0.028457024724630527
- prop_significant_levene: 0.029508196721311476
- prop_significant_ks: 0.0
- max_frobenius_normalized: 0.36197337358050335
- max_abs_batch_pc_correlation: 0.23662414605602292
- weighted_covariate_pc_correlation_top3: 0.17431883803890427
- prop_significant_icc: 0.0
--------------------------------------------------------------------------------------------------------------------------------------------
```

#### Method scorecard

```
2026-09-16 12:10:38,293 — INFO — [SECTION] Method scorecard
Method scorecard by diagnostic category (higher is better within each block):
Additive
                  method  median_abs_cohens_d  prop_large_abs_cohens_d  max_mahalanobis  additive_score
                  CovBat         7.731128e-03                 0.000000     7.270185e-01        0.885845
CovBat_Ref_UKB_Stockport         7.731128e-03                 0.000000     7.270185e-01        0.885845
                  ComBat         8.788991e-03                 0.000000     7.727645e-01        0.877981
```

```
           Combat_GAM        2.472825e-02              0.000000     9.916184e-01        0.827416
        Regression_OLS        2.425288e-14              0.000000     1.864822e-12        1.000000
                   Raw        2.800524e-01              0.161202     2.309018e+00        0.000000
--------------------------------------------------------------------------------------------------------------------------------
Multiplicative
                method  median_abs_log_variance_ratio  prop_significant_levene  multiplicative_score
                CovBat                       0.028457                 0.029508              0.988923
CovBat_Ref_UKB_Stockport                     0.028457                 0.029508              0.988923
                ComBat                       0.026421                 0.019126              1.000000
            Combat_GAM                       0.029089                 0.016940              0.985487
        Regression_OLS                       0.083551                 0.106831              0.689183
                   Raw                       0.210227                 0.106831              0.000000
--------------------------------------------------------------------------------------------------------------------------------
Linear Modelling
                method  median_icc  prop_high_icc  median_delta_r2_batch  linear_modeling_score
                CovBat         0.0       0.000000               0.000054                    1.0
CovBat_Ref_UKB_Stockport       0.0       0.000000               0.000054                    1.0
                ComBat         0.0       0.000000               0.000062                    1.0
            Combat_GAM         0.0       0.000000               0.000543                    1.0
        Regression_OLS         0.0       0.000000               0.000000                    1.0
                   Raw         0.0       0.122951               0.130022                    1.0
--------------------------------------------------------------------------------------------------------------------------------
Distributional
                method  prop_significant_ks  max_frobenius_normalized  distributional_score
                CovBat             0.000000                  0.361973              1.000000
CovBat_Ref_UKB_Stockport           0.000000                  0.361973              1.000000
                ComBat             0.000000                  0.385206              0.865285
            Combat_GAM             0.019809                  0.394931              0.795702
        Regression_OLS             0.010929                  0.448203              0.492721
                   Raw             0.750683                  0.444763              0.019945
--------------------------------------------------------------------------------------------------------------------------------
Principal Component Analysis
                method  max_abs_batch_pc_correlation  principal_component_analysis_score
                CovBat                      0.236624                            0.919602
CovBat_Ref_UKB_Stockport                    0.236624                            0.919602
                ComBat                      0.236644                            0.919487
            Combat_GAM                      0.222533                            1.000000
        Regression_OLS                      0.235713                            0.924801
                   Raw                      0.397801                            0.000000
--------------------------------------------------------------------------------------------------------------------------------
Biological Effects
                method  median_r2_covariates  median_r2_covariates_batch  weighted_covariate_pc_correlation_top3  biological_effects_sc
                CovBat              0.126393                    0.126483                                0.174319               1.000
CovBat_Ref_UKB_Stockport            0.126393                    0.126483                                0.174319               1.000
                ComBat              0.124365                    0.124432                                0.172017               0.979
            Combat_GAM              0.117605                    0.117994                                0.169154               0.933
        Regression_OLS              0.124916                    0.124916                                0.171245               0.977
                   Raw              0.017903                    0.155199                                0.074426               0.000
--------------------------------------------------------------------------------------------------------------------------------
Interpretation guide:
Additive metrics reward smaller absolute mean-shift and distance effects; multiplicative metrics reward stable variance structure;
linear-modelling metrics reward weaker residual batch random effects; distributional metrics reward closer overall distributions;
PCA metrics reward weaker batch-aligned structure in the principal components; and biological metrics reward stronger preserved covariate
```

#### Best method summary

```
2026-09-16 12:10:38,298 — INFO — [SECTION] Best method summary
Best overall method: CovBat
Strongest preserved biological effects: CovBat
Best mean_shift: Regression_OLS
Best variance_alignment: ComBat
Best batch_random_effect: CovBat
Best distribution_similarity: CovBat
Best covariance_alignment: CovBat
Best pc_batch_association: Combat_GAM
Best biological_effects: CovBat
Best overall method is CovBat, but diagnostics disagree across domains. This indicates trade-offs between harmonisation targets rather th
Saved scorecard CSV: /vols/Scratch/hpn007/Project_2/BIRAX_UKB_CAMCAN/Thickness/Thickness_Baselines_subsampled_comparison_post_harmonisati
2026-09-16 12:11:42,270 — INFO — === Report session finished ===
```

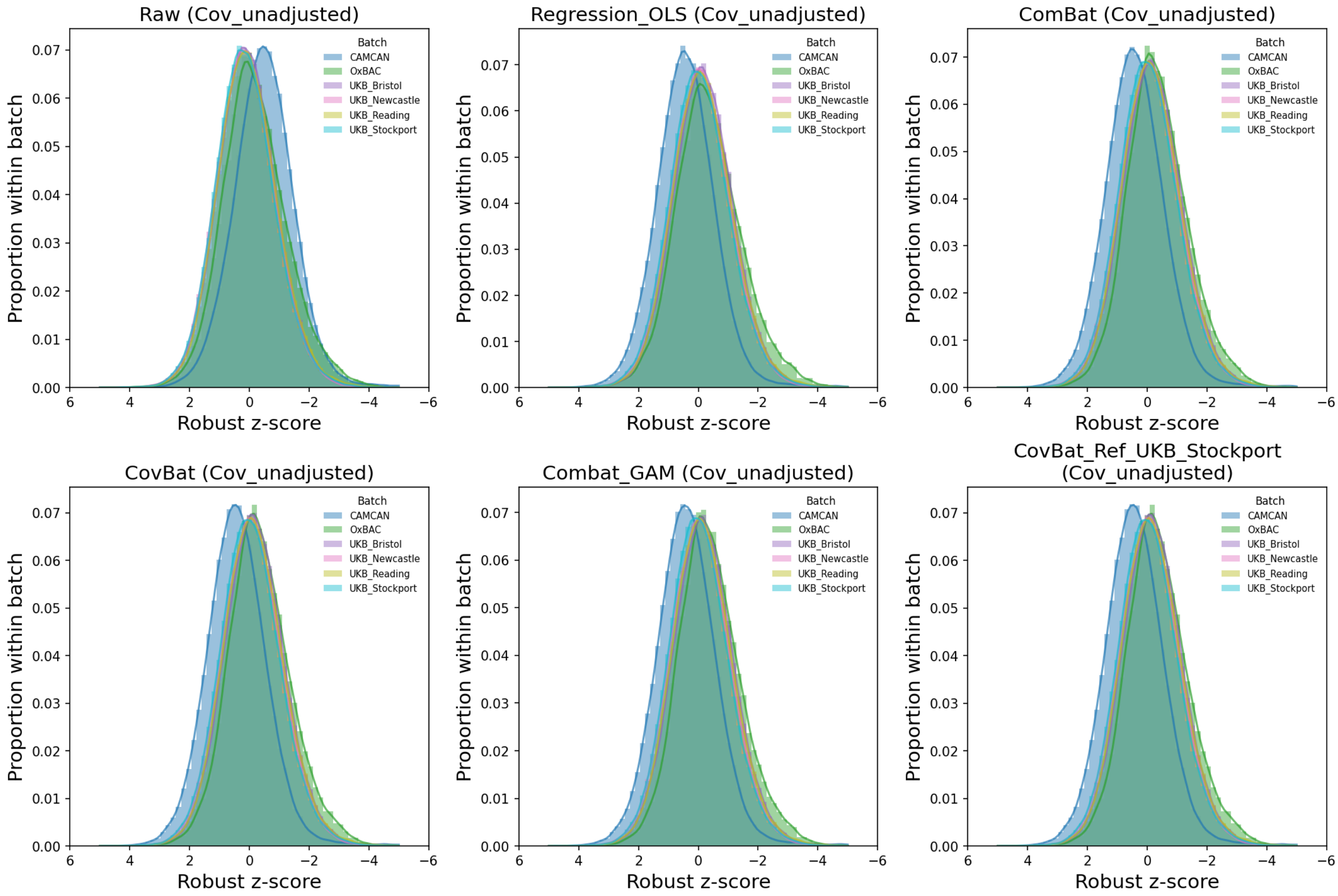


Comparison: raw z-score distributions

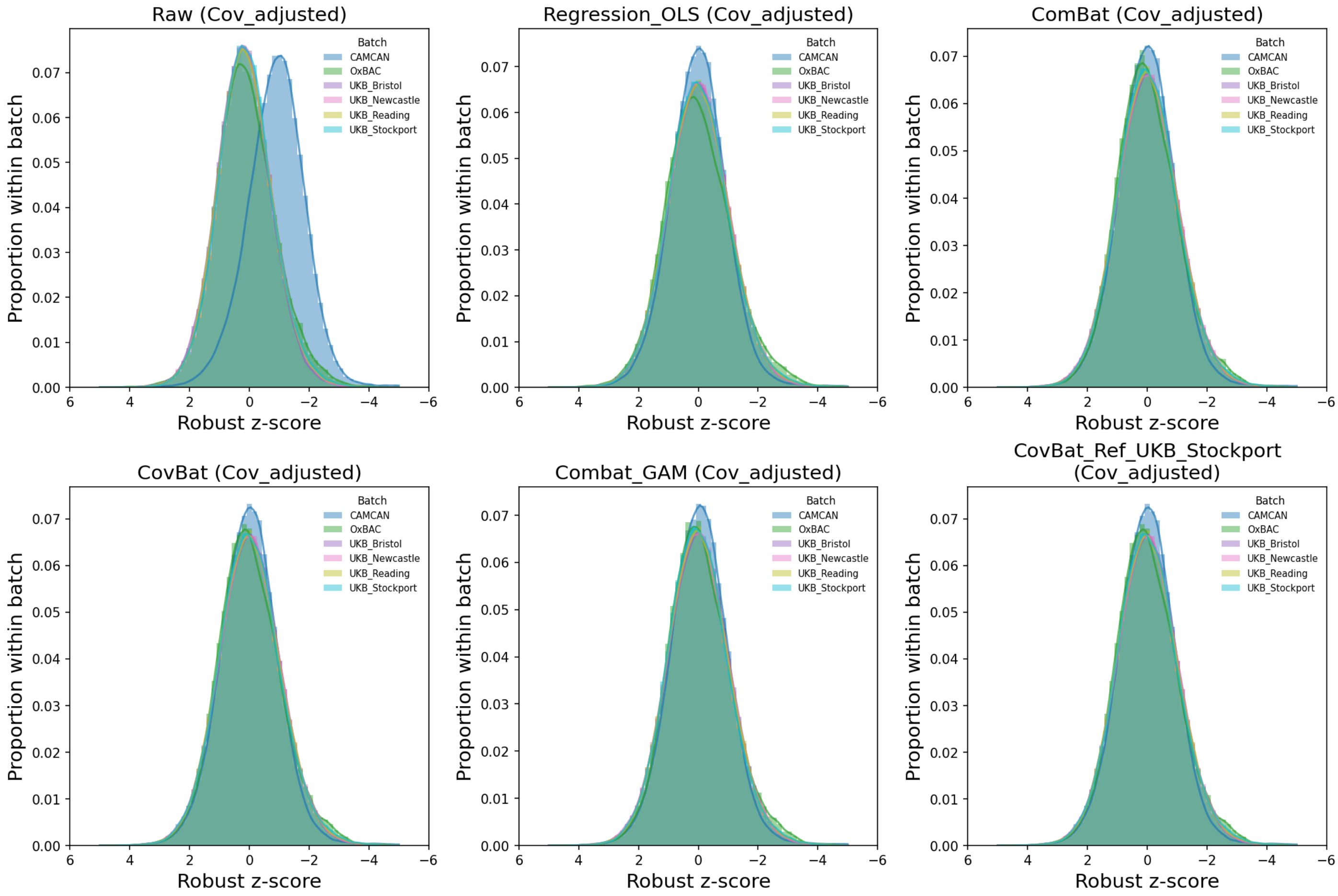

Comparison: residual z-score distributions

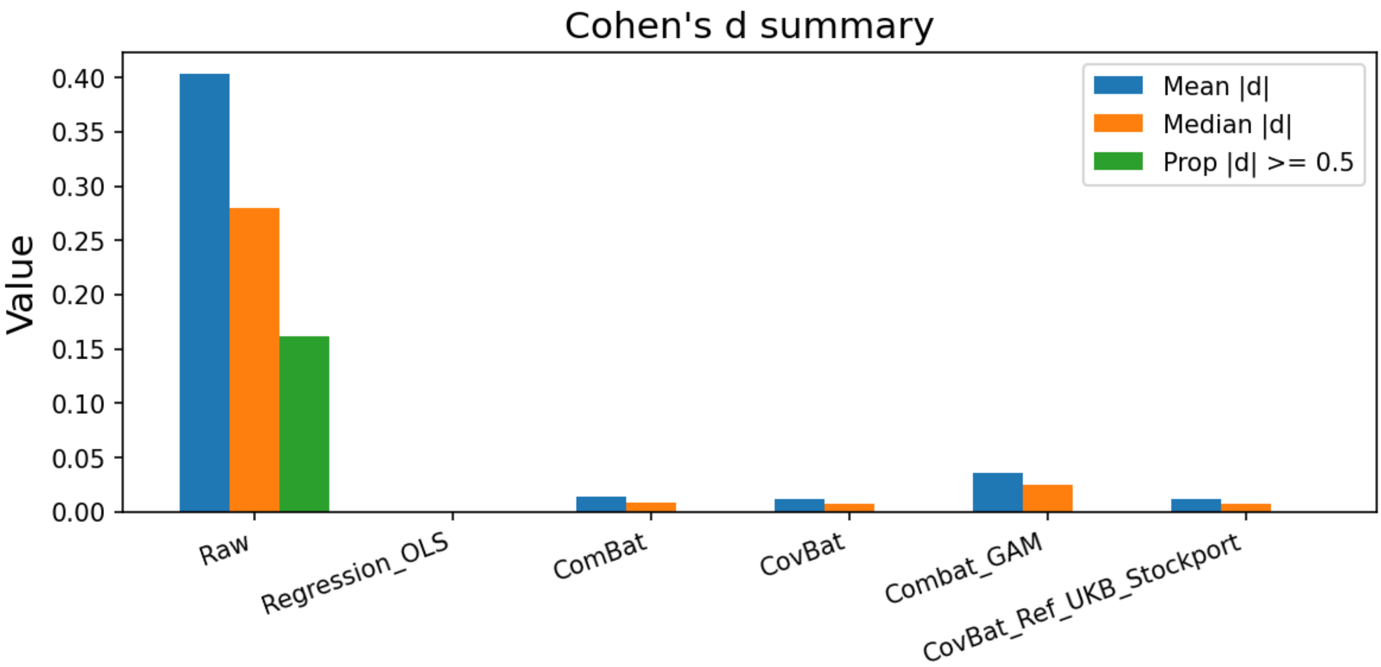


Comparison: Cohen's d summary

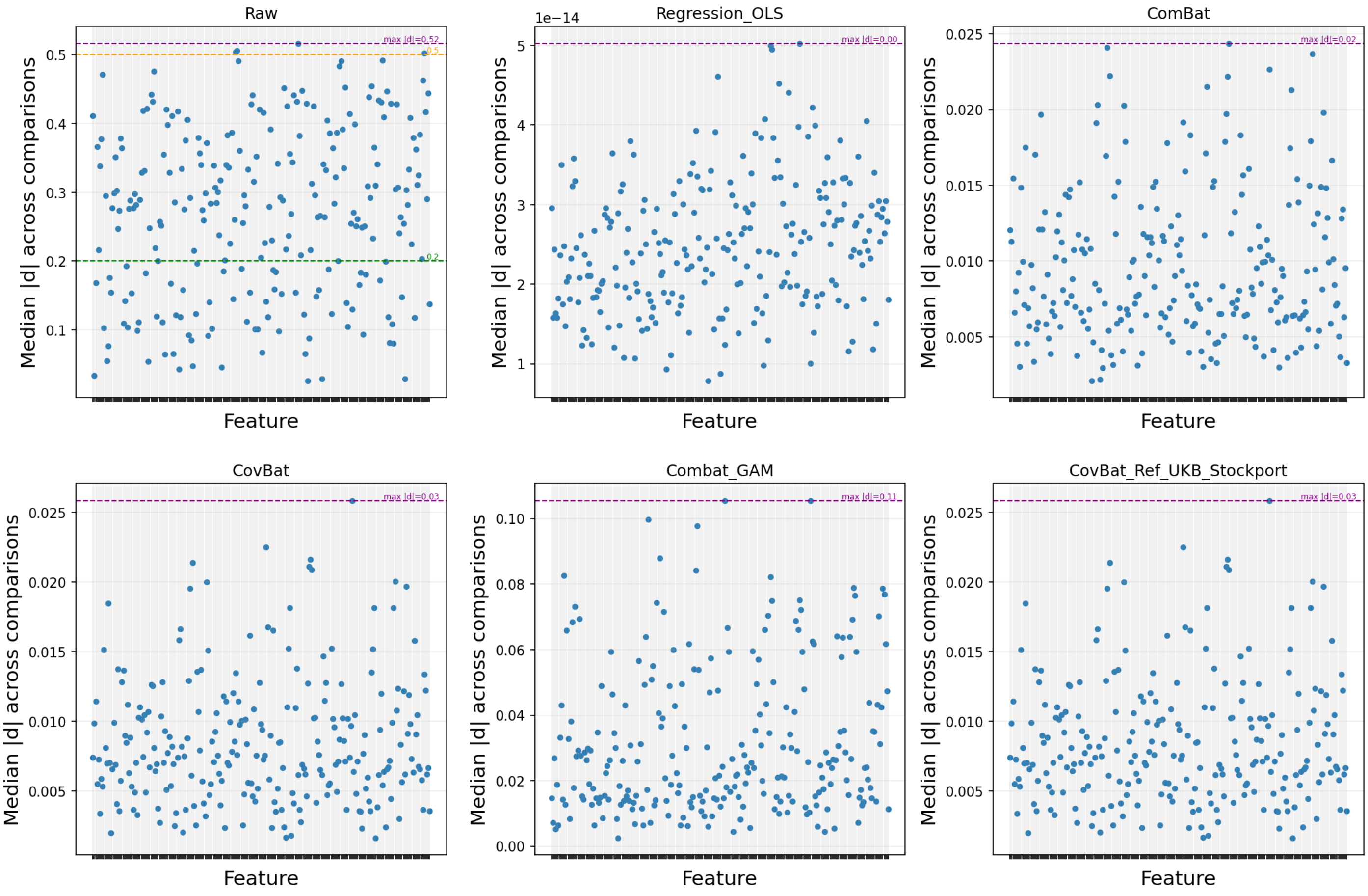


Comparison: Cohen's d feature-wise

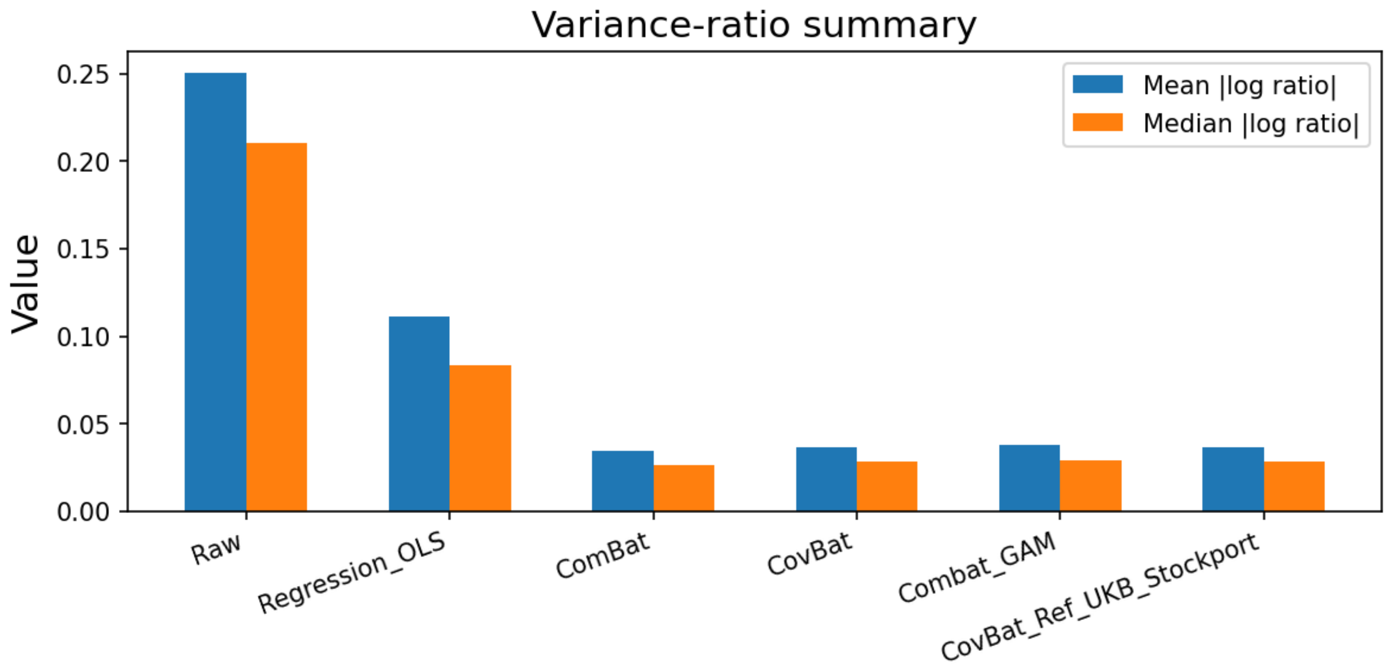


Comparison: variance ratios summary

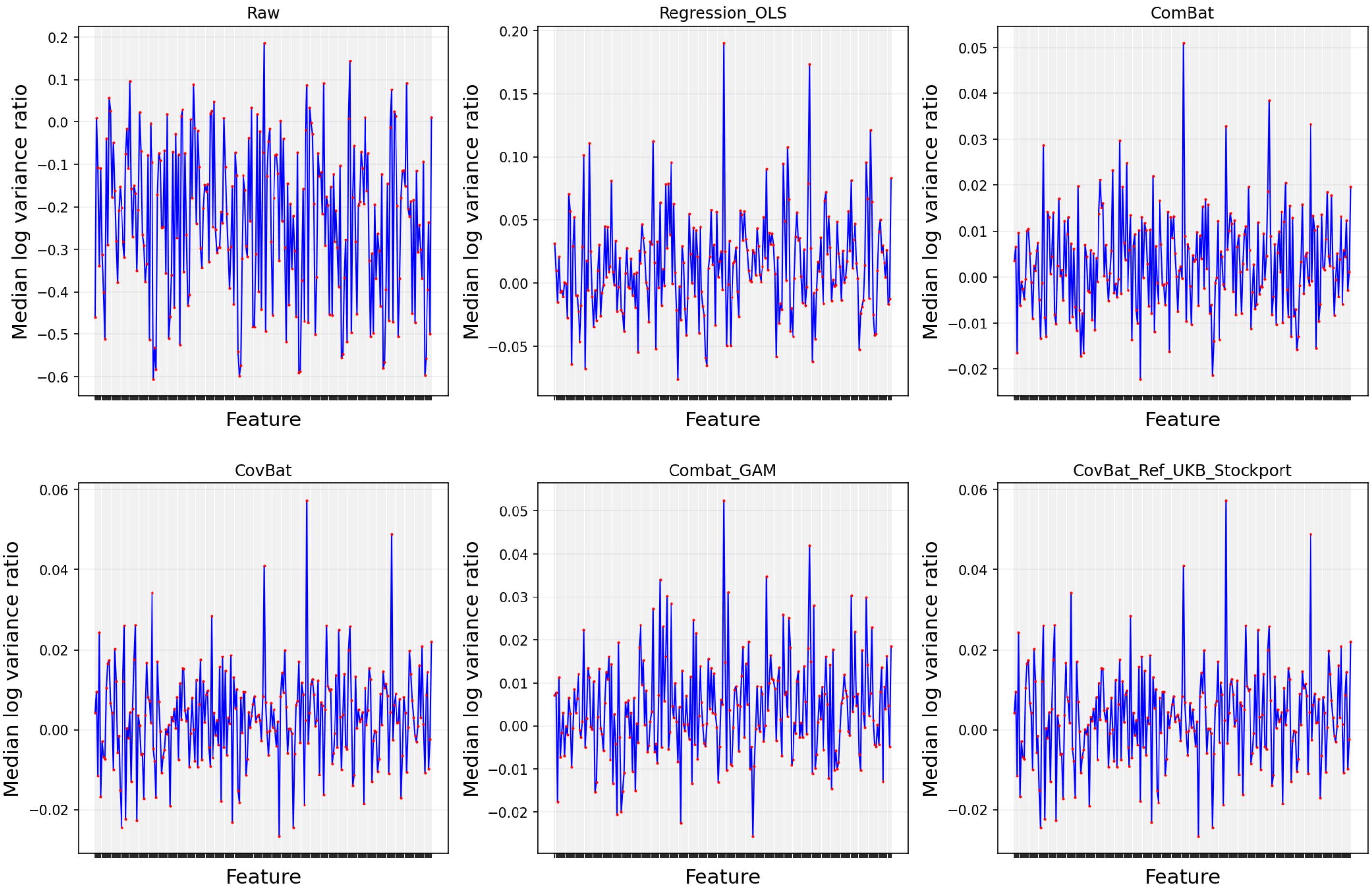


Comparison: variance ratios feature-wise

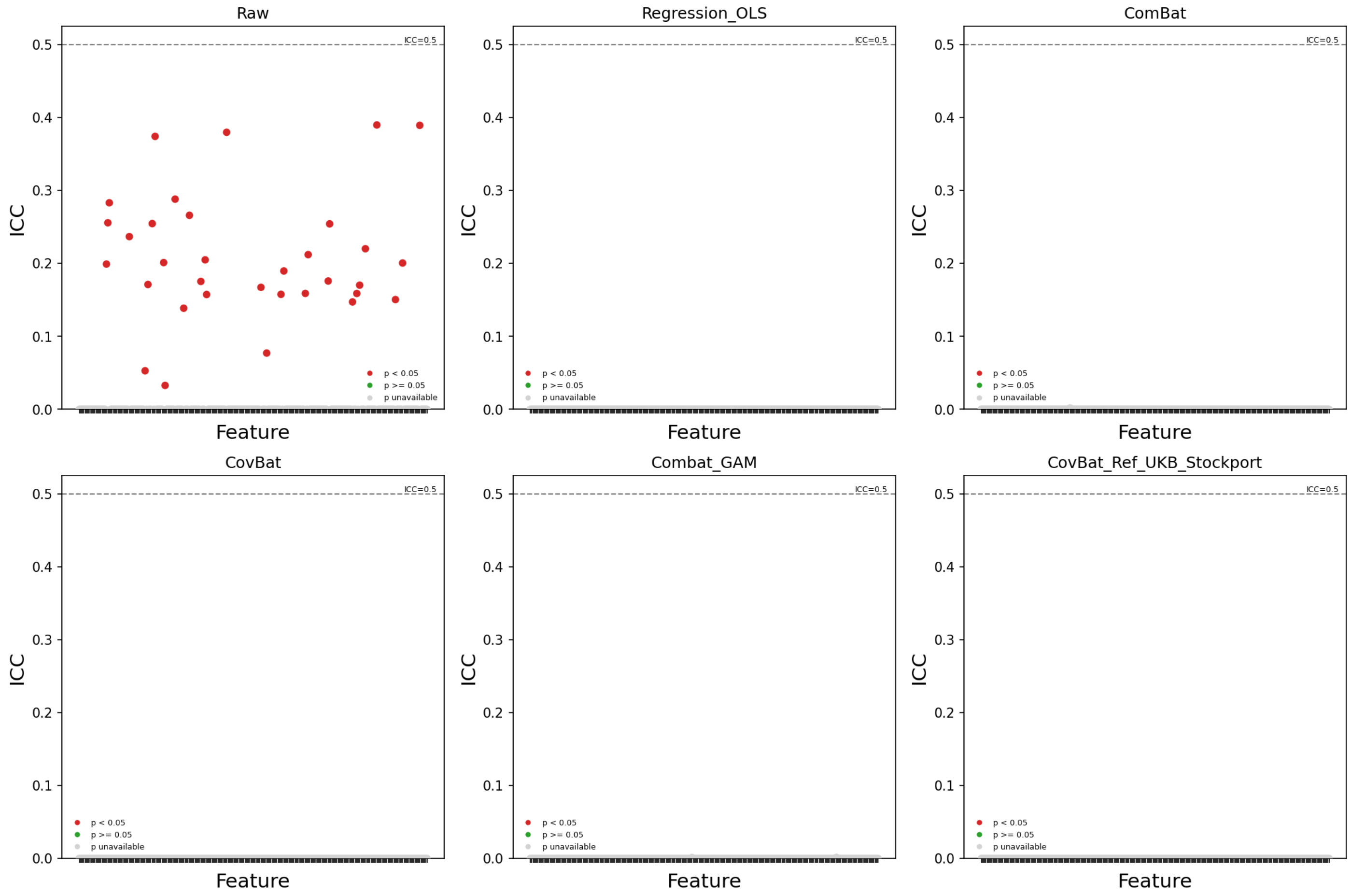


Comparison: LMM ICC per feature

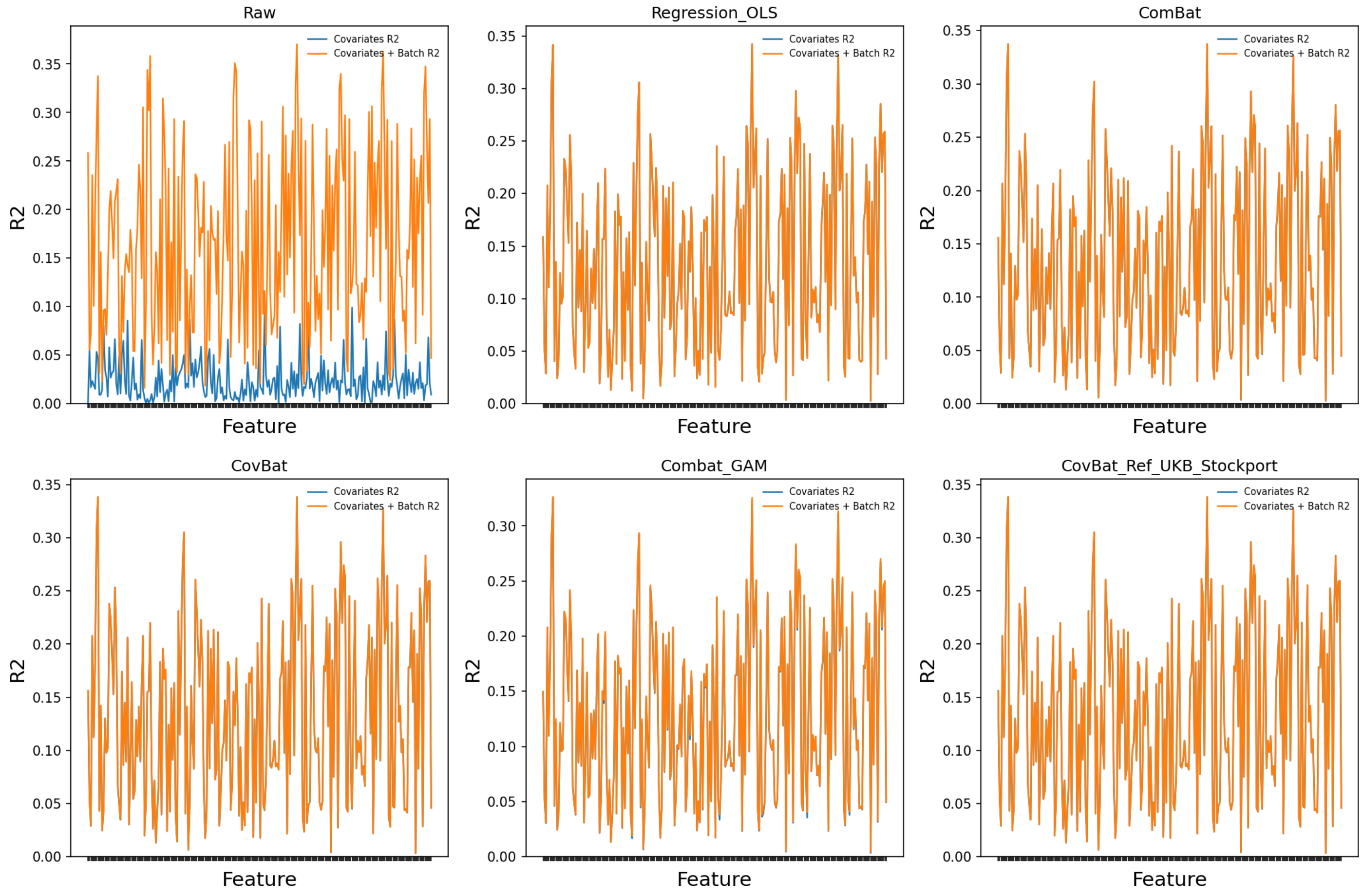


Comparison: covariates and covariates+batch R2

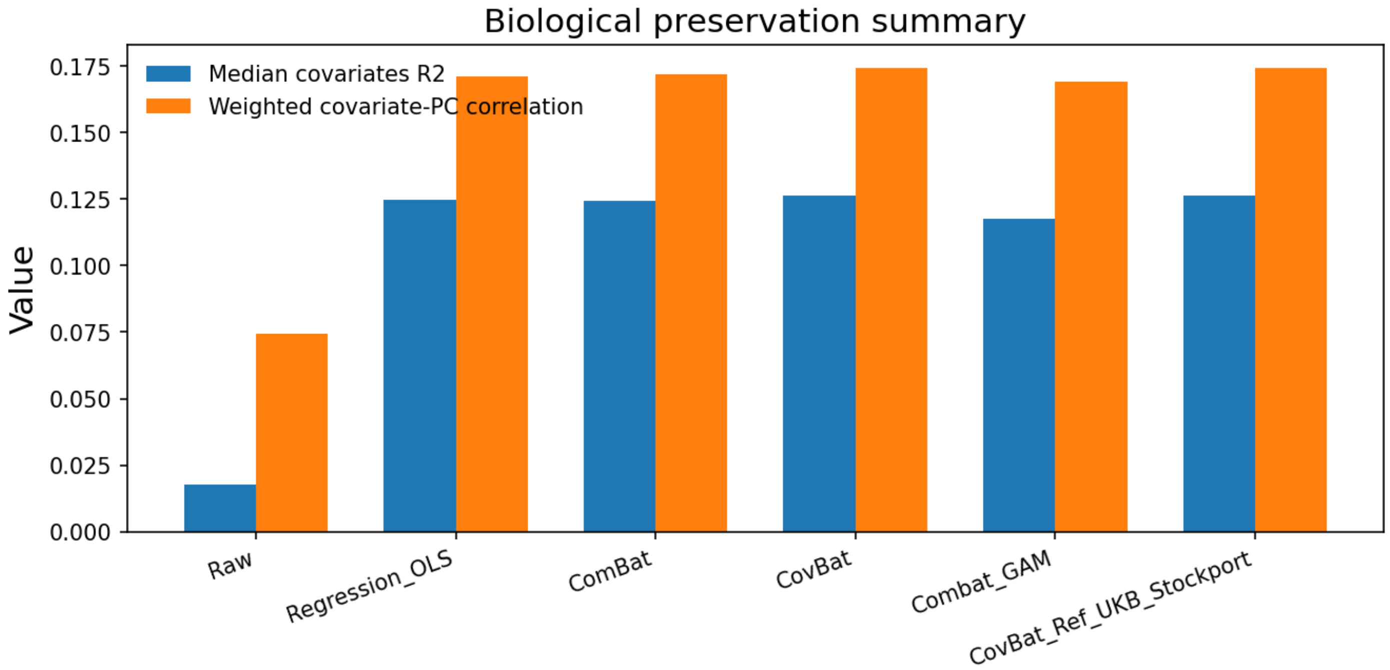


Comparison: biological preservation summary

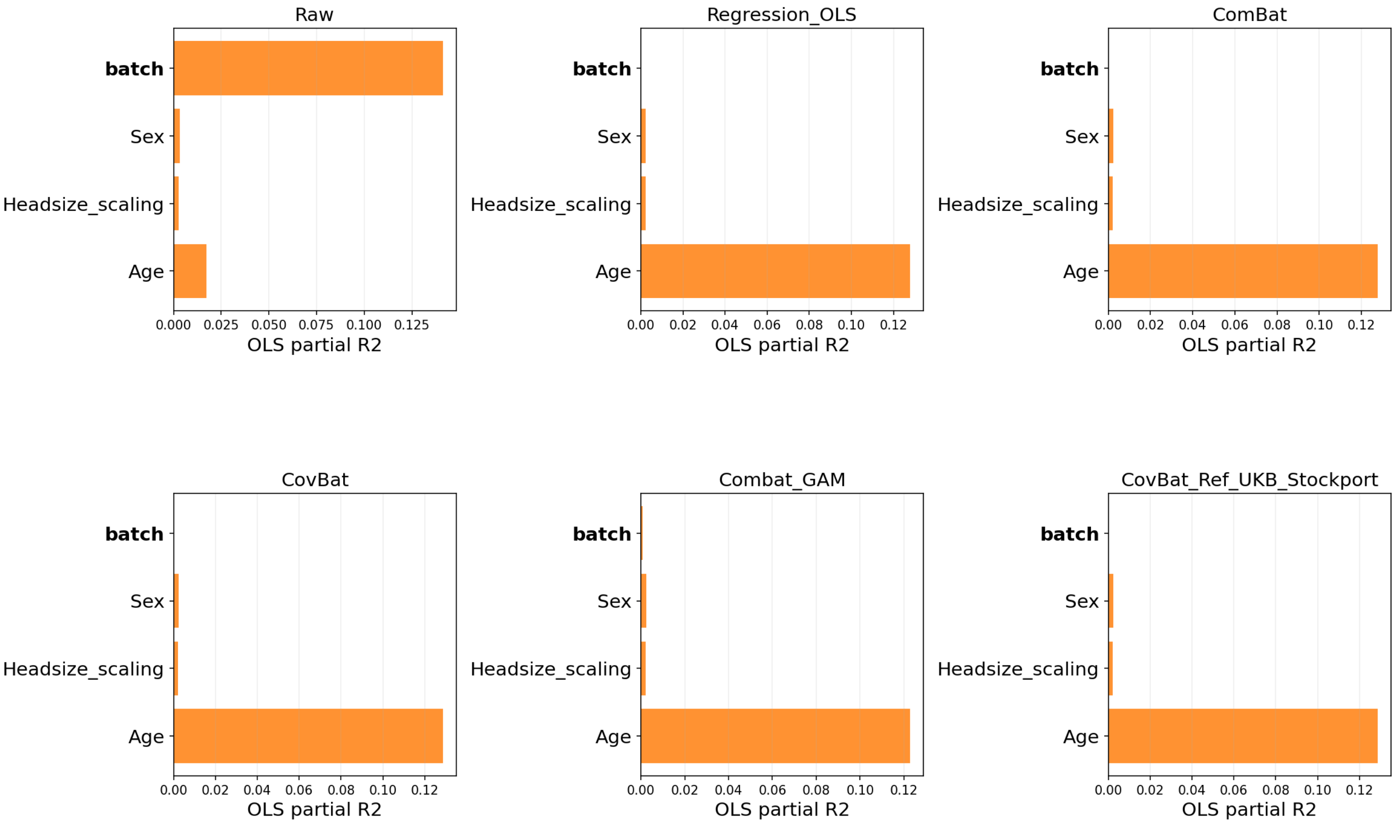


Comparison: covariate variance from OLS

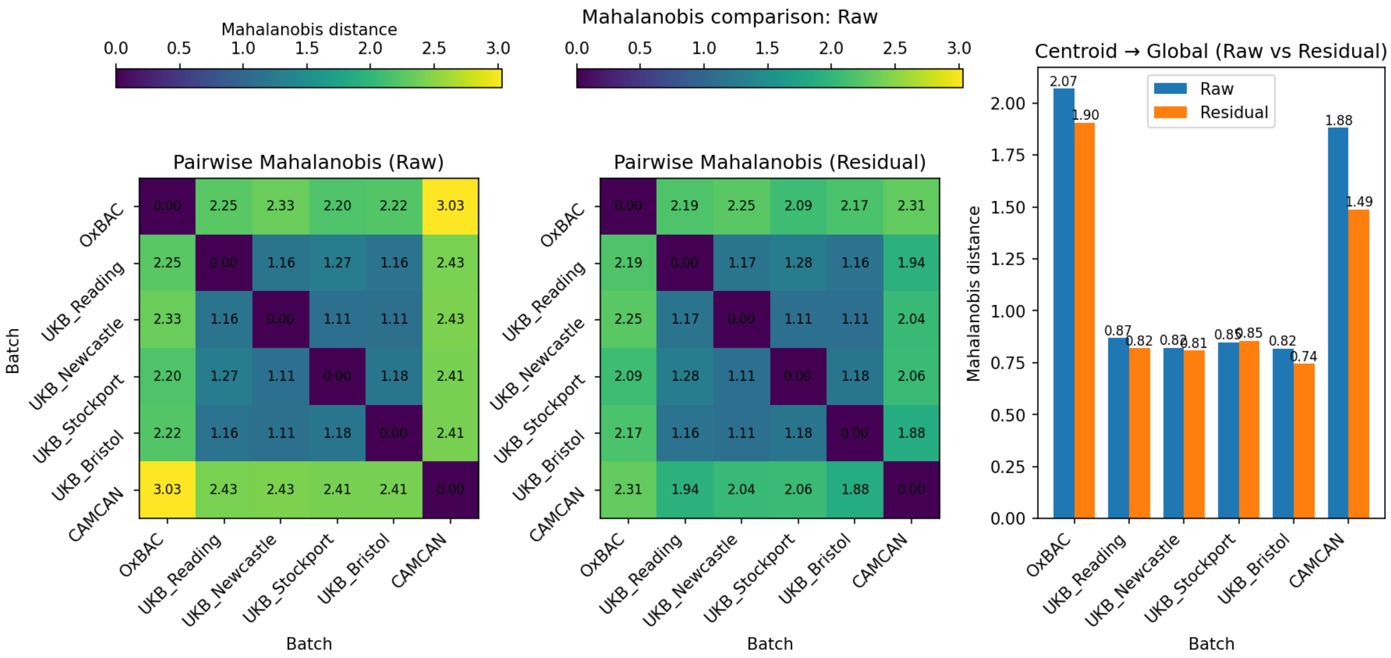


Comparison: Mahalanobis (Raw)

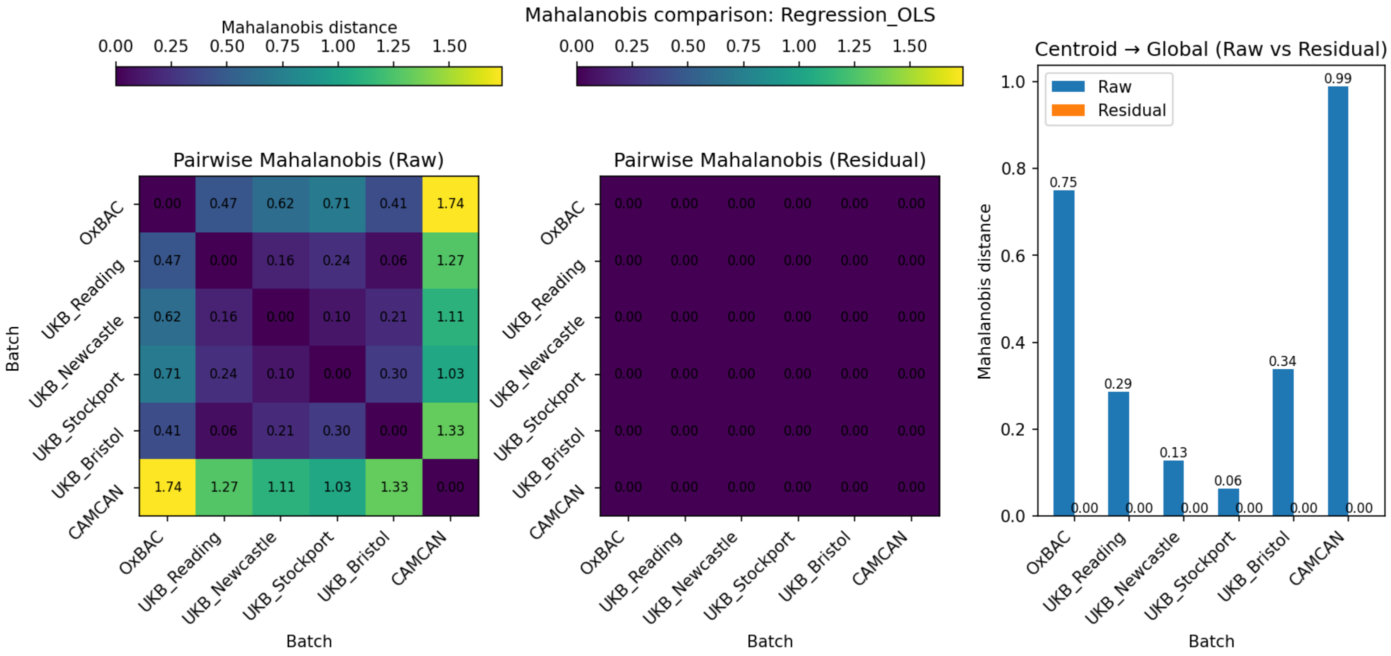


Comparison: Mahalanobis (Regression_OLS)

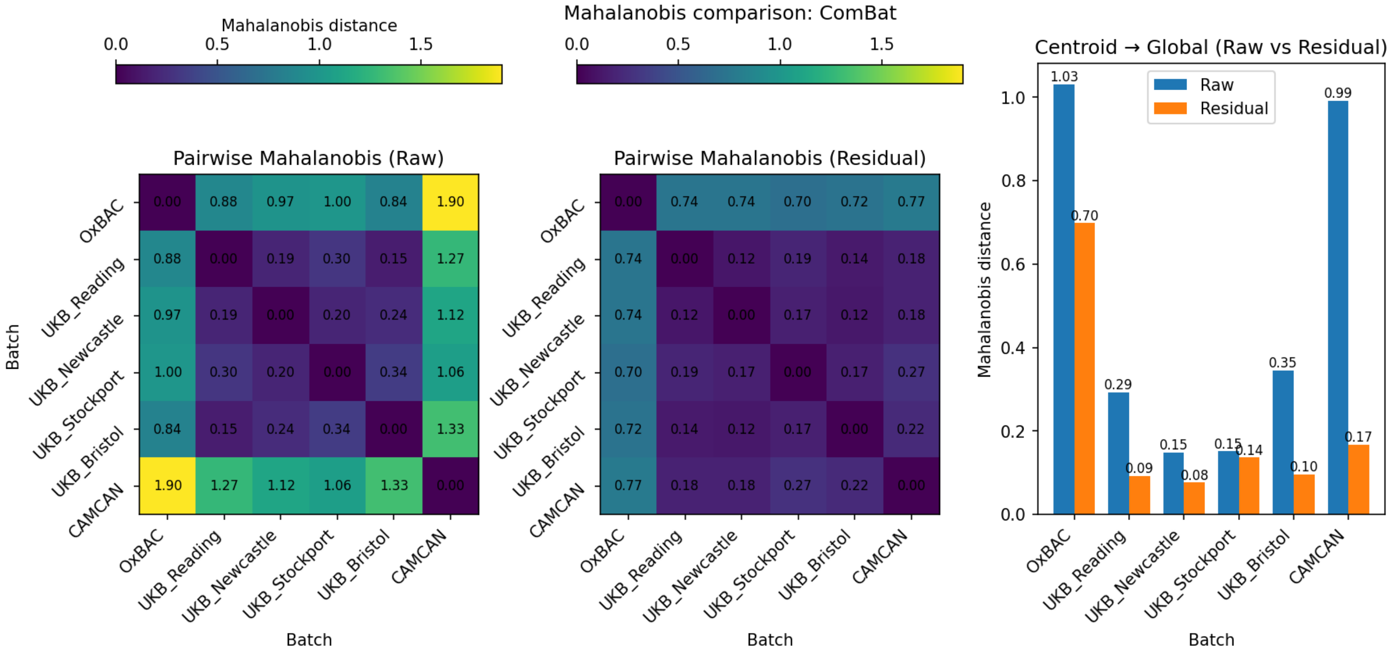


Comparison: Mahalanobis (ComBat)

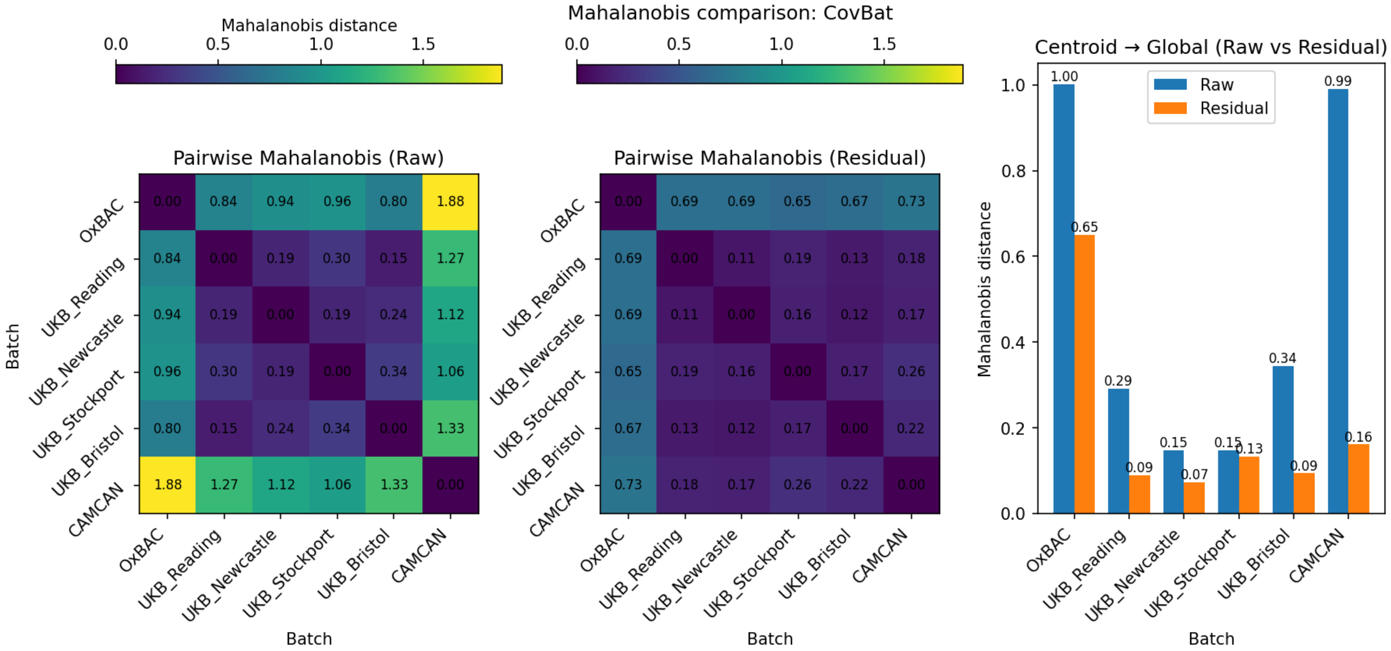


Comparison: Mahalanobis (CovBat)

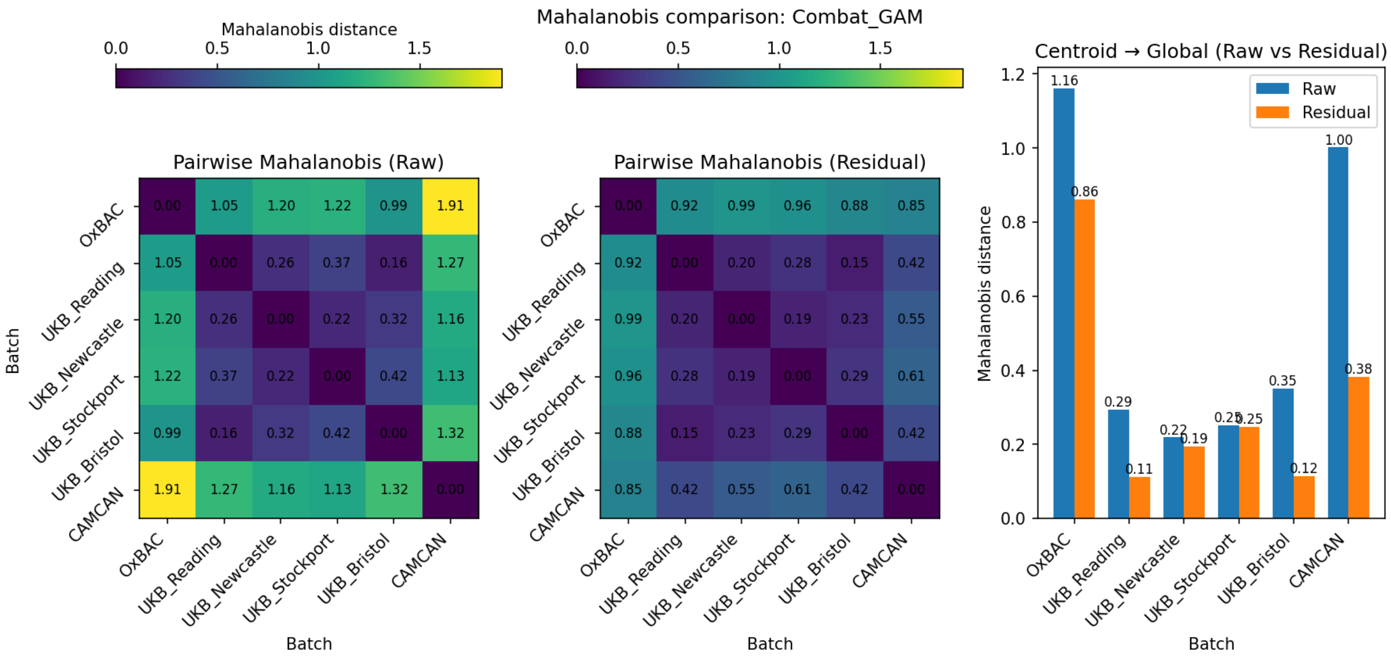


Comparison: Mahalanobis (Combat_GAM)

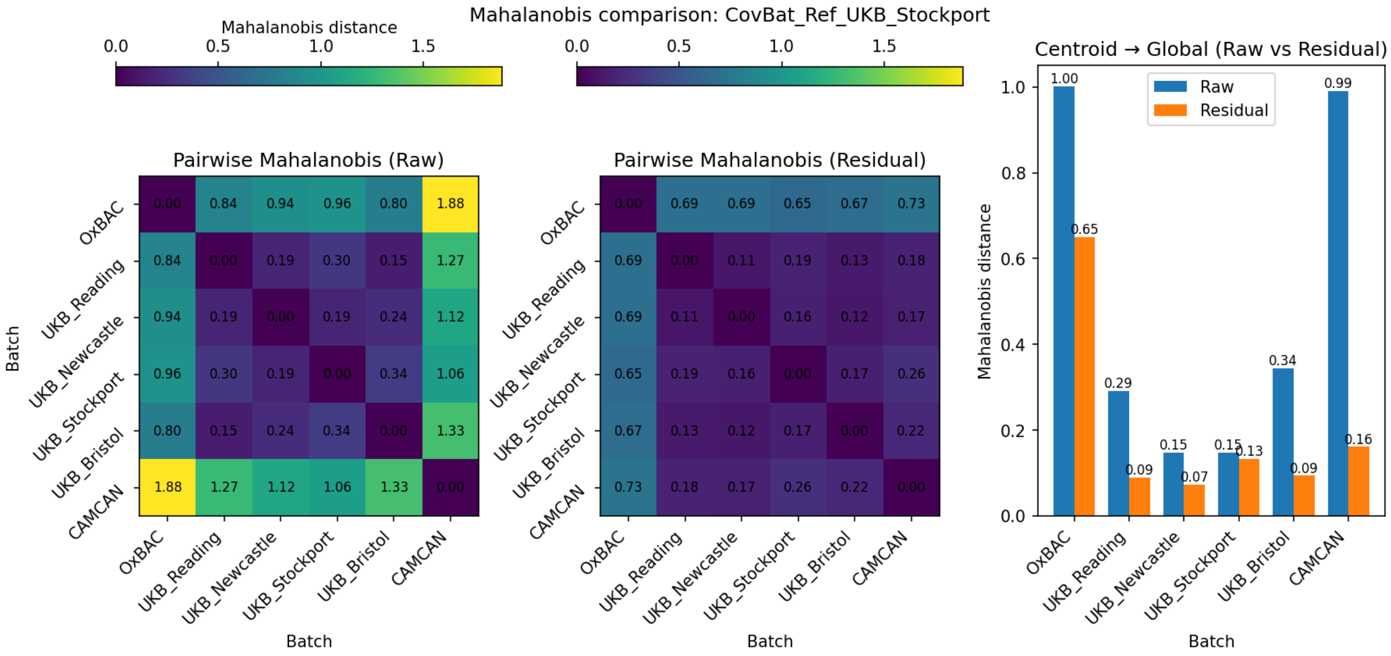


Comparison: Mahalanobis (CovBat_Ref_UKB_Stockport)

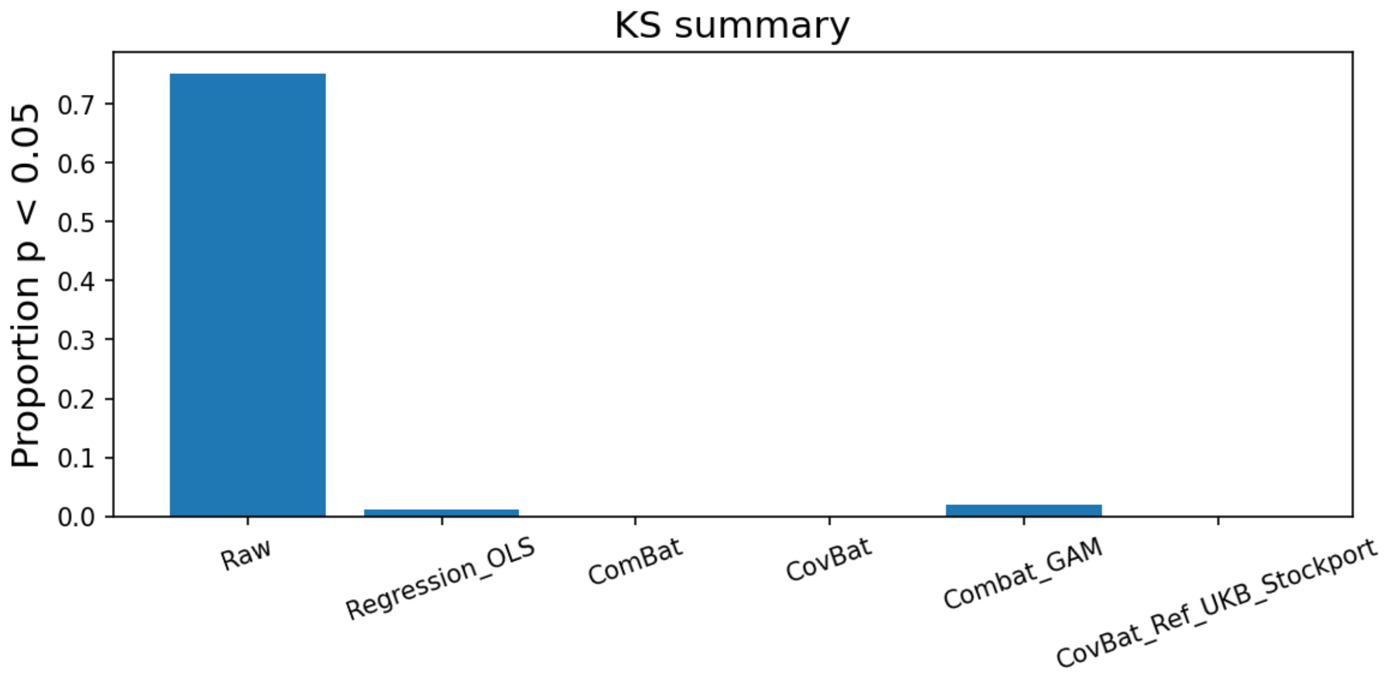


Comparison: KS summary

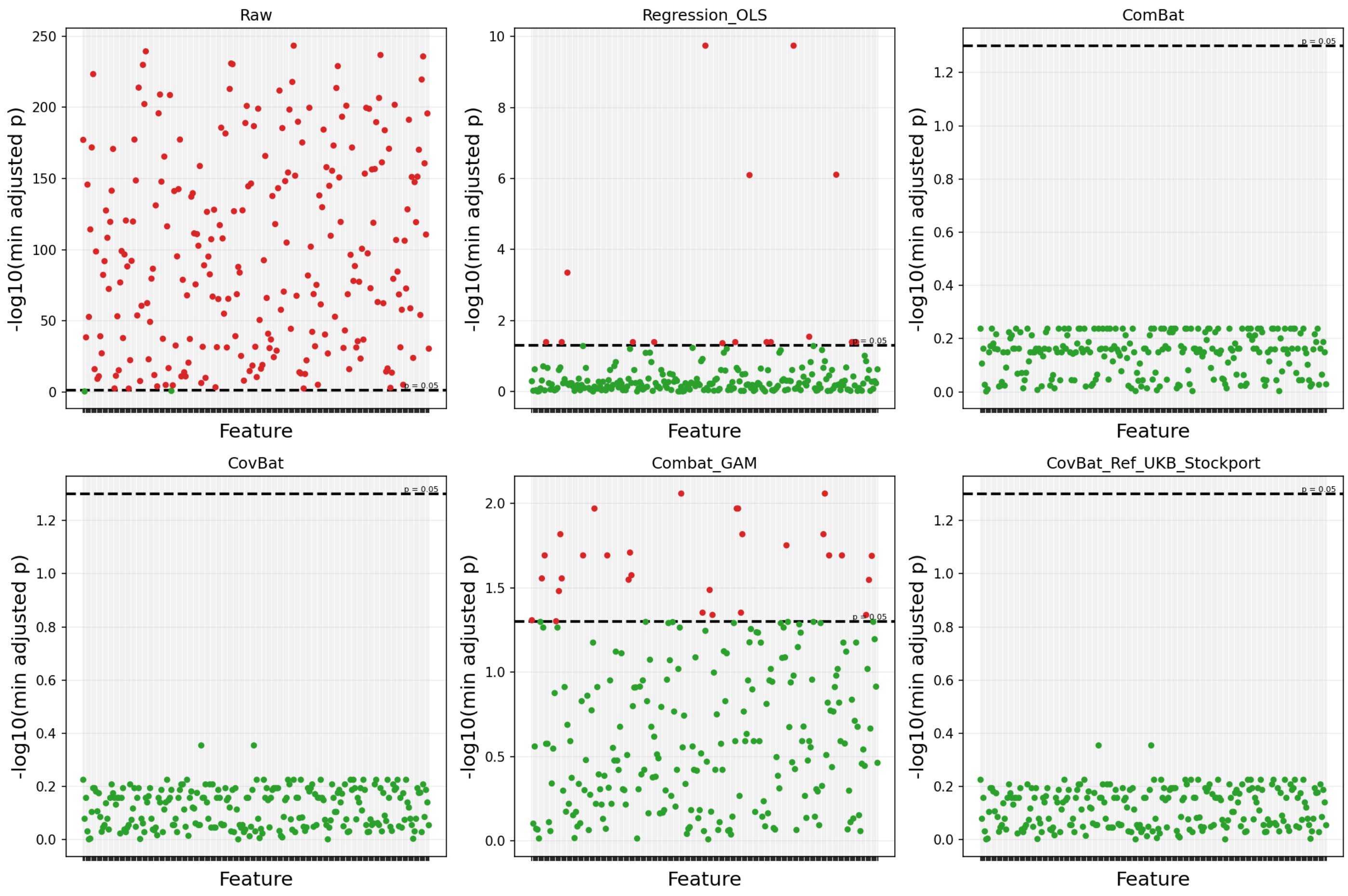


Comparison: KS feature-wise

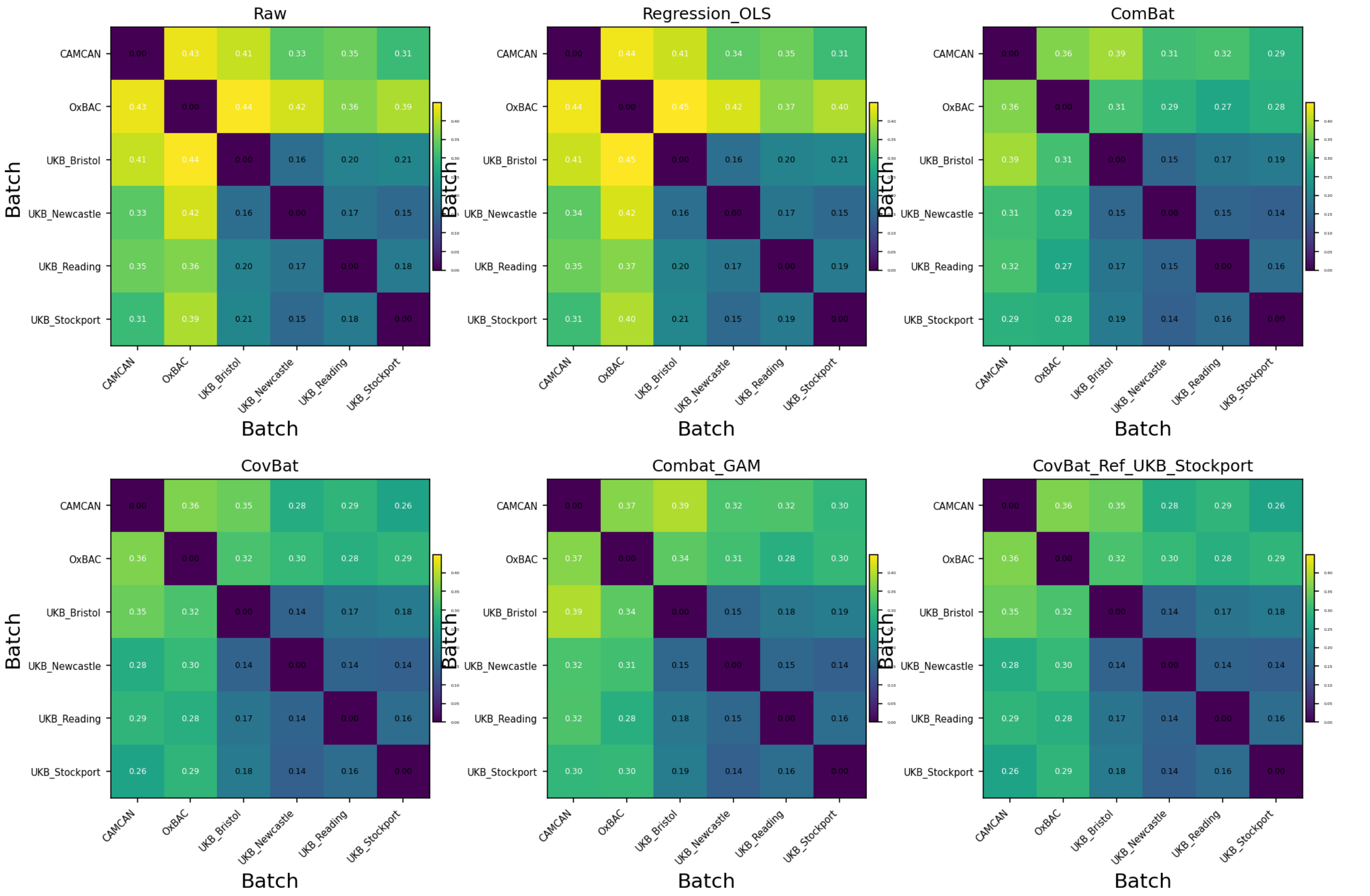


Comparison: covariance Frobenius

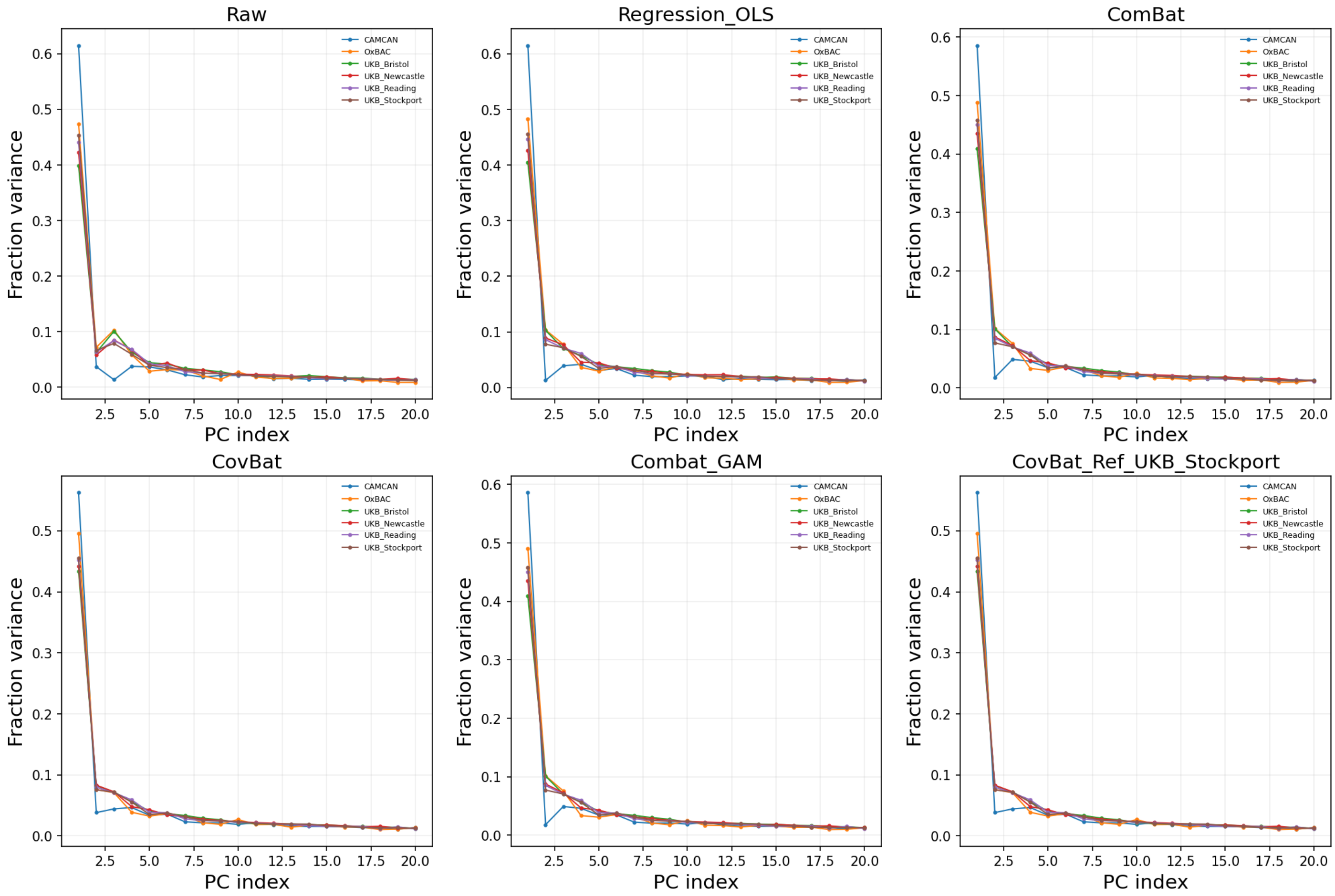

Comparison: batch scree plots

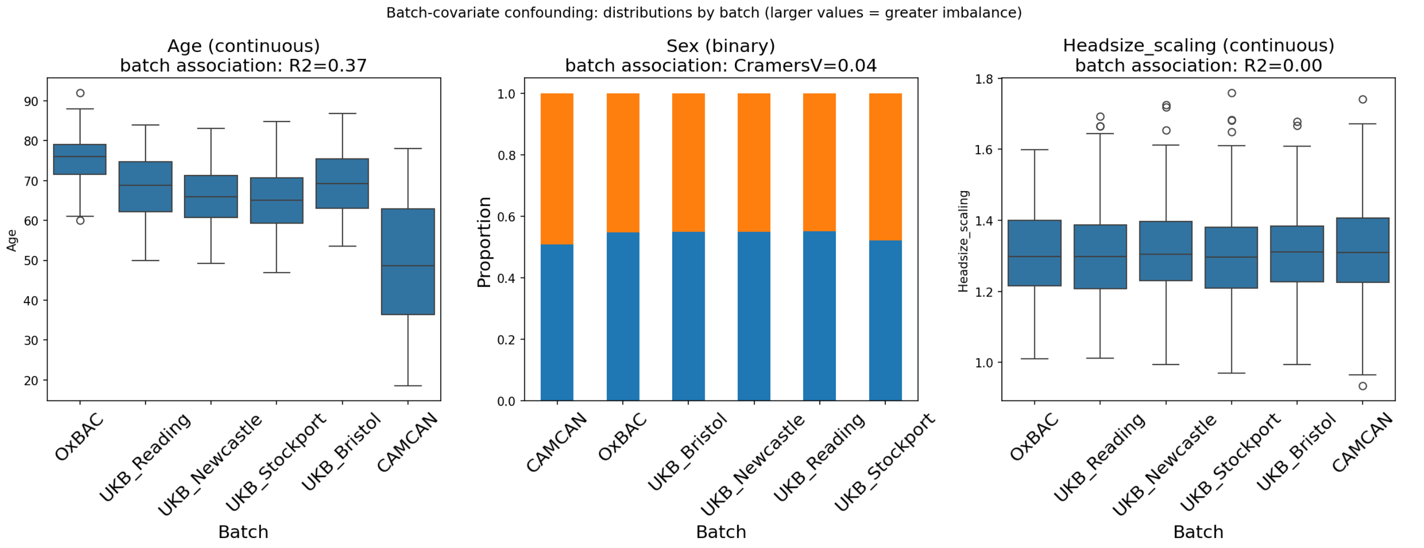


Batch-covariate confounding: distributions by batch

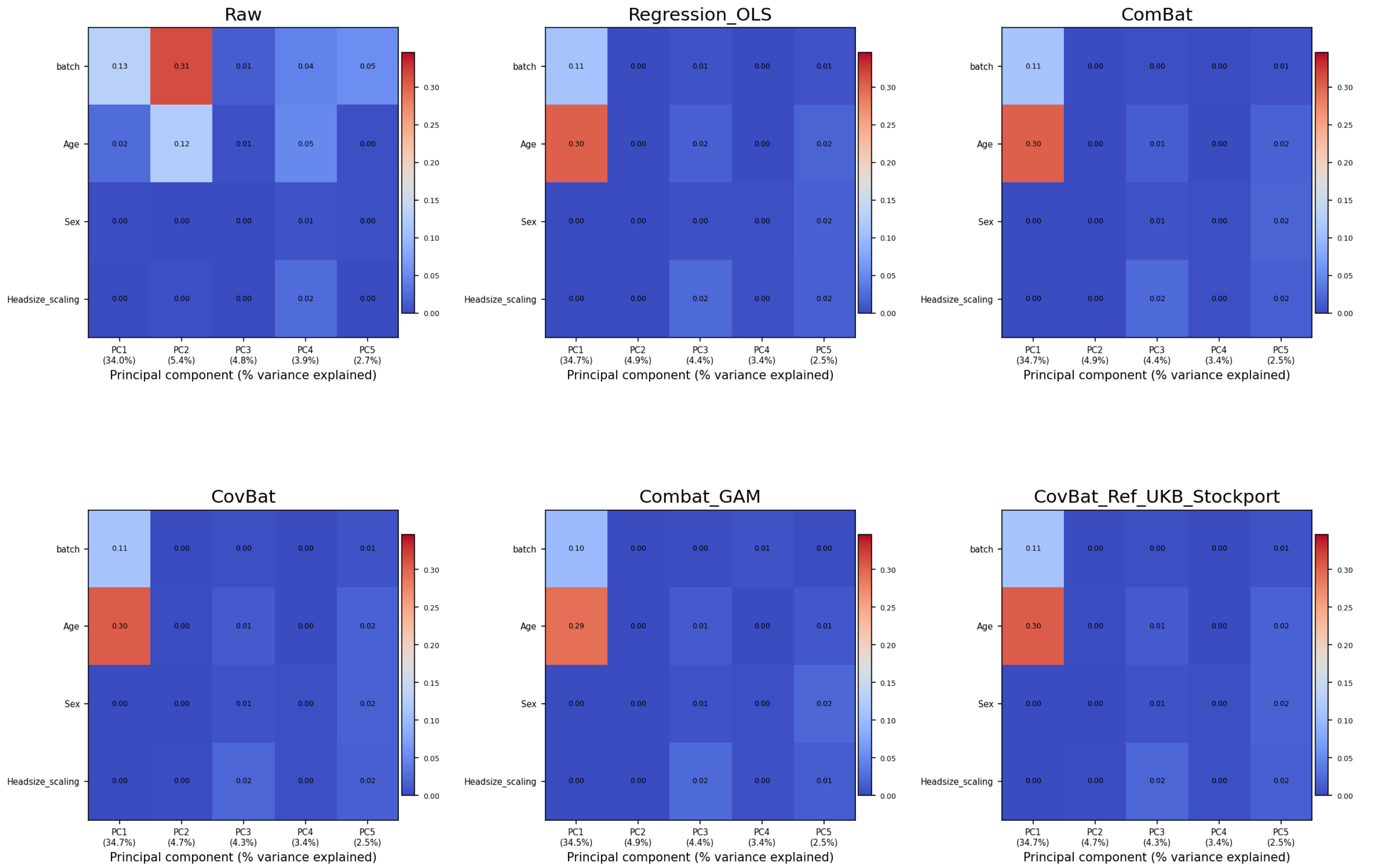


Comparison: variance associated with metadata (Omnibus R2)

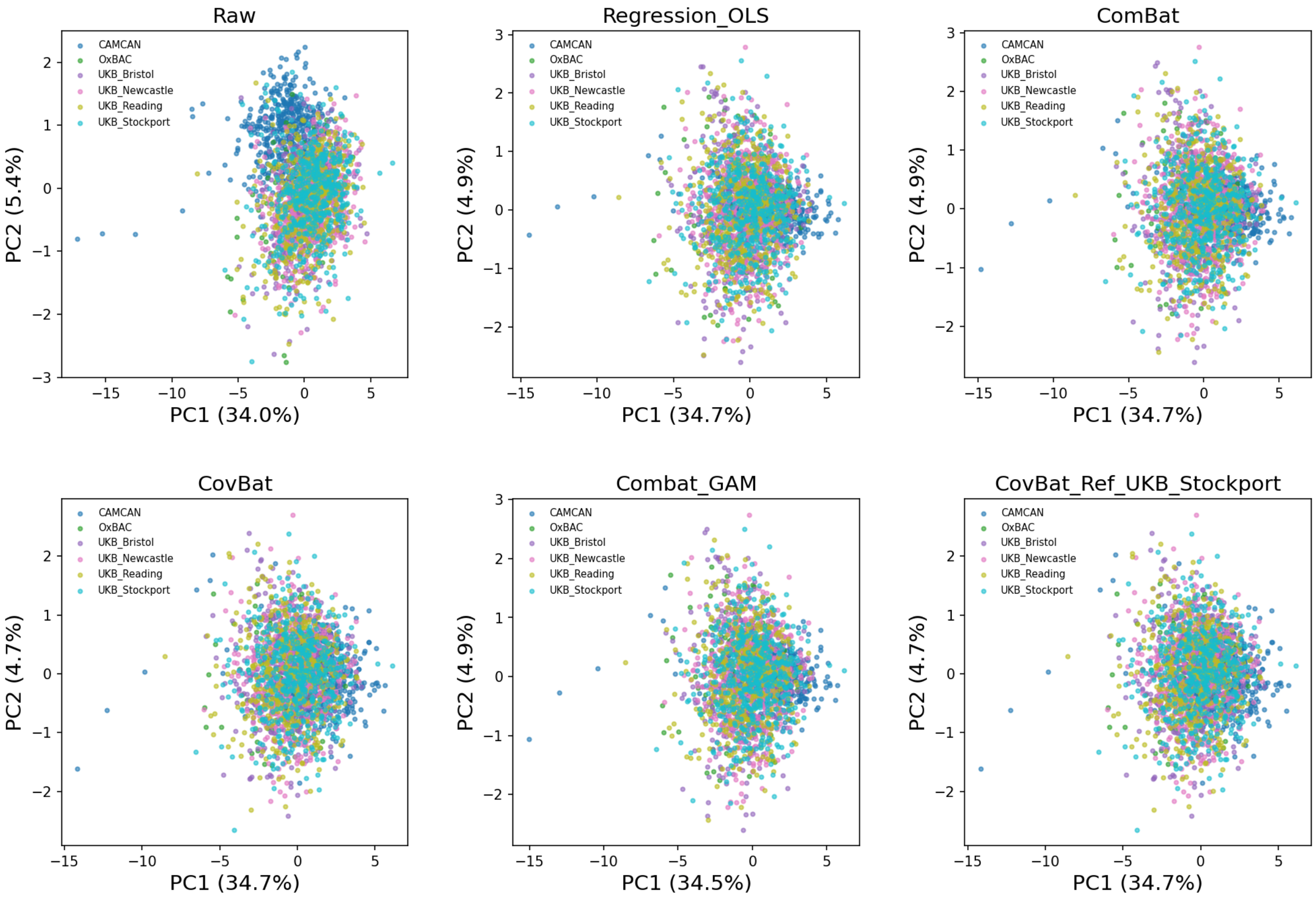


Comparison: PCA embeddings

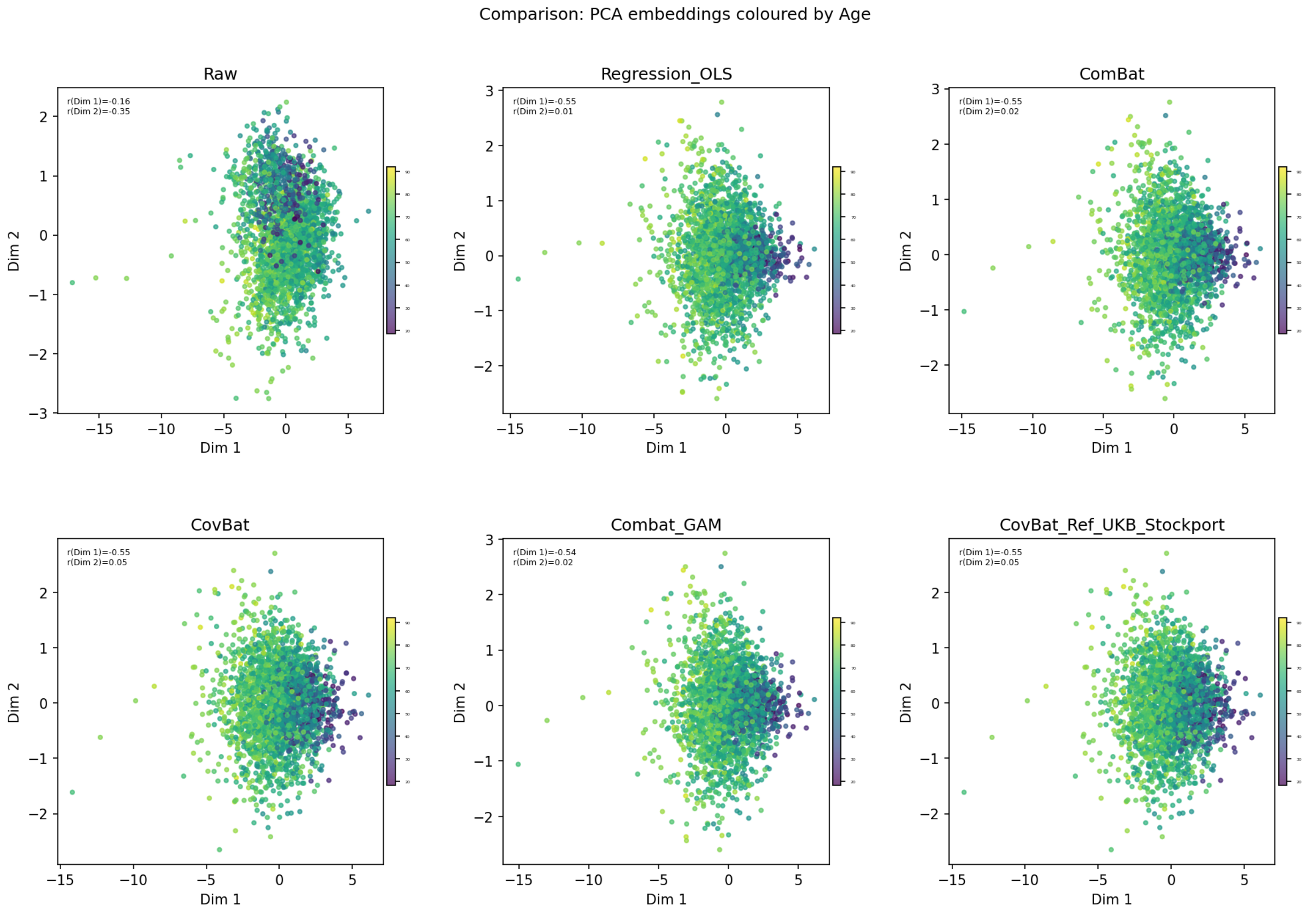


Comparison: PCA embeddings coloured by Age

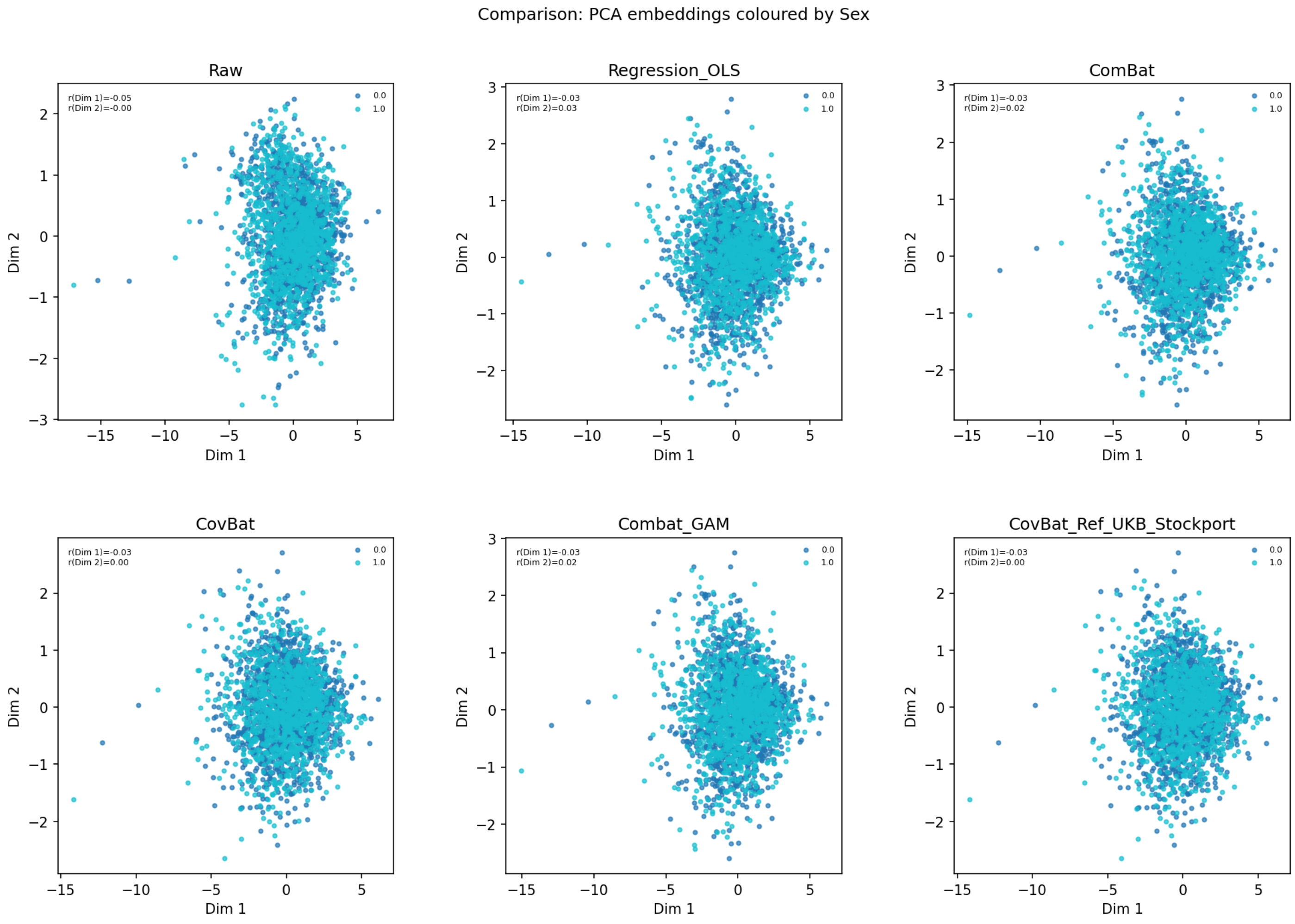


Comparison: PCA embeddings coloured by Sex

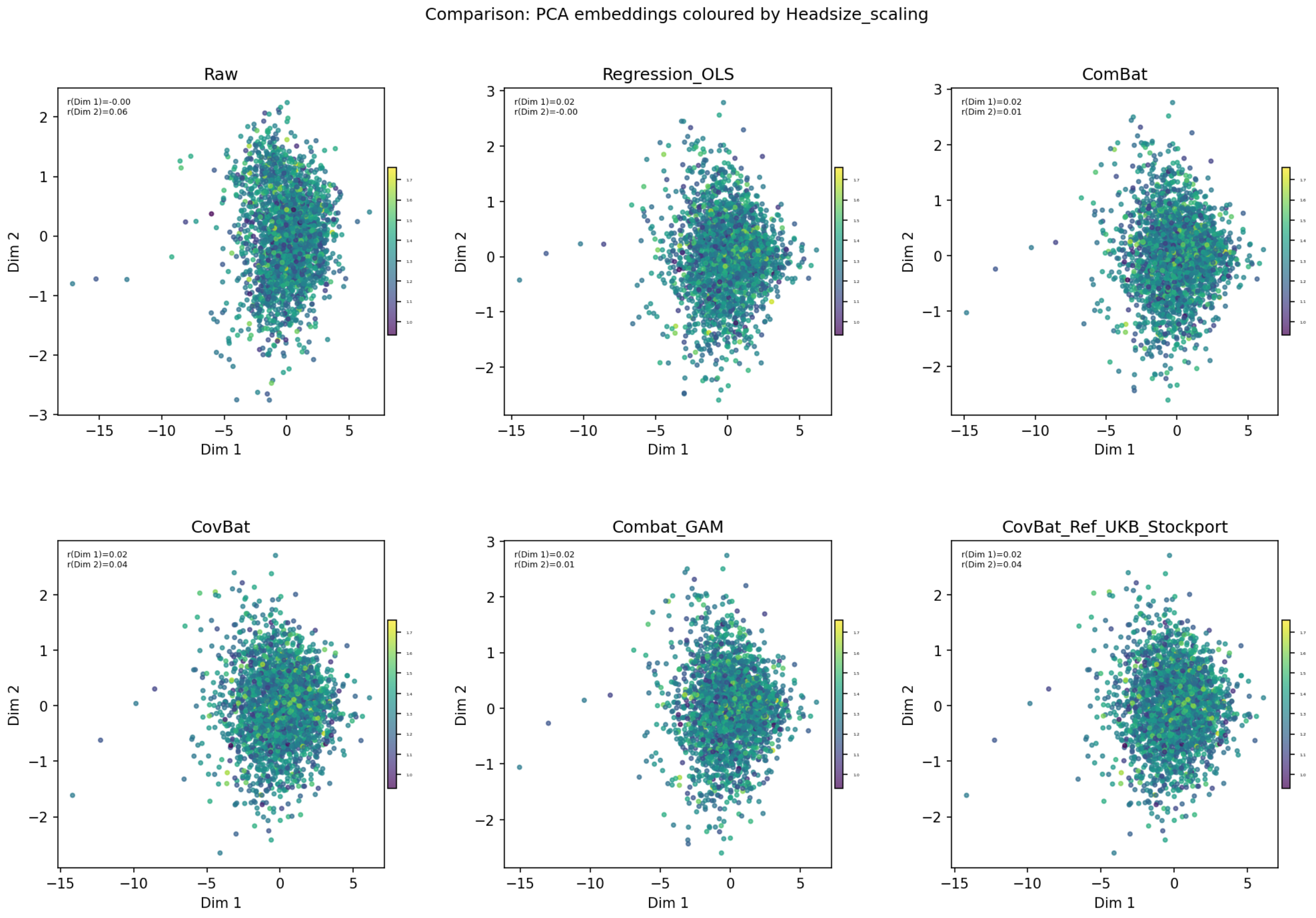


Comparison: PCA embeddings coloured by Headsize_scaling

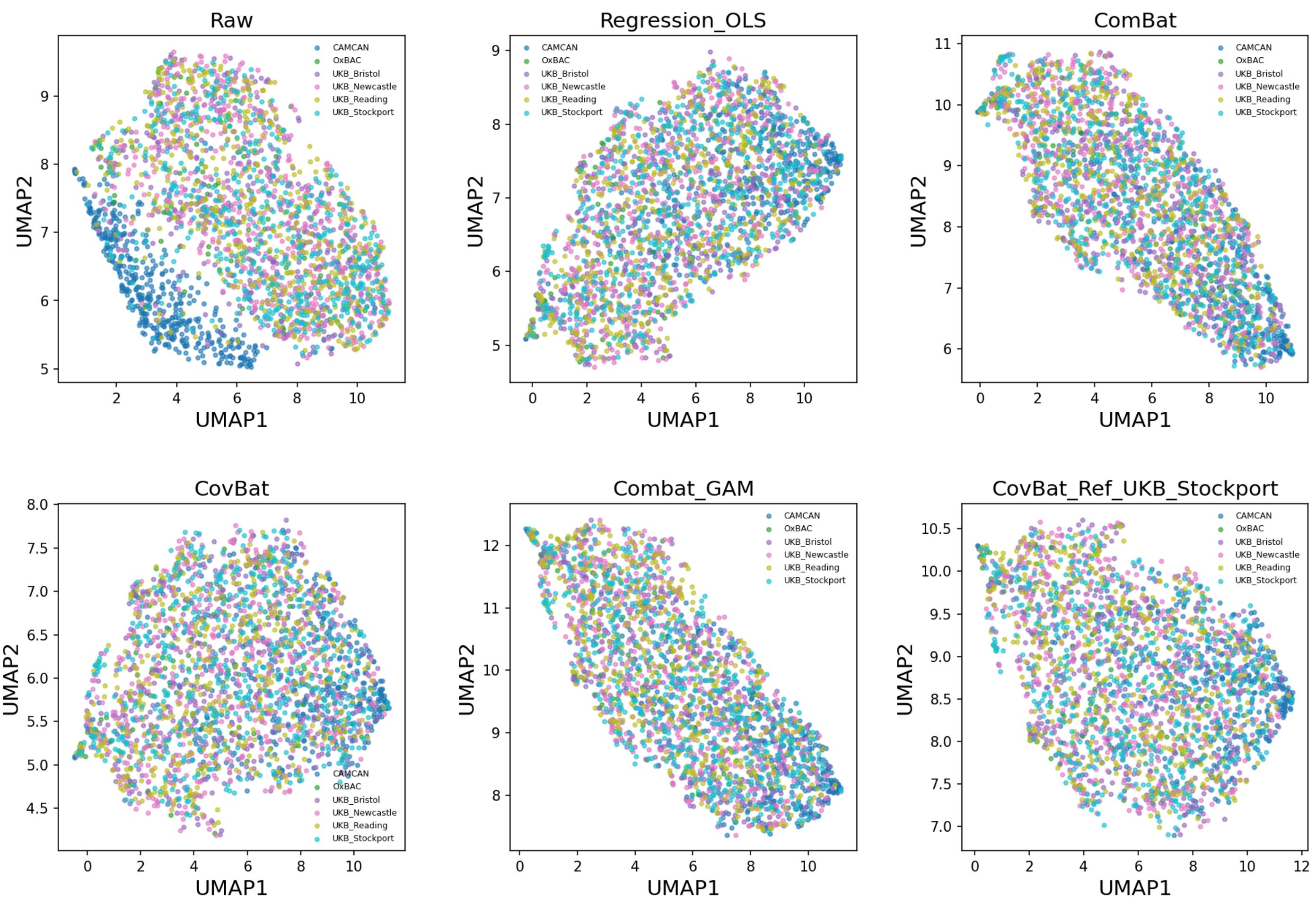


Comparison: UMAP embeddings

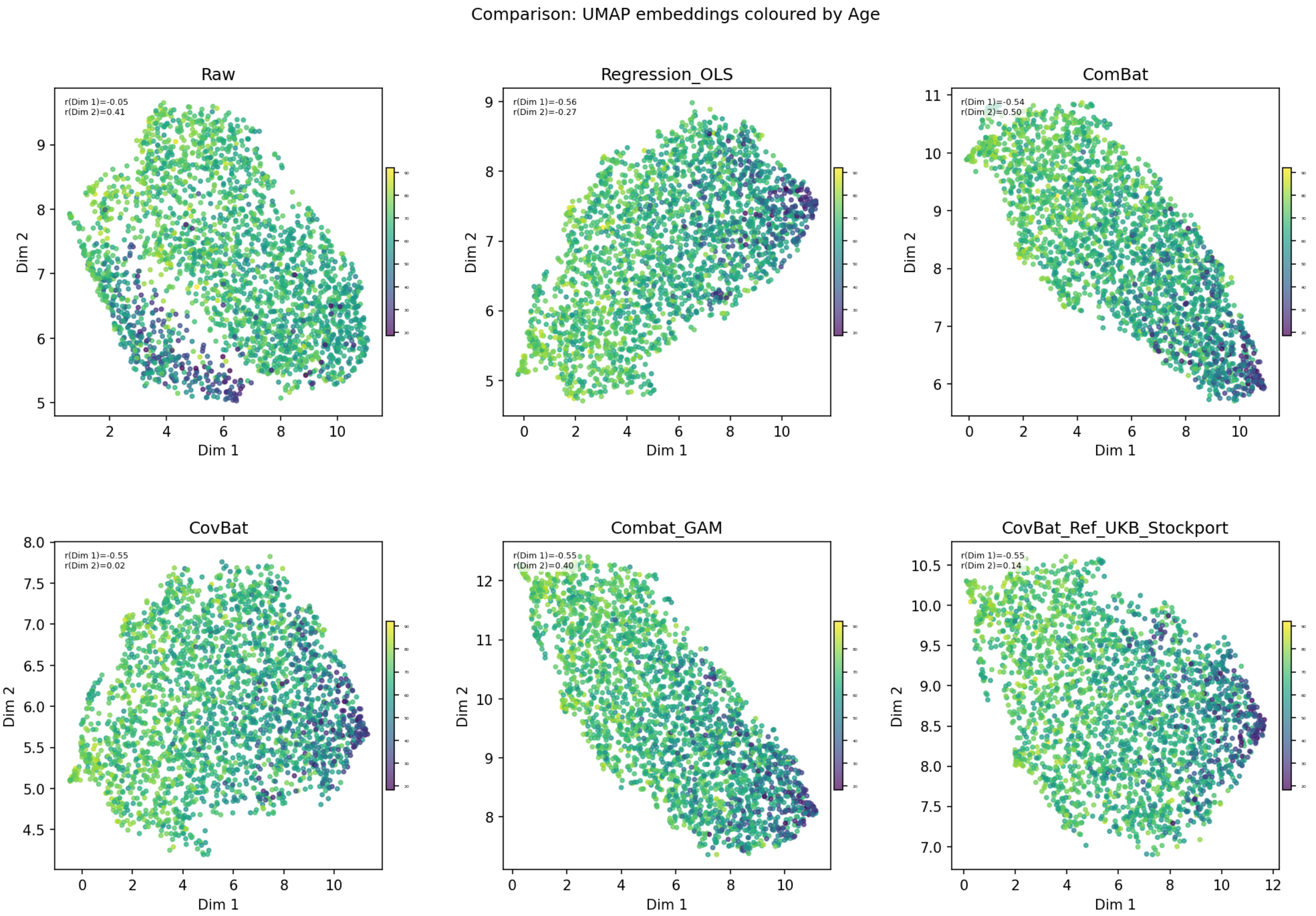


Comparison: UMAP embeddings coloured by Age

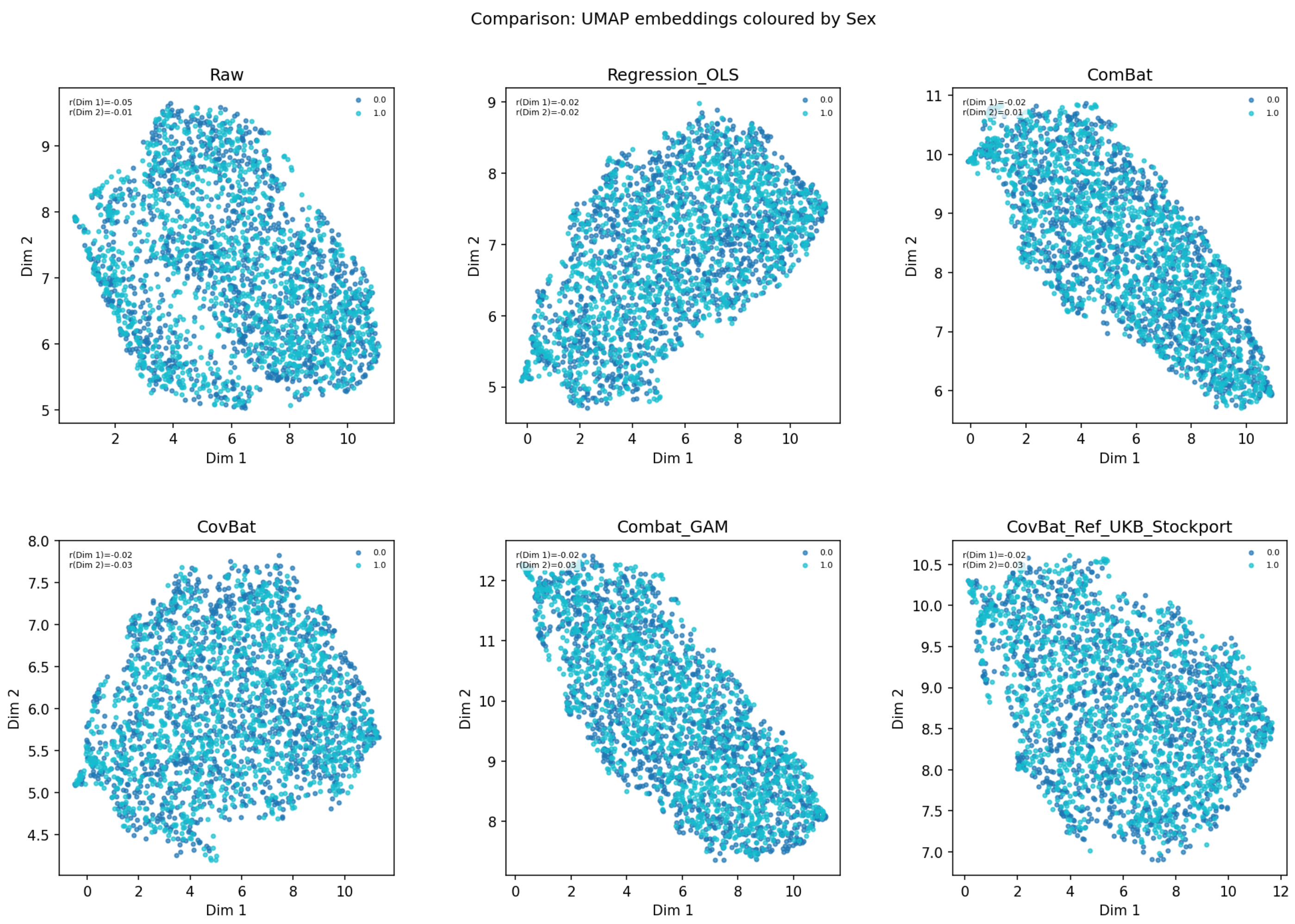


Comparison: UMAP embeddings coloured by Sex

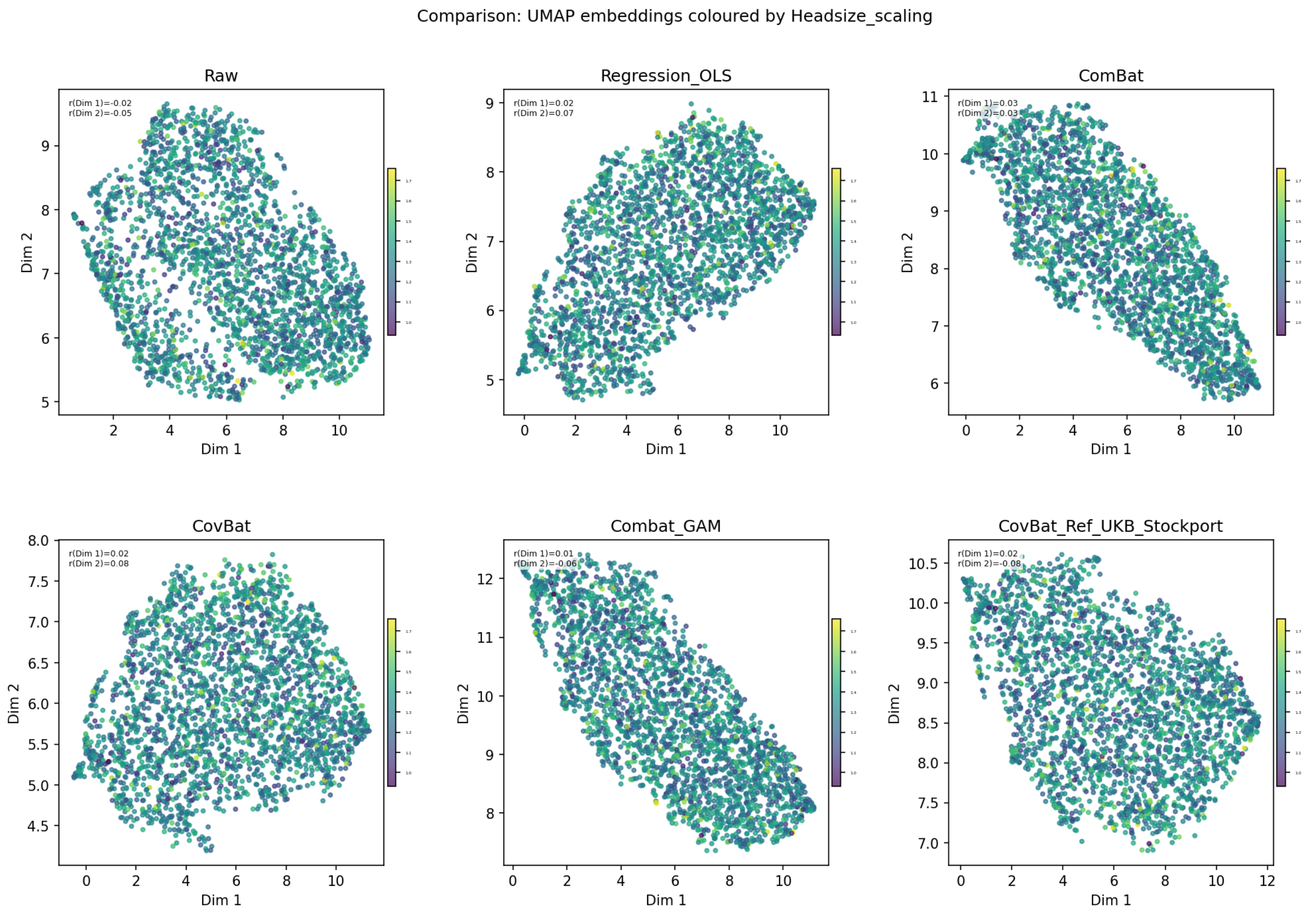


Comparison: UMAP embeddings coloured by Headsize_scaling

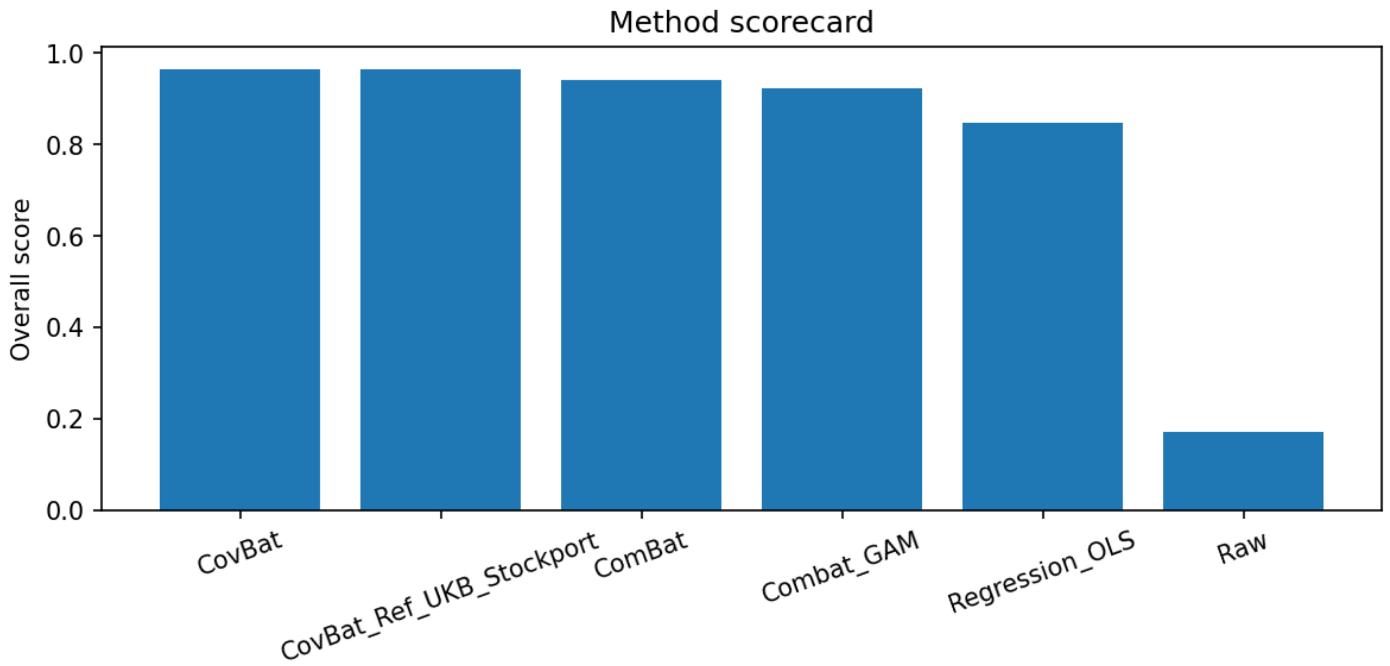


Comparison: method scorecard

### 5 Supplementary information for the longitudinal demonstration

The longitudinal report was generated after applying ComBat, CovBat and longitudinal ComBat within DHARM. All data were loaded from a CSV file containing Imaging Derived Phenotypes from the ON-Harmony study available on OpenNeuro (https://openneuro.org/datasets/ds004712/versions/2.0.1) (Warrington et al., 2023). The covariates were given to DHARM when running the longitudinal reports as a dictionary, but are required to be arrays or data-frames for the harmonisation methods. DHARM does allow for arrays and data-frames to be used when running the code as of version 1.1.8, but previous versions require covariates to be supplied as a dictionary.

```
covariate_dict = {
    name: covariate_array[:, i]
    for i, name in enumerate(covariate_names)
}

df_covariates_with_timepoint = df_covariates.copy()
df_covariates_with_timepoint["timepoint"] = timepoint

# Ensure all covariate columns are numeric where applicable
df_covariates = df2[["subject", "age", "sex"]].copy()
df_covariates_no_subject = df_covariates.drop(columns=["subject"])

scanner_array = (
    df["scan_session"]
    .astype(str)
    .str.strip()
    .str.replace(r"\d+$", "", regex=True)
    .tolist()
)

df_covariates_with_timepoint = df_covariates.copy()
df_covariates_with_timepoint["timepoint"] = timepoint

# Ensure all covariate columns are numeric where applicable
df_covariates = df2[["subject", "age", "sex"]].copy()
df_covariates_no_subject = df_covariates.drop(columns=["subject"])

######## RUN HARMONISATION METHODS ########

# Combat harmonisation
harmonised_data1 = HF.combat_modular(df[features],batch_array,mod=df_covariates_no_su
combat_IDP       = harmonised_data1["bayesdata"]

# CovBat harmonisation
harmonised_data2 = HF.combat_modular(df[features],batch_array,mod=df_covariates_no_su
covbat_IDP       = harmonised_data2["bayesdata"]

# Longitudinal combat with no timepoint effect and only subject random effect: (1|sub
harmonised_data3 = HF.long_combat(df[features], batch_array, model_inputs=df_covariate
Lcombat_noTP_IDP = harmonised_data3["Bayesdata"]


# Individual dataset reports can be run as follows:

DiagnosticReport.LongitudinalReport(data=data_array,
                                    subject_ids=subject_array,
                                    batch=batch_array,
                                    timepoints=timepoint,
```

```
                                          features=features,
                                          covariates=covariate_dict,
                                          SaveArtifacts=False,
                                          save_dir="Raw_report")

DiagnosticReport.LongitudinalReport(data=combat_IDP,
                                    subject_ids=subject_array,
                                    batch=batch_array,
                                    timepoints=timepoint,
                                    features=features,
                                    covariates=covariate_dict,
                                    SaveArtifacts=False,
                                    save_dir="combat_report")

DiagnosticReport.LongitudinalReport(data=covbat_IDP,
                                    subject_ids=subject_array,
                                    batch=batch_array,
                                    timepoints=timepoint,
                                    features=features,
                                    covariates=covariate_dict,
                                    SaveArtifacts=False,
                                    save_dir="covbat_report")

DiagnosticReport.LongitudinalReport(data=Lcombat_noTP_IDP,
                                    subject_ids=subject_array,
                                    batch=batch_array,
                                    timepoints=timepoint,
                                    features=features,
                                    covariates=covariate_dict,
                                    SaveArtifacts=False,
                                    save_dir="Lcombat_report")


#############################################################

######## RUN COMPARISON REPORT ########

#############################################################

datasets = {
    "Raw": data_array,
    "ComBat": combat_IDP,
    "CovBat": covbat_IDP,
    "LongComBat": Lcombat_noTP_IDP
}


comparison_report = (
    DiagnosticReport.LongitudinalComparisonReport(
        datasets=datasets,
        batch=batch_array,
        subject_ids=subject_array,
        #timepoints=timepoint,
        covariates=covariate_dict,
        covariate_names=list(
            covariate_dict.keys()
        ),
        features=features,
```

```
            save_data=True,
            save_dir="ONHarmony_OnlyBetweenScanner_Comparison_report",
            report_name="ONHarmony_OnlyBetweenScanner_Comparison_report",
            include_raw=True,
            raw_name="Raw",
            SaveArtifacts=False,
            show=False,
            max_cols_in_plotting_grid=2,
            height_in_plotting_grid=6,
            width_in_plotting_grid=8,

    )
)
```

##### 5.1 Longitudinal comparison report

### Longitudinal comparison of harmonisation methods

#### Contents


- Report preamble and overview
- Multi-method longitudinal comparison overview
- Batch-covariate confounding diagnostics
- Diagnostics for Raw
- Diagnostics for ComBat
- Diagnostics for CovBat
- Diagnostics for LongComBat
- Longitudinal method scorecard
- Best method summary


```
2026-10-20 10:22:59,613 — INFO — === Report session started ===
2026-10-20 10:22:59,613 — INFO — save_artifacts=False | save_dir=None
2026-10-20 10:22:59,615 — INFO — initialised HTML report at:
ONHarmony_OnlyBetweenScanner_Comparison_report/ONHarmony_OnlyBetweenScanner_Comparison_report_2026-10-20_10-22-59.html
2026-10-20 10:22:59,615 — INFO — [TITLE] Longitudinal comparison of harmonisation methods
```

#### Report preamble and overview

```
2026-10-20 10:22:59,615 — INFO — [SECTION] Report preamble and overview
This report compares multiple harmonisation methods using longitudinal diagnostics in the same execution order as the longitudinal single
Ranking prioritises subject stability, followed by residual batch effects and biological preservation.
```

#### Multi-method longitudinal comparison overview

```
2026-10-20 10:22:59,615 — INFO — [SECTION] Multi-method longitudinal comparison overview
Comparison dataset characteristics:
Validated methods: Raw, ComBat, CovBat, LongComBat
Total samples: 120
Features: 138
Unique subjects: 20
Unique timepoints: 0
Samples in each batch: {np.str_('NOT1ACH'): np.int64(20), np.str_('NOT2ING'): np.int64(20), np.str_('NOT3GEM'): np.int64(10), np.str_('NO
Covariates: age, sex
Missing data per method: Raw=0, ComBat=0, CovBat=0, LongComBat=0
```

#### Batch-covariate confounding diagnostics

```
2026-10-20 10:22:59,615 — INFO — [SECTION] Batch-covariate confounding diagnostics
Batch-covariate confounding quantifies potential imbalance of each covariate across batches,
 Continuous covariates are summarised by the omnibus R2 from covariate ~ batch; binary/categorical
covariates, including timepoint, are summarised by Cramer's V. Batch is always treated as nominal
categorical regardless of its input encoding. Larger values indicate greater covariate imbalance
across batches and therefore greater potential for confounding.

Variable       Type Statistic  Value
     age continuous        R2  0.009
     sex     binary  CramersV  0.182
```

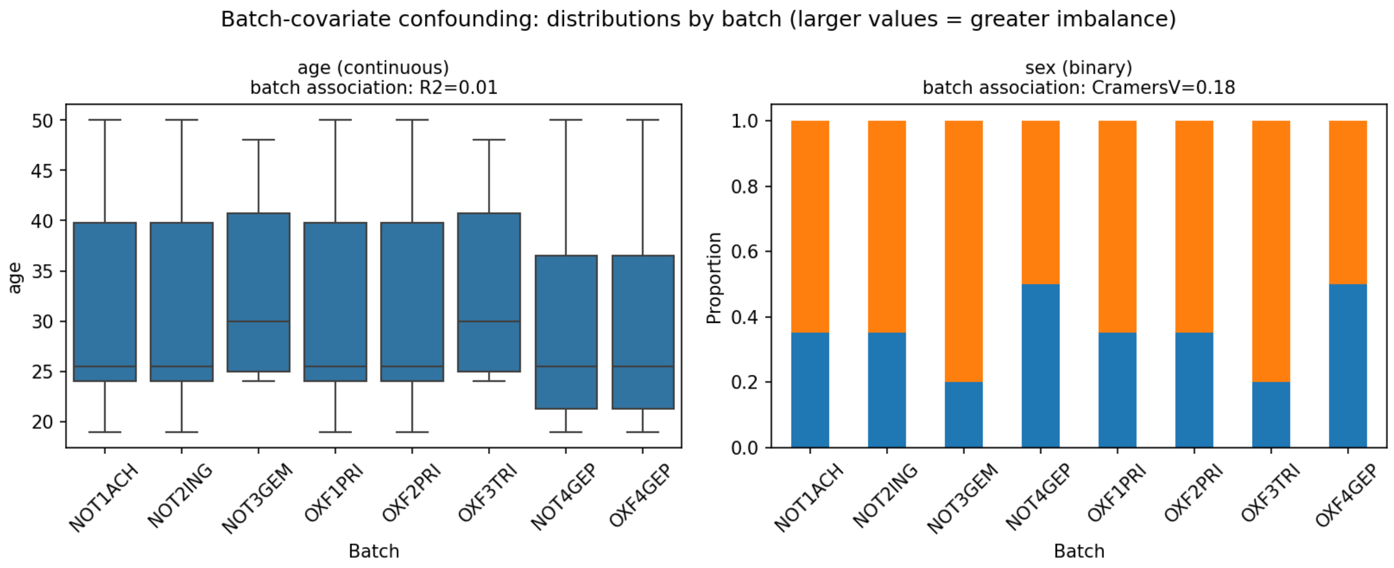


Batch-covariate confounding: distributions by batch

#### Diagnostics for Raw

```
2026-10-20 10:22:59,916 — INFO — [SECTION] Diagnostics for Raw
2026-10-20 10:22:59,916 — INFO — Running longitudinal diagnostics for method: Raw
Warnings for Raw:
- MultiVariateBatchDifference_long skipped because the analysis covariance matrix is rank deficient (112 of 138).
Unavailable diagnostics for Raw:
- subject_order: missing_timepoint (Subject-order analysis requires at least two known timepoints.)
Summary metrics for Raw:
- median_spearman_rho: nan
- prop_significant_spearman: nan
- median_within_subject_variability: 5.948736641417044
- median_icc: 0.8256553008389544
- prop_high_icc: 0.717391304347826
- mean_n_is_batchSig: 10.978260869565217
- prop_additive_significant: 0.9855072463768116
- prop_multiplicative_significant: 0.18115942028985507
- mean_mahalanobis_distance: nan
- biological_signal_score: 0.10869565217391304
----------------------------------------------------------------------------------------------------------------------------------------------
```

#### Diagnostics for ComBat

```
2026-10-20 10:23:10,516 — INFO — [SECTION] Diagnostics for ComBat
2026-10-20 10:23:10,516 — INFO — Running longitudinal diagnostics for method: ComBat
Warnings for ComBat:
- MultiVariateBatchDifference_long skipped because the analysis covariance matrix is rank deficient (112 of 138).
Unavailable diagnostics for ComBat:
- subject_order: missing_timepoint (Subject-order analysis requires at least two known timepoints.)
Summary metrics for ComBat:
- median_spearman_rho: nan
- prop_significant_spearman: nan
- median_within_subject_variability: 4.4557601238987035
- median_icc: 0.9044471504250433
- prop_high_icc: 0.9855072463768116
- mean_n_is_batchSig: 3.8115942028985508
- prop_additive_significant: 0.5217391304347826
- prop_multiplicative_significant: 0.014492753623188406
- mean_mahalanobis_distance: nan
- biological_signal_score: 0.11956521739130435
----------------------------------------------------------------------------------------------------------------------------------------------
```

#### Diagnostics for CovBat

```
2026-10-20 10:23:21,251 — INFO — [SECTION] Diagnostics for CovBat
2026-10-20 10:23:21,251 — INFO — Running longitudinal diagnostics for method: CovBat
Warnings for CovBat:
- MultiVariateBatchDifference_long skipped because the analysis covariance matrix is rank deficient (112 of 138).
Unavailable diagnostics for CovBat:
- subject_order: missing_timepoint (Subject-order analysis requires at least two known timepoints.)
Summary metrics for CovBat:
- median_spearman_rho: nan
- prop_significant_spearman: nan
- median_within_subject_variability: 4.378764281373105
```

```
- median_icc: 0.9036397235181104
- prop_high_icc: 1.0
- mean_n_is_batchSig: 3.7318840579710146
- prop_additive_significant: 0.42028985507246375
- prop_multiplicative_significant: 0.021739130434782608
- mean_mahalanobis_distance: nan
- biological_signal_score: 0.12318840579710146
--------------------------------------------------------------------------------------------------------------------------
```

#### Diagnostics for LongComBat

```
2026-10-20 10:23:32,123 — INFO — [SECTION] Diagnostics for LongComBat
2026-10-20 10:23:32,123 — INFO — Running longitudinal diagnostics for method: LongComBat
Warnings for LongComBat:
- MultiVariateBatchDifference_long skipped because the analysis covariance matrix is rank deficient (112 of 138).
Unavailable diagnostics for LongComBat:
- subject_order: missing_timepoint (Subject-order analysis requires at least two known timepoints.)
Summary metrics for LongComBat:
- median_spearman_rho: nan
- prop_significant_spearman: nan
- median_within_subject_variability: 4.701947072411114
- median_icc: 0.9033598735285062
- prop_high_icc: 0.9347826086956522
- mean_n_is_batchSig: 0.0
- prop_additive_significant: 0.0
- prop_multiplicative_significant: 0.0
- mean_mahalanobis_distance: nan
- biological_signal_score: 0.10869565217391304
--------------------------------------------------------------------------------------------------------------------------
```

#### Longitudinal method scorecard

```
2026-10-20 10:23:42,834 — INFO — [SECTION] Longitudinal method scorecard
Longitudinal scorecard summary:
    method  subject_stability_score  batch_removal_score  biological_preservation_score  overall_score  overall_rank diagnostics_availabl
    CovBat                 0.996584             0.704532                           1.00       0.909481           1.0                    4/
    ComBat                 0.966558             0.681131                           0.75       0.848446           2.0                    4/
LongComBat                 0.849860             1.000000                           0.00       0.767423           3.0                    4/
       Raw                 0.000000             0.000000                           0.00       0.000000           4.0                    4/
```

#### Best method summary

```
2026-10-20 10:23:42,834 — INFO — [SECTION] Best method summary
Best overall method: CovBat
Best subject stability: CovBat
Best batch removal: LongComBat
Best biological preservation: CovBat
Best subject_order: Not computed
Best within_subject_variability: CovBat
Best icc: ComBat
Best pairwise_batch_residual: LongComBat
Best additive_batch_residual: LongComBat
Best multiplicative_batch_residual: LongComBat
Best multivariate_batch_residual: Not computed
Best biological_signal: CovBat
Best overall method is CovBat. Longitudinal diagnostics show trade-offs across subject stability, batch removal, and biological preservat
Saved scorecard CSV: ONHarmony_OnlyBetweenScanner_Comparison_report/ONHarmony_OnlyBetweenScanner_Comparison_report_2026-10-20/Longitudina
2026-10-20 10:23:53,538 — INFO — === Report session finished ===
```

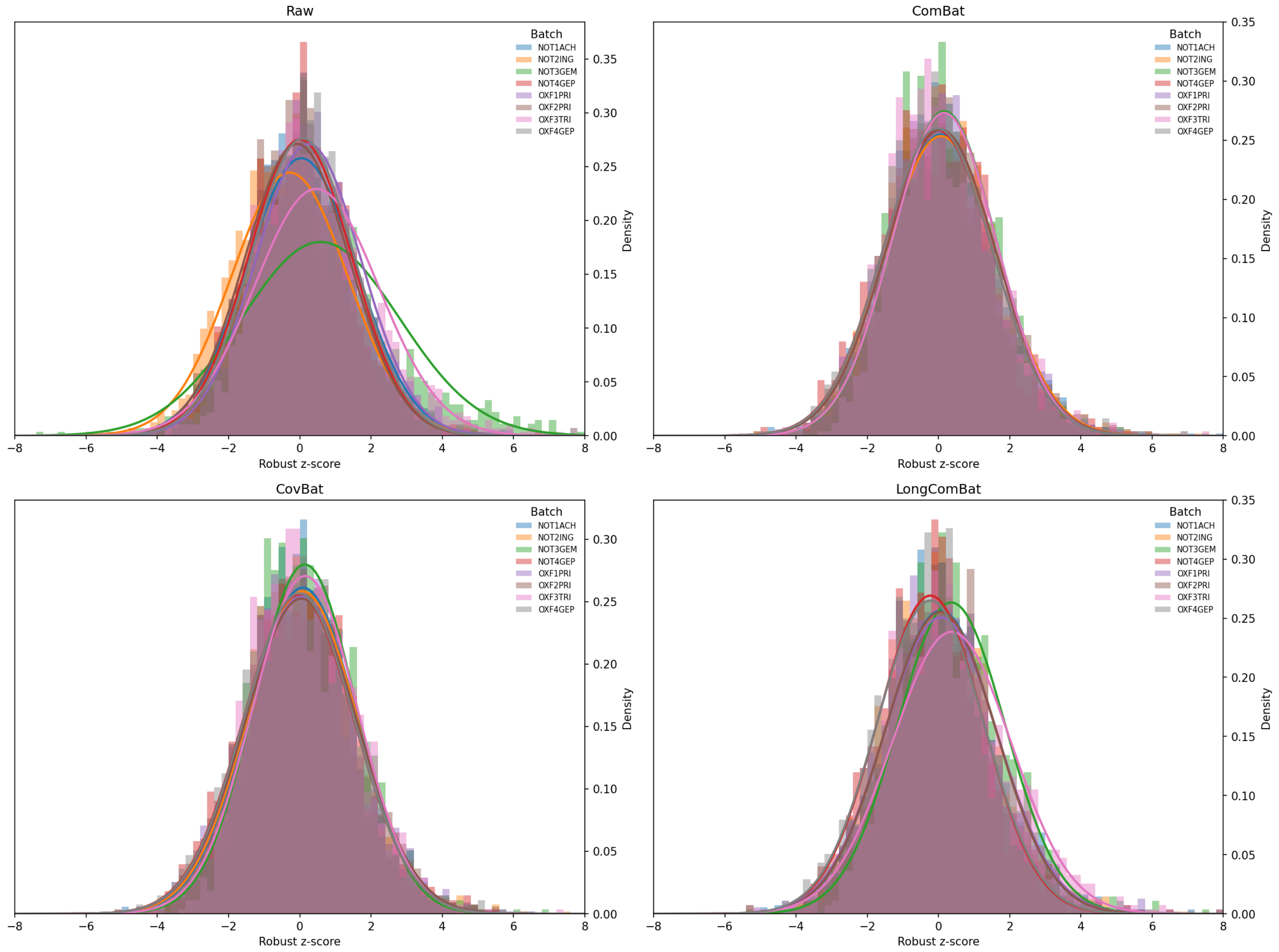


Longitudinal comparison: unadjusted z-score distributions

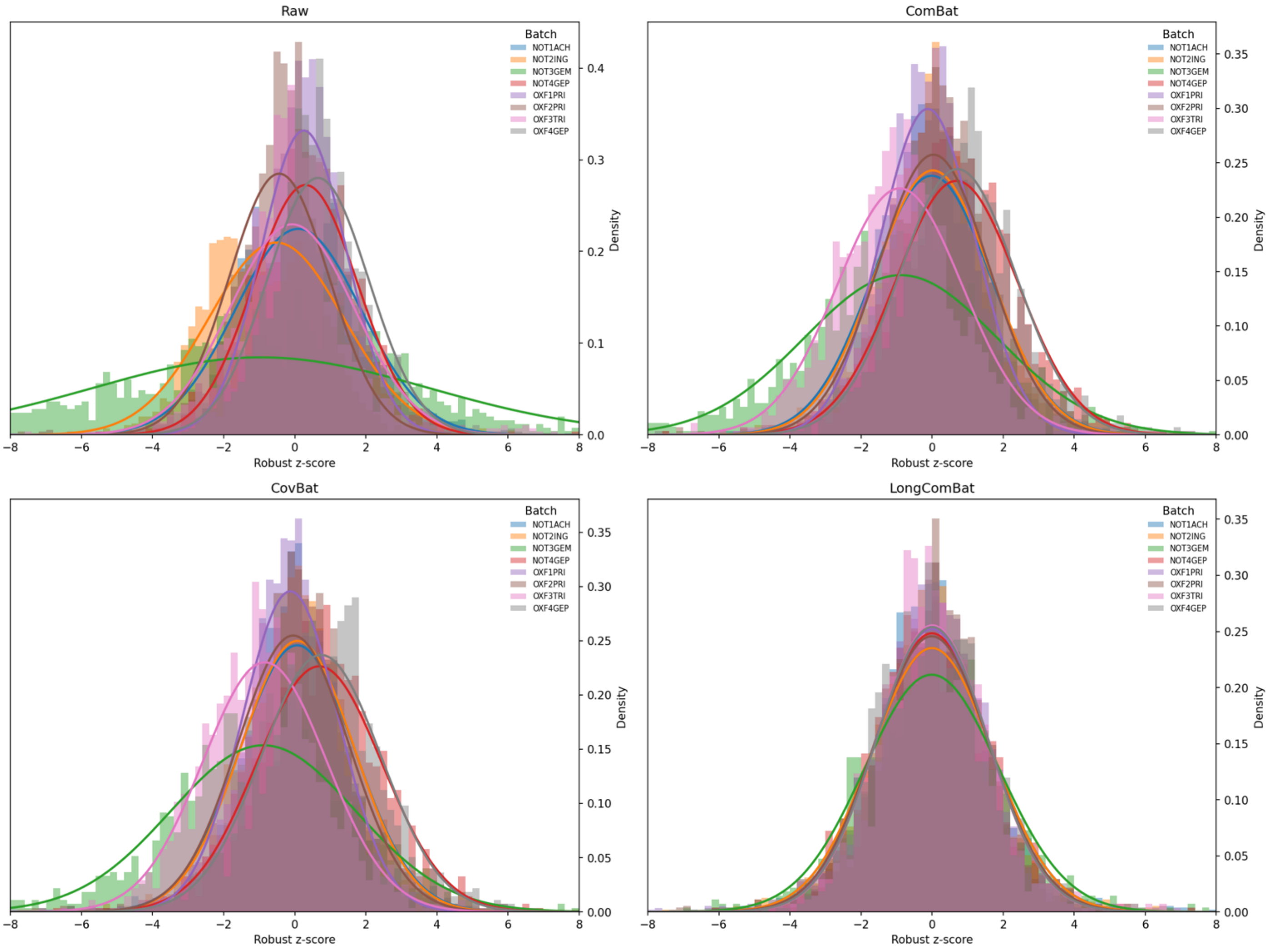


Longitudinal comparison: covariate-adjusted z-score distributions

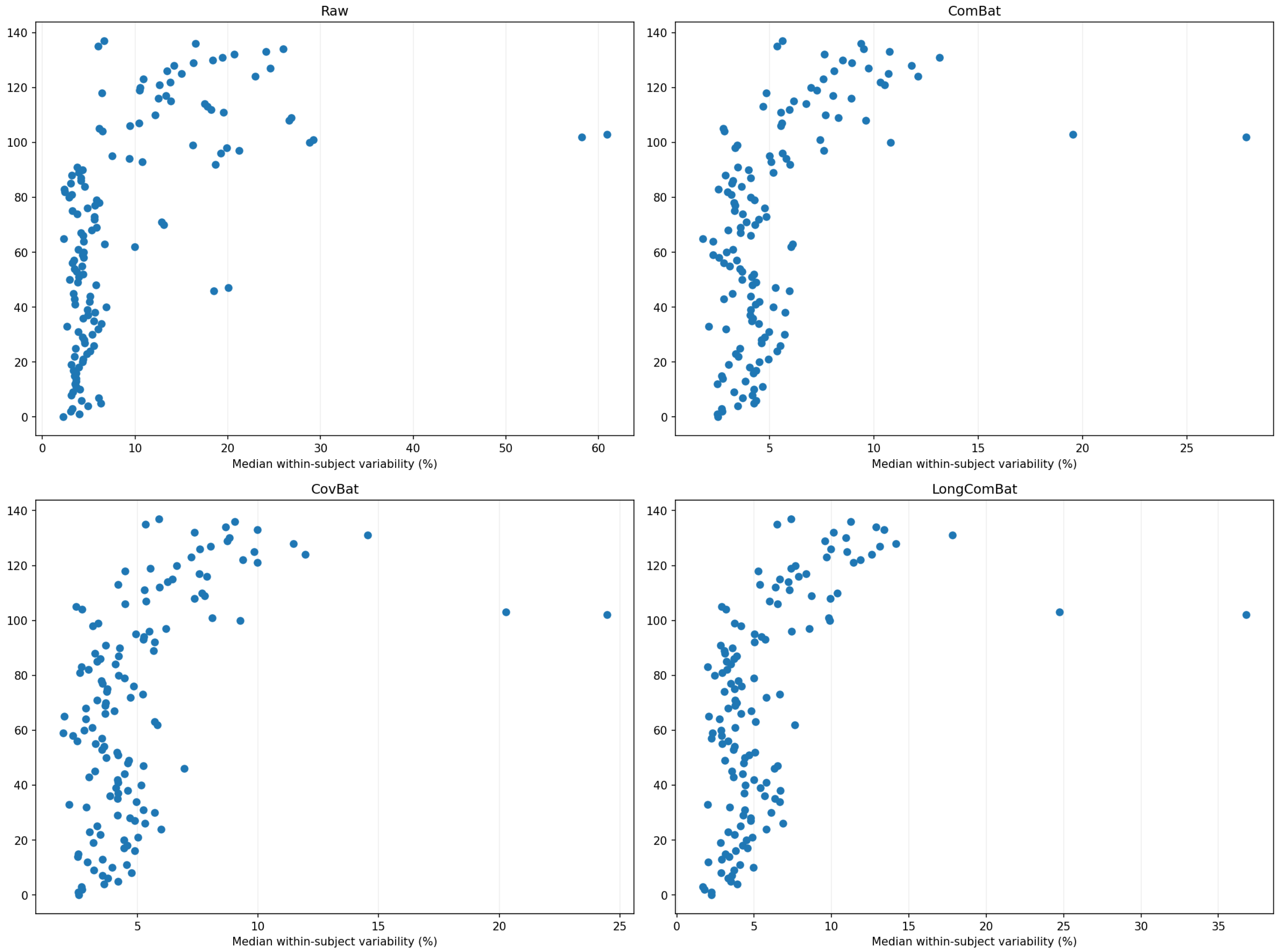


Longitudinal comparison: within-subject variability

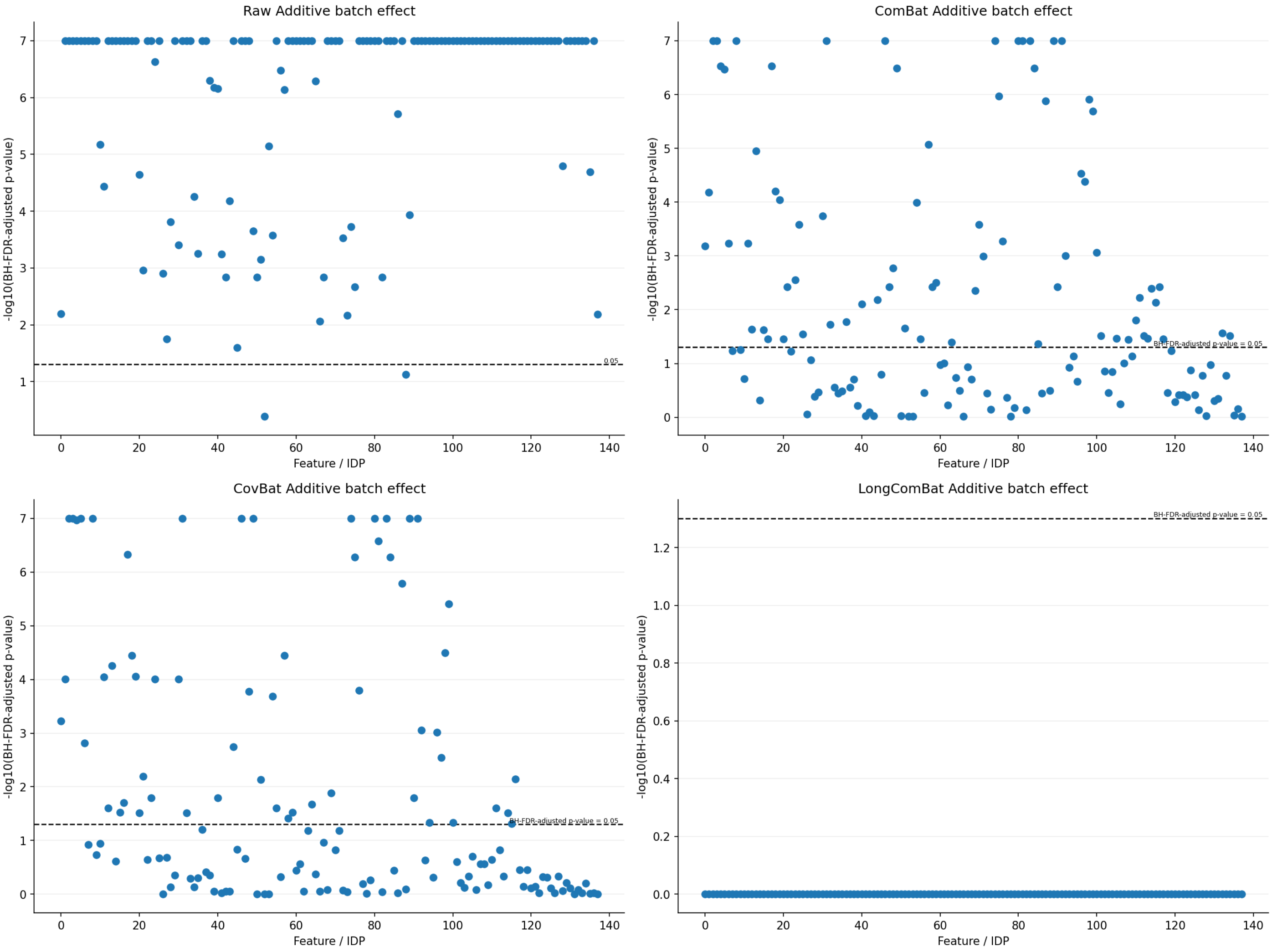


Longitudinal comparison: additive batch effects

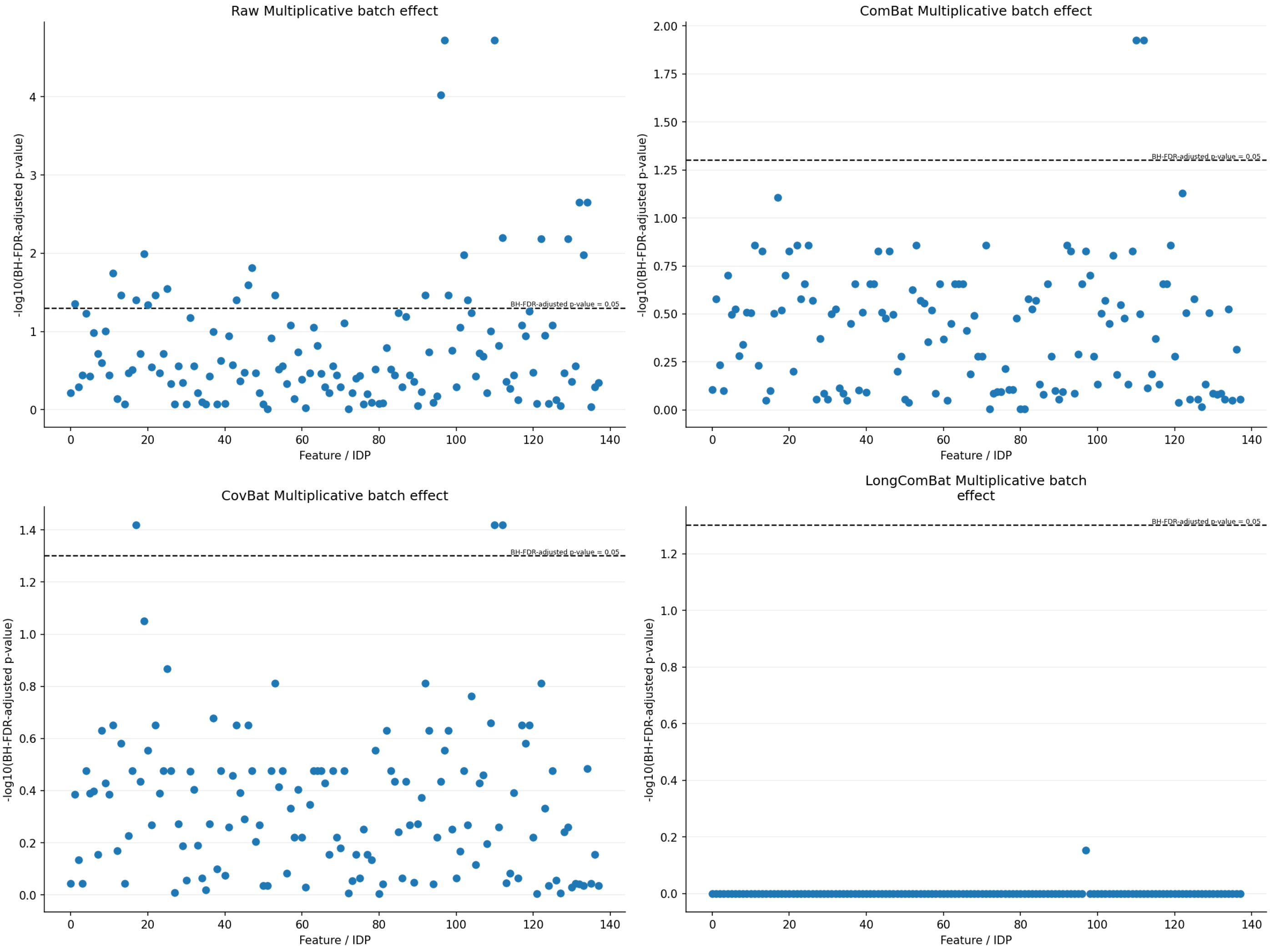


Longitudinal comparison: multiplicative batch effects

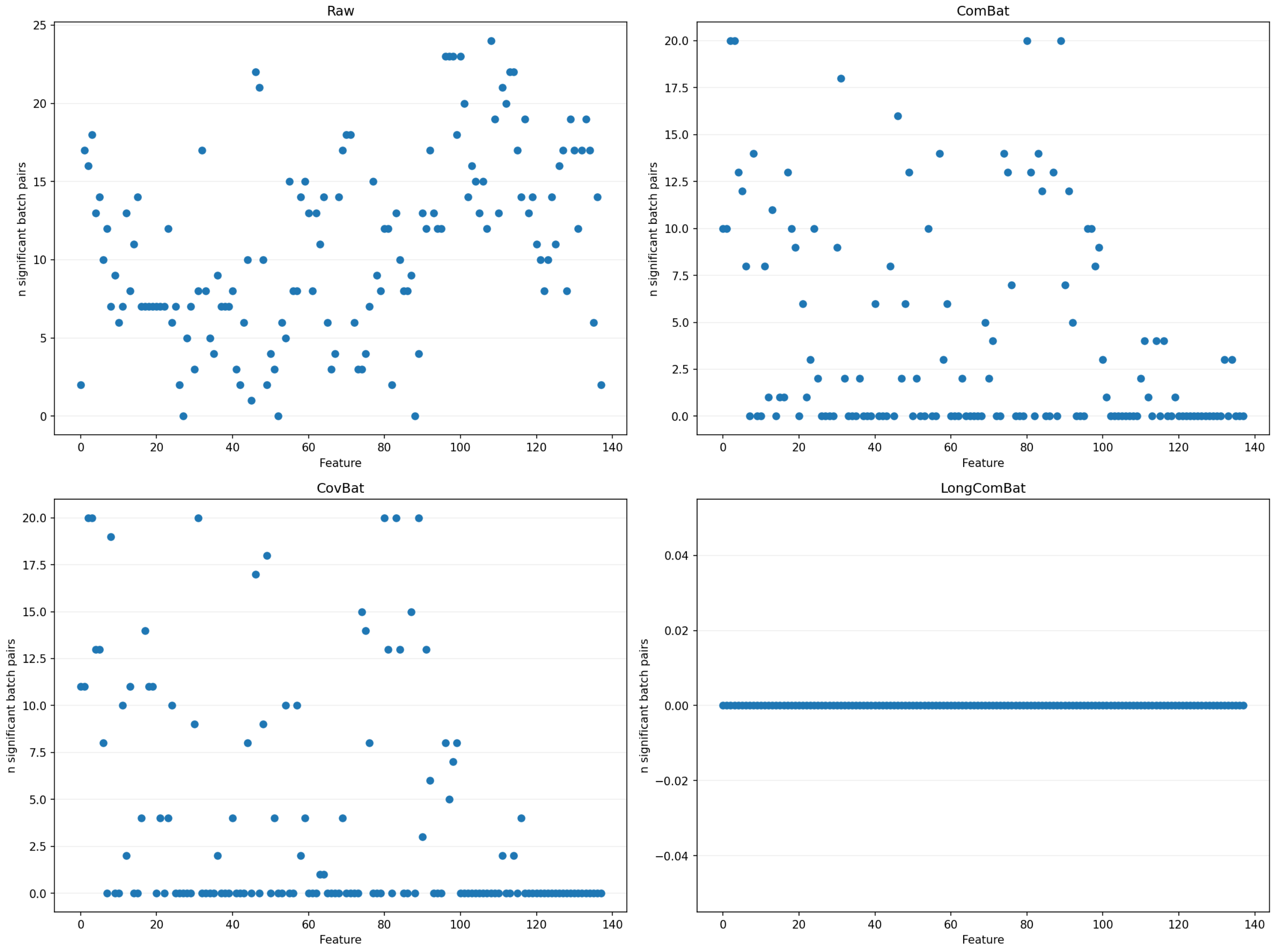


Longitudinal comparison: pairwise batch differences

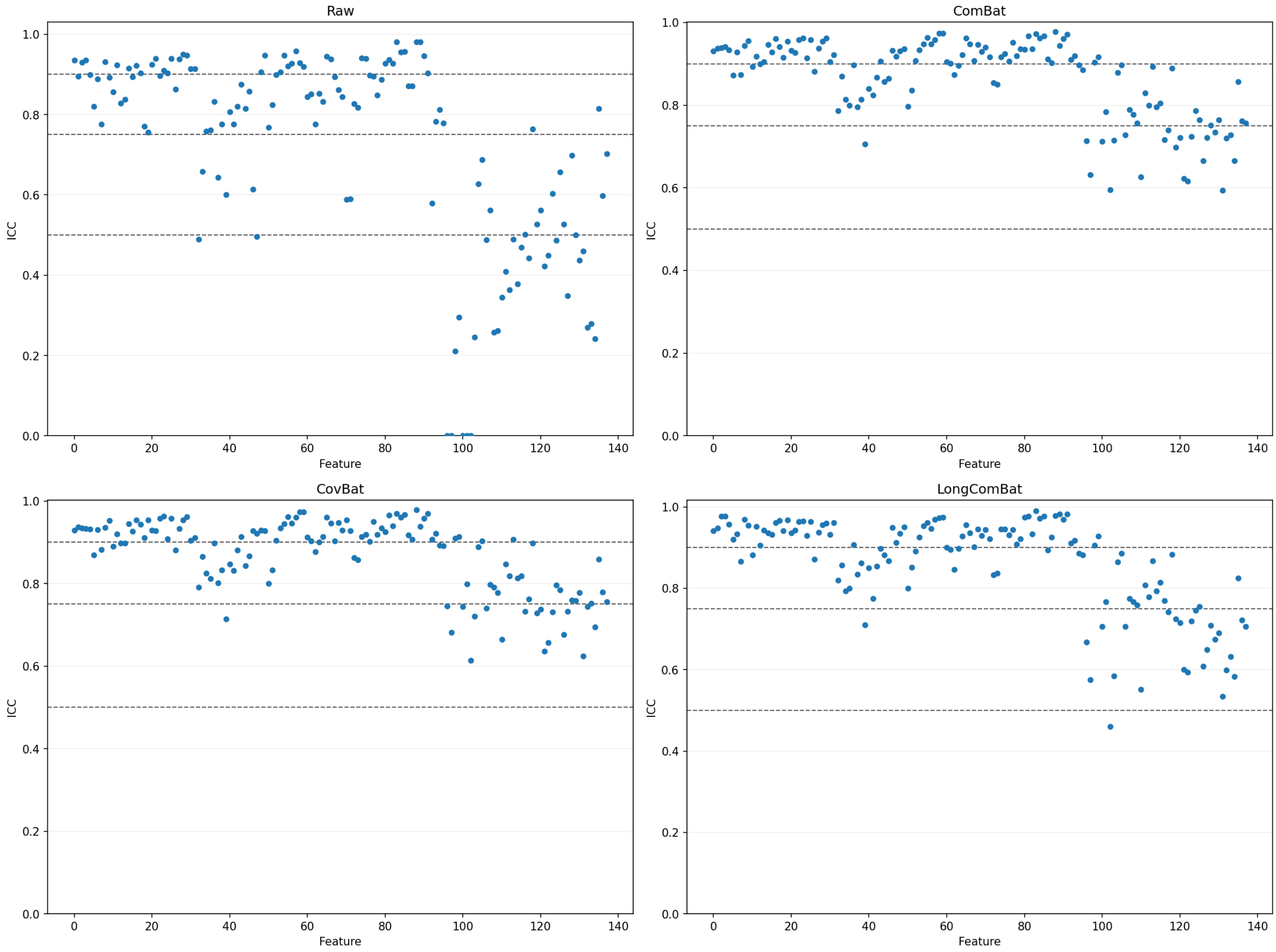


Longitudinal comparison: ICC per feature

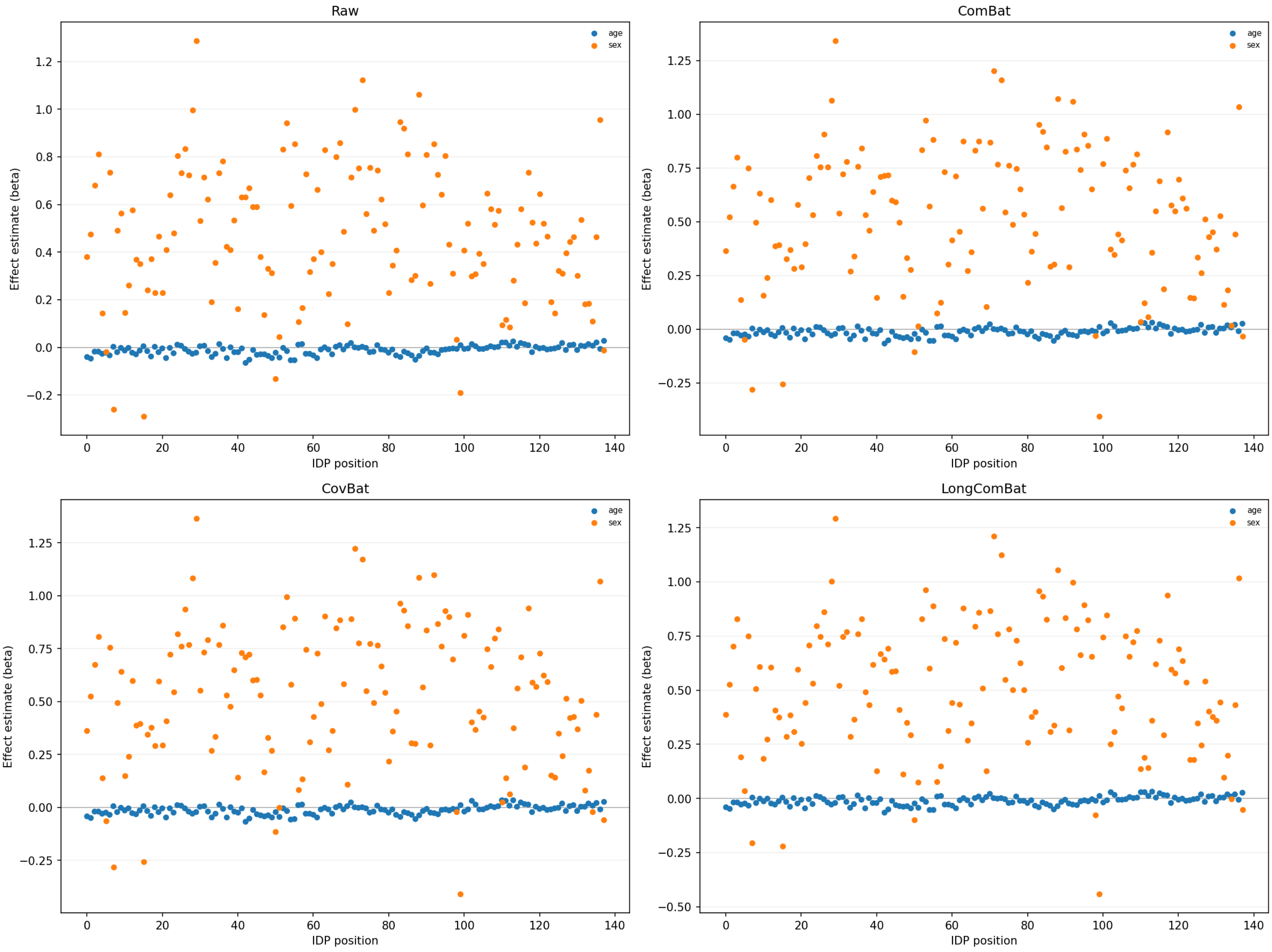


Longitudinal comparison: biological effect preservation

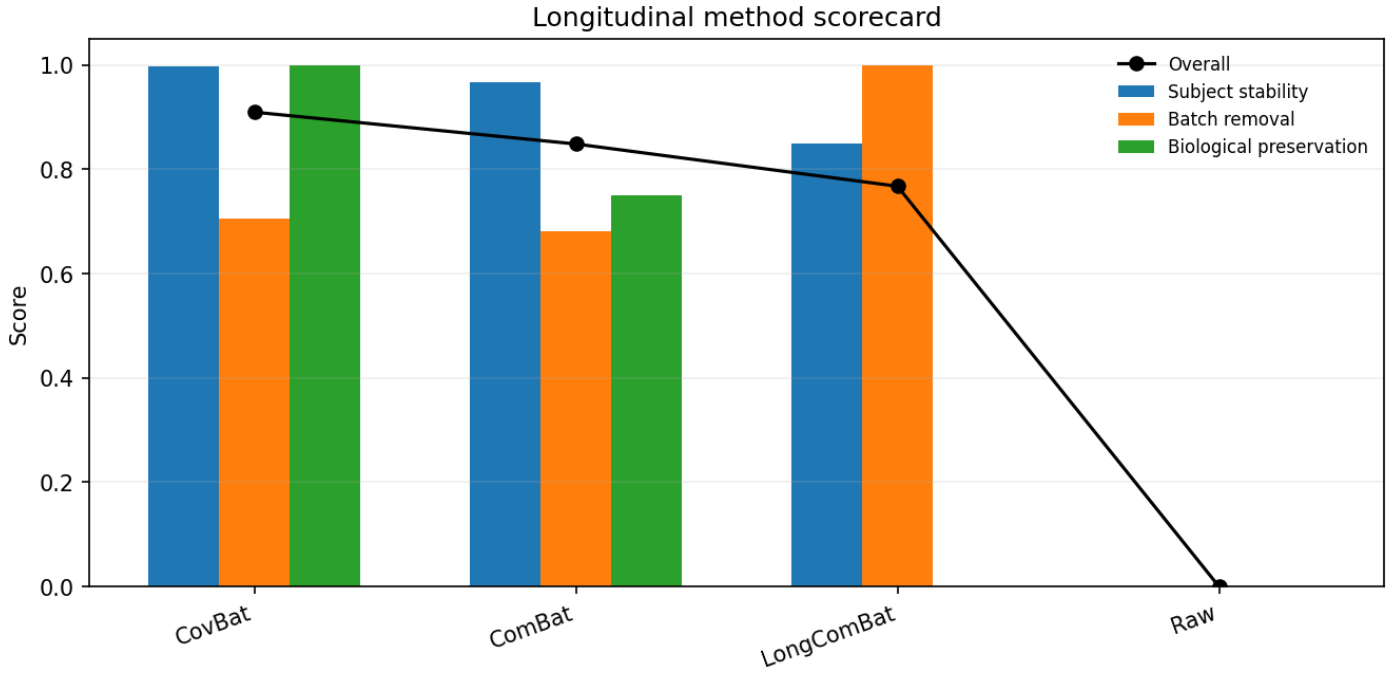


Longitudinal comparison: method scorecard
